# Assessment of AlphaFold structures and optimization methods for virtual screening

**DOI:** 10.1101/2023.01.10.523376

**Authors:** Yanfei Peng, Xia Wu, Liang Lin, Zhiluo Deng, Limin Zhao, Hao Ke

## Abstract

Recent advancements in artificial intelligence such as AlphaFold, have enabled more accurate prediction of protein three-dimensional structure from amino acid sequences. This has attracted significant attention, especially for the application of AlphaFold in drug discovery. However, how to take full advantage of AlphaFold to assist with virtual screening remains elusive. We evaluate the AlphaFold structures of 51 selected targets from the DUD-E database in virtual screening. Our analyses show that the virtual screening performance of about 35% of the AlphaFold structures is equivalent to that of DUD-E structures, and about 25% of the AlphaFold structures yield better results than the DUD-E structures. Remarkably, AlphaFold structures produce slightly better results than the Apo structures. Moreover, we develop a new consensus scoring method based on Z-score standardization and exponential function, which shows improved screening performance compared to traditional scoring methods. By implementing a multi-stage virtual screening process and the new consensus scoring method, we are able to improve the speed of virtual screening by about nine times without compromising the enrichment factor. Overall, our results provide insights into the potential use of AlphaFold in drug discovery and highlight the value of consensus scoring and multi-stage virtual screening.

## Introduction

AlphaFold is deep learning-based program that can predict the three-dimensional structure of proteins from amino acid sequences^1,2^. It is one of the low-cost computing methods to obtain highly accurate protein structures. Currently, the AlphaFold protein structure database contains over 200 million entries, covering the human proteome and the proteome of 47 other important organisms crucial to research and global health. The availability of large numbers of easily accessible and highly accurate protein structures have made AlphaFold a valuable resource for research fields related to protein structure, particularly for half of understudied (dark) human proteins^3^.

Several studies have explored the use of AlphaFold protein structure for drug discovery^4–6^. At present, only a few recent studies assess the performance of AlphaFold protein structure in VS (virtual screening). Wong et al. evaluate the effectiveness of AlphaFold protein structure to predict the binding affinity of 296 E. coli proteins and 218 small molecules with known antibacterial activity, and compared the results to the experimental structure of 12 proteins^7^. Scardino et al. estimate the performance of the structure in the PDB database and AlphaFold structure in virtual screening using four docking software and ECR (Exponential Consensus Ranking, based on ranking) on 16 protein targets, and the PRC (Pose/Ranking Consensus, based on docking pose and ranking)^8^. Zhang et al. examine the efficiency of Holo, Apo and AlphaFold structures in virtual screening for 28 targets with Glide molecular docking method^9^. Alon, et al. show that although the AlphaFold structure of σ2 receptor is very similar to the crystal structure, the score of small molecules with the AlphaFold structure is lower than that of the crystal structure^10^.

Virtual screening based on molecular docking is a widely used method in computer-aided drug design. With scoring function, molecular docking can effectively identify small molecules that interact with the ligand binding pocket of receptor protein. Researchers also provide detailed guidelines for the use of molecular docking based on VS to discover inhibitors and agonists of target proteins^11^, which effectively reduces the cost of drug discovery and accelerate the process. There are many different molecular docking methods widely used in virtual screening, including AutoDock Vina^12,13^, Qvina2^14^, idock^15^, ICM^16^, Glide^17^, Gold^18^, etc. However, molecular docking still has high false positive rate which limits its use in drug discovery. Some recent studies have used the consensus scoring method to integrate the score reported by different docking software to reduce the false positive rate and improve the enrichment factor^19,20^. Researches also have taken into account the docking pose in consensus scoring, which can further improve the performance of the virtual screening^21^. Therefore, further optimization of the consensus scoring method could be of great significance for improving the value of AlphaFold in virtual screening.

With the rapid expansion of the small molecule library for virtual screening, for example, the ZINC database has provided more than 120 million purchasable drug-like compounds^22^, large-scale virtual screening plays an increasingly important role in drug discovery. However, large-scale virtual screening requires a substantial amount of computing resources. To reduce the runtime and save computing resources, the multi-stage screening method has been used in large-scale virtual screening. Gorculla et al. firstly conducted an ultra-large-scale screening with 1.3 billion compounds using a multi-stage screening method. In the first stage, Qvina2 is used for rapid screening with low accuracy. In the second stage, 13 residues of the receptor are considered flexible, and AutoDock Vina and Smina Vinardo are used to re-score the top 3 million compounds from the first stage with a higher accuracy^23,24^.

Protein structure can be categorized into Holo structure and Apo structure based on the presence of small molecules in the binding pocket. Holo structures have small molecules in the binding pocket, while Apo structures do not. Previous studies have shown that the category of the protein structure has a significant impact on virtual screening, and the results of Holo structures are generally better than that of Apo structures ^25,26^.

To comprehensively evaluate the impact of using AlphaFold protein structures, Holo and Apo structures on virtual screening, we select 51 protein targets from the widely used DUD-E database^27^ and more than 400,000 small molecules. A novel consensus scoring function is developed to achieve a lower false discovery rate. In the benchmark, various molecular docking software, scoring functions, and consensus scoring including the novel consensus scoring method are used. Our analyses show that the virtual screening performance of about 35% of the AlphaFold structures is equivalent to that of DUD-E structures, and about 25% of the AlphaFold structures yield better results than the DUD-E structures. Remarkably, for the 23 targets, AlphaFold structures produce slightly better results than the Apo structures. Notably, the new consensus scoring method is superior to the traditional scoring method and can be used to design a multi-stage virtual screening process that improves the virtual screening speed by about 9 times without compromising the enrichment factor EF1%.

## Results

### Virtual screening based on protein structure in DUD-E database and AlphaFold

We selected 51 targets in the DUD-E database (http://dude.docking.org/targets) as our test targets representing eight types of protein targets namely protease, nuclear receiver, kinase, cytochrome P450, ion channel, GPCR, other enzymes and miscellaneous proteins (Fig. 1a).Then, we compared the difference of scores from DUD-E structures and AlphaFold structures. With autodock4 scoring function, we found all the metrics did not show substantial difference (Fig. 1b, Fig. 1c) (*p* value for logAUC between DUD-E and AlphaFold is 0.081). However, with the other scoring functions, DUD-E structures performed better than AlphaFold structures overall in terms of all metrics except for AUC (Fig. 1c, Table 1 and S1-S8). For idock the EF1% was 6.342 ± 8.808 (Data are mean ± s.d.) for DUD-E structure and 3.985 ± 5.191 for AlphaFold structure (*p* value = 0.010). The logAUC based on idock for DUD-E structure were 0.252 ± 0.113 and 0.228 ± 0.088 for AlphaFold structure (*p* value = 0.021). Notably, the virtual screening based on autodock4 scoring function yielded better results on all metrics. The results suggested that the protein structure in DUD-E database was slightly better than AlphaFold predicted structure for VS.

**Table 1.** Virtual screening results of protein structure in DUD-E database and AlphaFold.

| | AUC | logAU<br>C | BEDROC($\alpha=$<br>321.9) | BEDROC($\alpha$<br>=80.5) | BEDROC($\alpha$<br>=20.0) | EF1% | EF5% | EF10<br>% |
| --- | --- | --- | --- | --- | --- | --- | --- | --- |
| AF ad4 | 0.622 | 0.240 | 0.183 | 0.150 | 0.196 | 5.284 | 3.238 | 2.522 |
| AF idock | 0.635 | 0.228 | 0.126 | 0.114 | 0.166 | 3.985 | 2.624 | 2.226 |
| AF rf_score | 0.609 | 0.208 | 0.087 | 0.086 | 0.140 | 2.961 | 2.254 | 1.984 |
| AF vinardo | 0.614 | 0.217 | 0.128 | 0.109 | 0.157 | 3.892 | 2.531 | 2.105 |
| DUD-E ad4 | 0.645 | 0.270 | 0.229 | 0.194 | 0.235 | 7.002 | 3.915 | 2.896 |
| DUD-E<br>idock | 0.647 | 0.252 | 0.204 | 0.156 | 0.199 | 6.342 | 3.235 | 2.574 |
| DUD-E<br>rf_score | 0.625 | 0.228 | 0.118 | 0.116 | 0.172 | 4.040 | 2.865 | 2.336 |
| DUD-E<br>vinardo | 0.639 | 0.245 | 0.172 | 0.152 | 0.198 | 5.436 | 3.282 | 2.509 |

The mean values for 51 targets of AUC, logAUC, BEDROC ( $\alpha=321.9$ , $\alpha=80.5$ , and $\alpha=20.0$ ), EF1%, EF5%, and EF10% were used to evaluate the screening results. AF is the abbreviation of AlphaFold, indicating the results of AlphaFold structures. DUD-E represents the results of structures in DUD-E database. Ad4 means using the autodock4 scoring function, idock means using the idock scoring function, rf\_score means using the rf\_score scoring function in the idock software, and vinardo means using the vinardo scoring function.

**Figure 1.**
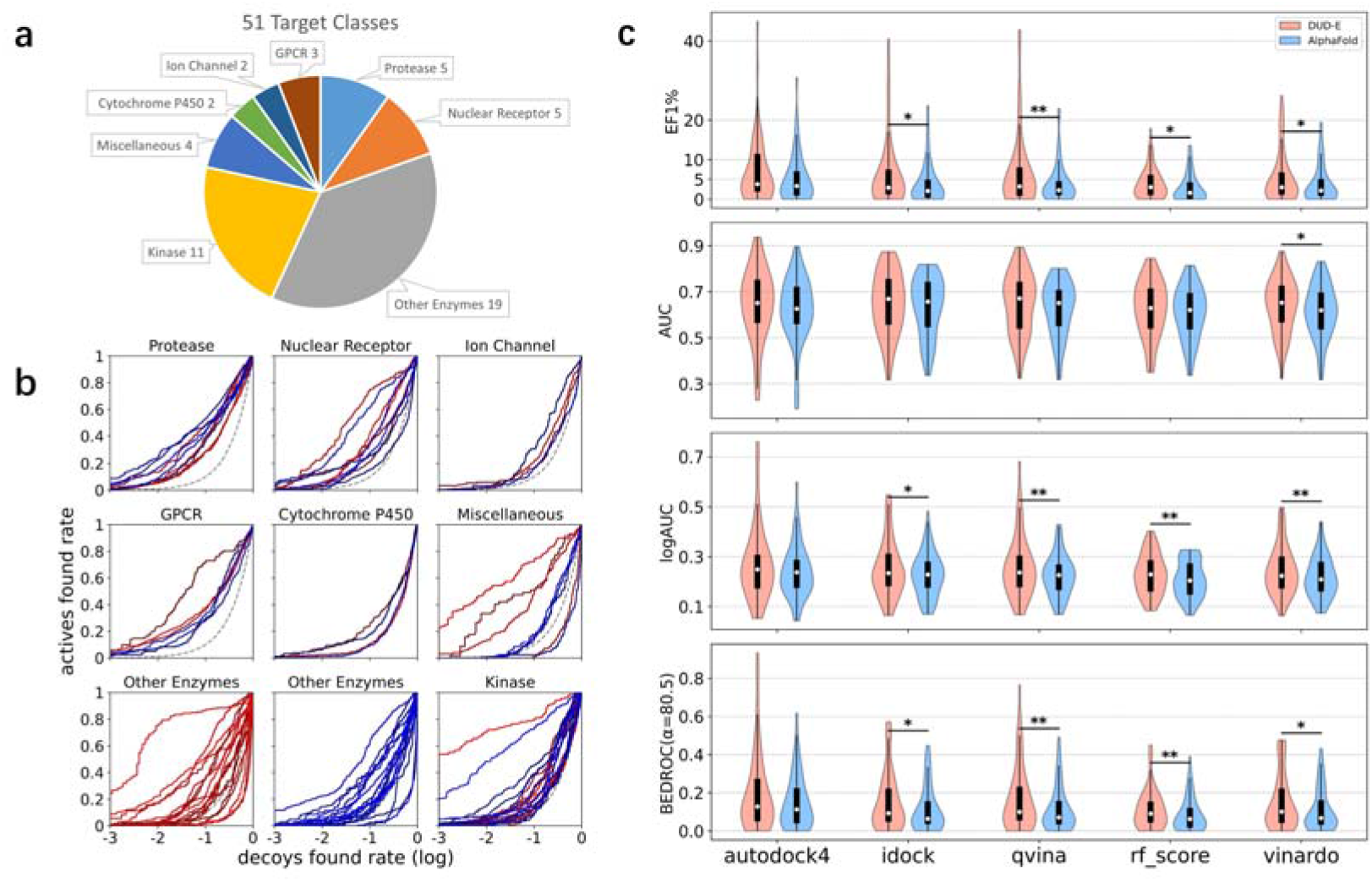
Virtual screening of protein structure in DUD-E database and AlphaFold. (a) Number distribution of protein categories of 51 targets selected from the DUD-E database. (b) Semilogarithmic ROC curve (λ = 0.001) based on the virtual screening results of 51 targets by the AudoDock4 scoring function. The red curves depict the screening results of structures in the DUD-E database, and the blue curves depict the screening results of AlphaFold structures. (c) The violin chart of screening results of autodock4, idock, qvina, rf_score and vinardo scoring function for 51 targets, EF1%, AUC, logAUC, BEDROC (α=80.5) metrics are used to evaluate the screening effect, among which red is the screening result of structure in DUD-E database, and blue is the screening result of AlphaFold structure. According to paired t-test, two groups of data (red and blue) with the same scoring function and the same metric have been marked with *p* values that are statistically different. ** represents p<0.01, and * represents p<0.05.

### Virtual screening results of Holo, AlphaFold and Apo protein structures

Previous studies have shown that the virtual screening result of Holo structures is generally better than that of Apo structures. The protein structures in the DUD-E database are Holo structures. To evaluate whether AlphaFold structures produce better results compared to Apo structures in virtual screening, we tried to collect the Apo structures for the 51 selected targets in DUD-E database. However, only 23 out of these 51 targets have Apo structures available in the PDB database with known binding pocket. We then performed virtual screening with the Apo structures of these 23 targets.

Molecular docking is a technique that involves fitting small molecules into the binding pocket of a protein structure and allows for the identification of appropriate conformations. Therefore, the binding pocket is essential for virtual screening using molecular docking. When analyzing the protein structures, we found that HIVRT had clear binding pocket in its Holo and Apo structures encompassing the eutectic small molecules. However, no apparent binding pocket was found in its AlphaFold structure (Fig. 2a), HIVRT was not included in the 23 targets mentioned above. On the other hand, AKT2, has clear binding pockets in its Holo and AlphaFold structures but not in the Apo structures (Fig. 2b). Therefore, AKT2 was also excluded from the selected targets.

**Figure 2.**
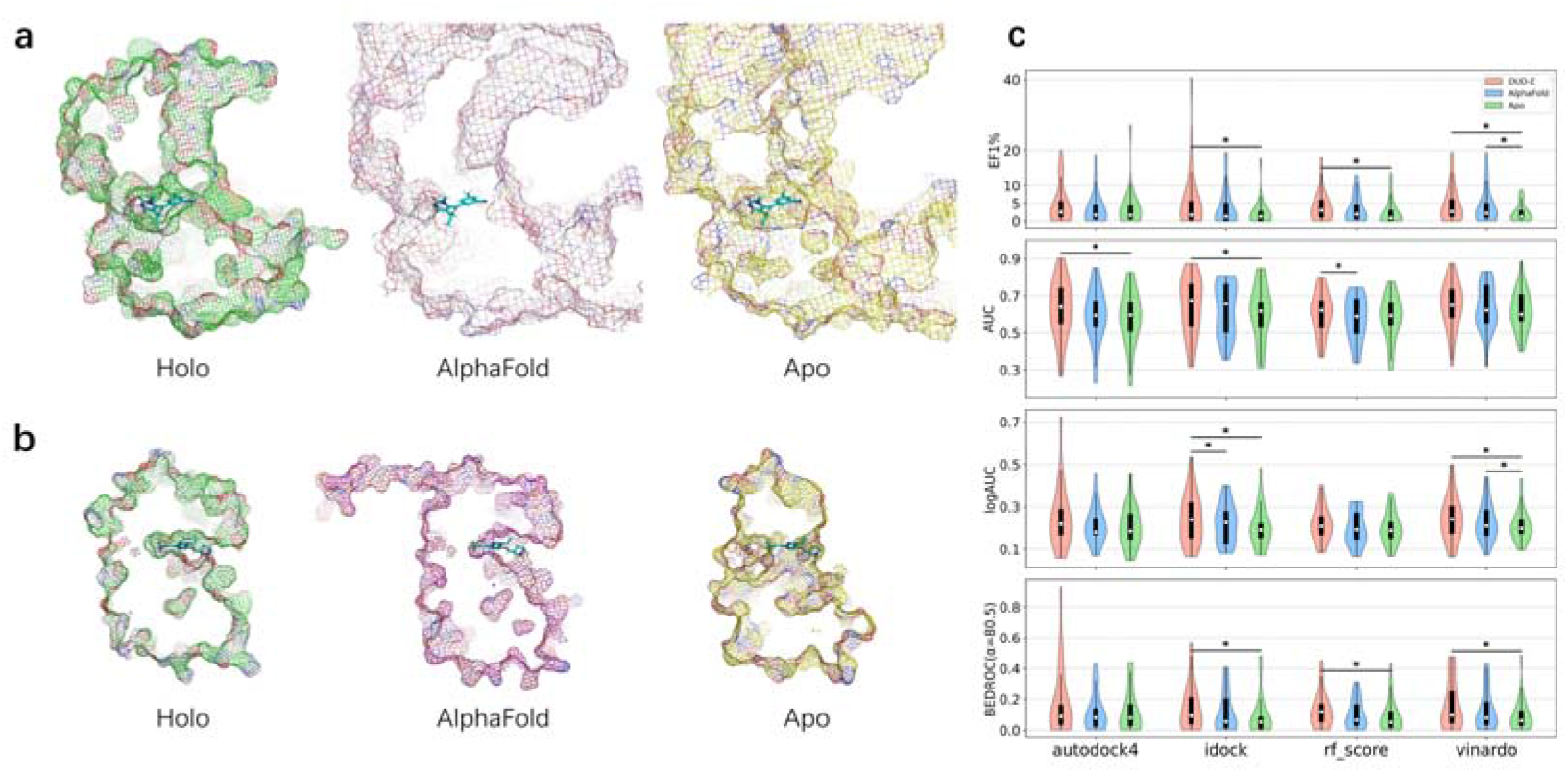
Binding pocket structure and virtual screening results of Holo, AlphaFold and Apo protein structures. (a-b) Water molecules have been removed from protein structures. AlphaFold structure and Apo structure (obtained from PDB database) are aligned with Holo structure (obtained from DUD-E database), displayed in grid form, and cut the same section. Green is Holo structure, purple is AlphaFold structure, and yellow is Apo structure. PyMOL is used for operation and drawing. HIVRT (a) and AKT2 (b) protein structures of Holo, AhphaFold and Apo three-dimensional grid diagram sections were displayed. (c) The violin chart of screening results of autodock4, idock, qvina, rf_score and vinardo scoring function for 23 targets, EF1%, AUC, logAUC, BEDROC (α=80.5) metrics are used to evaluate the screening effect. Among them, red group represent the screening result of protein structure (Holo) in DUD-E database, blue group represent screening result of AlphaFold structure, and green group represent screening result of Apo structure. Three groups of data (red, blue and green) with the same scoring function and the same metric are all marked according to paired t-test with *p* values that are statistically different. ** represents p<0.01, and * represents p<0.05.

The AUC values between Holo and Apo groups were significantly different when using scoring function autodock4 (Holo =0.628 ± 0.154, Apo =0.580 ± 0.142, *p* value = 0.040) and idock (Holo = 0.641 ± 0.152, Apo = 0.603 ± 0.144, *p* value = 0.043). With the vinardo function, AlphaFold structures were significantly better than Apo structures in terms of EF1% (AlphaFold 4.183 ± 4.919, Apo 2.311 ± 2.455, *p* value = 0.032) and logAUC (AlphaFold 0.230 ± 0.087, Apo 0.210 ± 0.074, *p* value = 0.042). It showed that the protein structures in DUD-E database were better than Apo structure in virtual screening. The protein structures in DUD-E database were somewhat better than AlphaFold structures for virtual screening, and AlphaFold structures were slight better than Apo structures in virtual screening.

It is worth noting that the difference between DUD-E and AlphaFold groups in Figure 2c was relatively smaller than the difference in Figure 1c. It was because the number of protein targets tested in Figure 2c was less. The virtual screening of protein structure in DUD-E database (the mean of logAUC for idock is 0.244) was better than the mean value of AlphaFold structure (the mean of logAUC for idock is 0.220) in different scoring functions and different metrics. And the virtual screening of AlphaFold structures was better than the overall scores of Apo structures (the mean of logAUC for idock is 0.202), while different scoring functions had a significant impact on the results of virtual screening (Table 2 and S9-S12).

**Table 2.** Virtual screening results of Holo, AlphaFold, Apo protein structure.

| | AUC | logAU<br>C | BEDROC($\alpha=$<br>321.9) | BEDROC($\alpha$<br>=80.5) | BEDROC($\alpha$<br>=20.0) | EF1% | EF5% | EF10<br>% |
| --- | --- | --- | --- | --- | --- | --- | --- | --- |
| AF ad4 | 0.587 | 0.212 | 0.141 | 0.121 | 0.167 | 3.638 | 2.497 | 2.100 |
| AF idock | 0.625 | 0.220 | 0.129 | 0.115 | 0.161 | 3.700 | 2.291 | 2.020 |
| AF rf_score | 0.576 | 0.200 | 0.103 | 0.101 | 0.148 | 3.330 | 2.439 | 1.951 |
| AF vinardo | 0.633 | 0.230 | 0.152 | 0.126 | 0.176 | 4.183 | 2.622 | 2.306 |
| Apo ad4 | 0.580 | 0.206 | 0.124 | 0.122 | 0.164 | 3.593 | 2.396 | 2.029 |
| Apo idock | 0.603 | 0.202 | 0.093 | 0.089 | 0.139 | 2.308 | 2.104 | 1.797 |
| Apo<br>rf_score | 0.581 | 0.198 | 0.099 | 0.092 | 0.146 | 2.440 | 2.249 | 1.955 |
| Apo<br>vinardo | 0.617 | 0.210 | 0.074 | 0.095 | 0.157 | 2.311 | 2.466 | 2.035 |
| DUD-E ad4 | 0.628 | 0.248 | 0.191 | 0.172 | 0.212 | 4.757 | 3.267 | 2.505 |
| DUD-E<br>idock | 0.641 | 0.244 | 0.179 | 0.147 | 0.196 | 5.455 | 3.025 | 2.436 |
| DUD-E<br>rf_score | 0.603 | 0.218 | 0.134 | 0.127 | 0.171 | 4.016 | 2.804 | 2.164 |
| DUD-E<br>vinardo | 0.645 | 0.245 | 0.171 | 0.160 | 0.207 | 4.667 | 3.333 | 2.583 |

The mean values for 51 targets of AUC, logAUC, BEDROC ( $\alpha=321.9$ , $\alpha=80.5$ , and $\alpha=20.0$ ), EF1%, EF5%, and EF10% were used to evaluate the screening results. AF is the abbreviation of AlphaFold, indicating the results of AlphaFold structures. DUD-E represents the results of structures in DUD-E database. Ad4 means using the autodock4 scoring function, idock means using the idock scoring function, rf\_score means using the rf\_score scoring function in the idock software, and vinardo means using the vinardo scoring function.

### Evaluation of consensus scoring method based on score or rank

Previous studies have shown that consensus scoring based on multiple scoring functions^19,20,28,29^ can take advantages of each scoring function to achieve better screening results. However, different scoring functions may need to be standardized before they can be integrated. Common standardization methods include ranking, AASS, Average of auto scaled scores, Z-score scaling.

In this study, we selected four appropriate scoring functions (autodock4, idock, rf_score, and vinardo) and combined them using 12 different consensus scoring methods (3 standardization methods × 4 consensus calculation methods). We tested these methods and RBV (rank by vote) method on 51 previously selected targets, with the results of the four single scoring functions as controls.

Different scoring functions may produce similar results, which will not help improving the consensus outcome. So, we tested the correlation between each one another among the five scoring functions including autodock4, idock, qvina, rf_score and vinardo. The square of pearson correlation coefficient R^2^ between qvina and idock was greater than 0.9 (logAUC = 0.937, BEDROC(α=80.5) = 0.930) and the fitting line (logAUC: *y* = 0.909*x* + 0.018, BEDROC(α=80.5): *y* = 1.058*x* + 0.002) was close to y=x (Fig. 3a), while the square of pearson correlation coefficient R^2^ between other pairs were less than 0.75. These results indicated that qvina and idock had a high similarity. In fact, qvina and idock were both developed based on AutoDock Vina. Therefore, we finally selected autodock4, idock, rf_score and vinardo scoring functions for consensus scoring.

**Figure 3.**
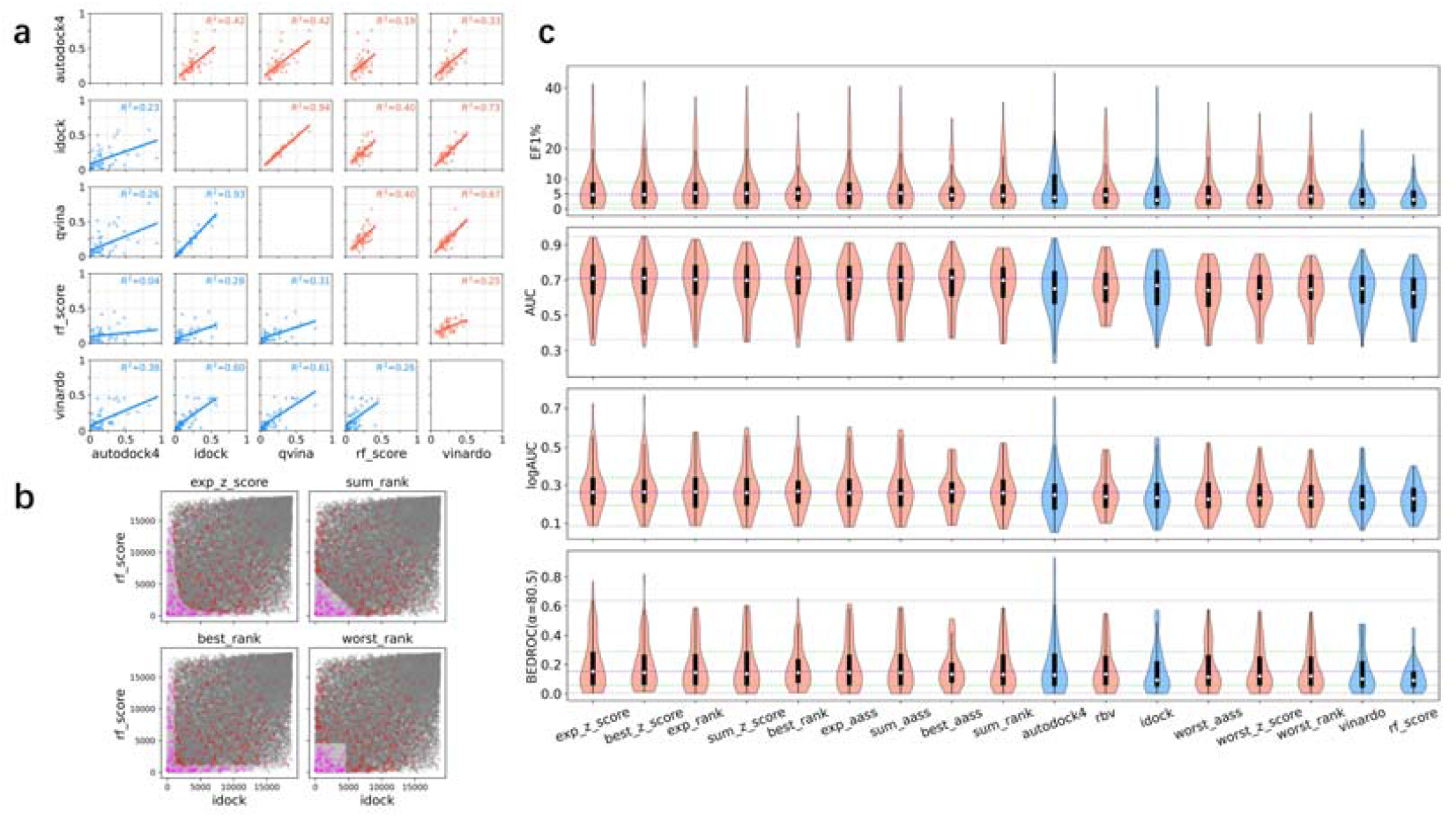
Evaluation of consensus scoring method based on score or rank. (a) Scatter plots and fitting lines are drawn between different scoring functions according to the results of autodock4, idock, qvina, rf_score and vinardo for 51 targets. The red is logAUC, and the blue is BEDROC (α=80.5). (b) Scatter plots are drawn according to the results of four consensus scoring methods: exp_z_score, sum_rank, best_rank, and worst_rank (all with idock and rf_score as inputs) for the representative BACE1 target in DUD-E database. The abscissa is the rank of small molecules in idock, and the ordinate is the rank of small molecules in rf_score, in which that gray and silver are decoys, red and purple are actives, and silver and purple are the top 10% of the ranking obtained by the corresponding consensus scoring method. (c) The violin chart of screening results of 4 single scoring functions (blue) and 13 consensus scoring methods (red, with autodock4, idock, rf_score, vinardo as the input) for 51 targets, EF1%, AUC, logAUC, BEDROC (α=80.5) metrics are used to evaluate the screening effect. The x-axis scoring function is sorted by the mean value of logAUC, with marked exp_z_score corresponding to the upper limit and lower limit of the data (gray dotted line), quartile (green dotted line), and median (blue dotted line).

To show four different calculation methods of consensus scoring, we compared the distribution score of idock, rf_score and consensus evaluation methods on the representative BACE1 protein (Fig. 3b). From the distribution of active compounds and decoy compounds, the top small molecules were more likely to be active compounds (for idock the EF10% was 2.66, while for rf_score the EF10% was 2.87), indicating that idock and rf_score had certain ability to enrich active compounds. Moreover, we found that the best exp_z_score, consensus scoring of exponential function based on Z-score scaling, was generally better than that of autodock4, idock and rf_score and vinardo scoring functions. The exp_z_score was also superior to most other consensus scoring methods (Fig. 3c).

The exp_rank (ECR) was also a consensus scoring method based on ranking exponential function. Compared the ECR and exp_z_score groups by paired *t*-test, exp_z_score showed better than ECR method (for example, logAUC for exp_z_score groupwas 0.290 while ECR group was 0.284), except for the AUC (exp_z_score group is 0.686 while ECR group was 0.689). Notably, BEDROC(α=20.0) and EF10% of exp_z_score (0.253 and 3.202) were higher than that of ECR (0.244 and 3.104) with significant difference (*p* value is 0.028 and 0.028) (Table 3 and S13-S15).

**Table 3.** Evaluation consensus scoring method based on score or rank.

| | AUC | logAUC | BEDROC($\alpha=321.9$) | BEDROC($\alpha=80.5$) | BEDROC($\alpha=20.0$) | EF1% | EF5% | EF10% |
| --- | --- | --- | --- | --- | --- | --- | --- | --- |
| exp_z_score | 0.686 | 0.290 | 0.248 | 0.204 | 0.253 | 7.736 | 4.226 | 3.202 |
| best_z_score | 0.687 | 0.285 | 0.222 | 0.193 | 0.246 | 7.038 | 4.162 | 3.101 |
| exp_rank | 0.689 | 0.284 | 0.247 | 0.193 | 0.244 | 7.423 | 4.047 | 3.104 |
| sum_z_score | 0.676 | 0.282 | 0.250 | 0.197 | 0.243 | 7.697 | 4.059 | 3.068 |
| best_rank | 0.688 | 0.280 | 0.187 | 0.180 | 0.243 | 6.533 | 4.138 | 3.118 |
| exp_aass | 0.671 | 0.279 | 0.248 | 0.195 | 0.240 | 7.617 | 3.982 | 3.021 |
| sum_aass | 0.669 | 0.278 | 0.248 | 0.194 | 0.238 | 7.506 | 3.955 | 2.971 |
| best_aass | 0.678 | 0.272 | 0.192 | 0.173 | 0.230 | 6.356 | 3.893 | 2.998 |
| sum_rank | 0.673 | 0.272 | 0.234 | 0.179 | 0.227 | 6.782 | 3.722 | 2.901 |
| ad4 | 0.645 | 0.270 | 0.229 | 0.194 | 0.235 | 7.002 | 3.915 | 2.896 |
| rbv | 0.659 | 0.260 | 0.211 | 0.171 | 0.223 | 6.735 | 3.636 | 3.003 |
| idock | 0.647 | 0.252 | 0.204 | 0.156 | 0.199 | 6.342 | 3.235 | 2.574 |
| worst_aass | 0.633 | 0.250 | 0.204 | 0.164 | 0.206 | 6.187 | 3.430 | 2.608 |
| worst_z_score | 0.634 | 0.249 | 0.223 | 0.166 | 0.204 | 6.366 | 3.363 | 2.534 |
| worst_rank | 0.635 | 0.249 | 0.223 | 0.164 | 0.202 | 6.258 | 3.297 | 2.507 |
| vinardo | 0.639 | 0.245 | 0.172 | 0.152 | 0.198 | 5.436 | 3.282 | 2.509 |
| rf_score | 0.625 | 0.228 | 0.118 | 0.116 | 0.172 | 4.040 | 2.865 | 2.336 |
| *p* value | 0.054 | 0.075 | 0.961 | 0.102 | **0.028** | 0.399 | 0.068 | **0.028** |

The mean values for 51 targets of AUC, logAUC, BEDROC ( $\alpha=321.9$ , $\alpha=80.5$ , and $\alpha=20.0$ ), EF1%, EF5%, and EF10% were used to evaluate the screening results. Ad4 means to use the autodock4 scoring function, idock means to use the idock scoring function, rf\_score means to use rf\_score scoring function in idock software, and vinardo means to use vinardo scoring function. In the last line, the *p* value is the result of paired t-test on exp\_z\_score index data and exp\_rank (ECR) index data. The data ( $p<0.05$ ) has been bold. The remaining data are the screening results of consensus scoring methods.

### Re-evaluation of Holo, AlphaFold and Apo protein structures using consensus scoring method

We also evaluated the virtual screening results of Holo, AlphaFold and Apo structures with four representative consensus scoring methods. With the consensus scoring functions, DUD-E structure generated better results compared to AlphaFold structure (such as for exp_z_score, the logAUC for DUD-E was 0.290 ± 0.133, while the logAUC for AlphaFold was 0.256 ± 0.096, and the *p* value for logAUC between DUD-E and AlphaFold was 0.014) (Fig. 4a). The mean values of main data groups had substantial differences, while the direct mean values of a few data groups did not have significant differences. Remarkably, AlphaFold structure was slightly better than Apo structure in virtual screening with the consensus scoring methods (such as for exp_z_score, the EF1% for AlphaFold was 5.161 ± 5.284 while the EF1% for Apo was 3.223 ± 5.321, and the *p* value for EF1% between AlphaFold and Apo is 0.050) (Fig. 4b).

**Figure 4.**
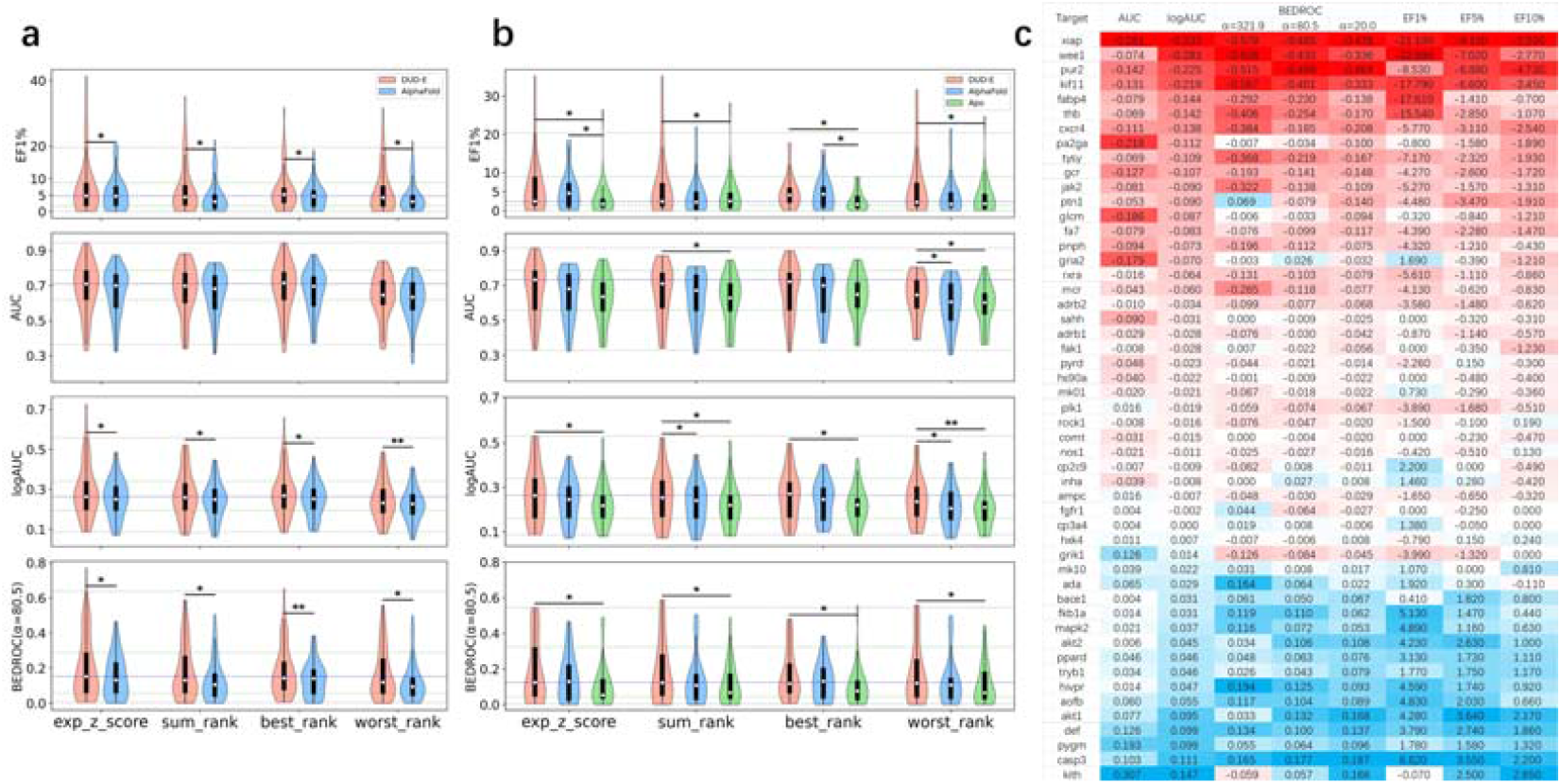
Re-evaluation of Holo, AlphaFold and Apo protein structures using consensus scoring method. (a-b) The results of four consensus scoring methods (all with autodock4, idock, rf_score, vinardo as input): exp_z_score, sum_rank, best_rank and worst_rank. And EF1%, AUC, logAUC, BEDROC (α=80.5) metrics are used to evaluate the screening effect. (a) The violin chart shows the screening results of DUD-E structure (red) and AlphaFold structure (blue) of 51 targets. (b) The violin chart shows the screening results of DUD-E structure (red), AlphaFold structure (blue) and Apo structure (green) of 23 targets. (c) The results of exp_z_score for 51 targets were used to count the difference between structures in DUD-E database and AlphaFold structure of AUC, logAUC, BEDROC (α=321.9, α=80.5, and α=20.0) and EF (1%, 5%, and 10%). Negative number (red) indicates that the AlphaFold structure is worse than the structure in DUD-E database, and positive number (blue) indicates that the AlphaFold structure is better than the structure in DUD-E database. The data in the table is sorted by the difference of logAUC without decreasing. There are 20 targets (about 39% of protein targets) whose difference of logAUC is less than −0.03, 18 (about 35%) targets whose difference of logAUC is greater than −0.03, and less than or equal to 0.03, 13 targets (about 25%) with a difference of logAUC greater than 0.03.

In addition, for each of the 51 previously selected targets, we also made a difference for each target between AlphaFold and Holo results, namely AlphaFold – DUD-E (Fig. 4c). If the difference of defined logAUC was less than −0.03, it indicates that the virtual screening result of AlphaFold structures was worse than that of DUD-E structures. When the difference of defined logAUC was greater than or equal to −0.03 and less than or equal to 0.03, it indicated that the virtual screening result of AlphaFold structures was equivalent to that of DUD-E structures. When the difference of defined logAUC was greater than 0.03, it indicated that the virtual screening result of AlphaFold structures was better than that of DUD-E structures. For 20 targets (about 39%) that AlphaFold structures were worse than DUD-E structures, for 18 targets (about 35%) that AlphaFold structures were equivalent to DUD-E structures, and for 13 targets (about 25%) that AlphaFold structures were better than DUD-E structures.

We also studied the 51 Holo structures and AlphaFold structures for exp_z_score by paired t-test. The mean values of AlphaFold minus DUD-E of all metrics were negative value, and there was a significant difference between the mean values of AlphaFold and DUD-E of most metrics (Table 4). These similar results were also observed in the situation of that of Apo minus AlphaFold (Table 5 and S16-S18). With consensus scoring of virtual screening, Holo structures from DUD-E database were better than AlphaFold structure, and Holo structures and AlphaFold structures were better than Apo structures.

**Table 4.**
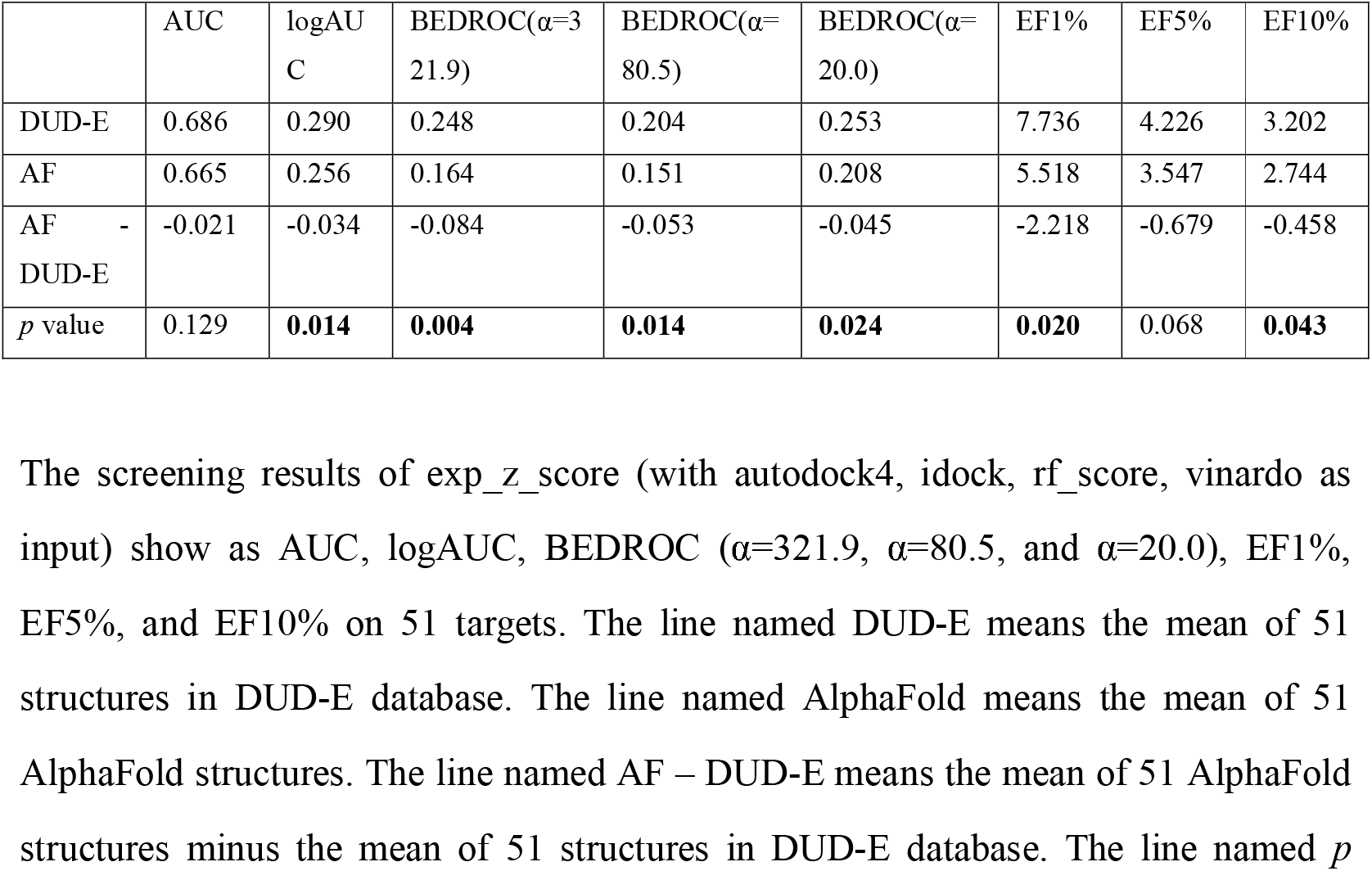
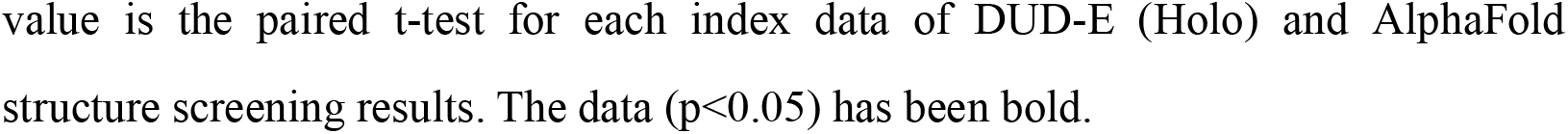
Evaluation of Holo, AlphaFold protein structure using consensus scoring methods.

| | AUC | logAUC | BEDROC($\alpha=321.9$) | BEDROC($\alpha=80.5$) | BEDROC($\alpha=20.0$) | EF1% | EF5% | EF10% |
| --- | --- | --- | --- | --- | --- | --- | --- | --- |
| DUD-E | 0.686 | 0.290 | 0.248 | 0.204 | 0.253 | 7.736 | 4.226 | 3.202 |
| AF | 0.665 | 0.256 | 0.164 | 0.151 | 0.208 | 5.518 | 3.547 | 2.744 |
| AF - DUD-E | -0.021 | -0.034 | -0.084 | -0.053 | -0.045 | -2.218 | -0.679 | -0.458 |
| *p* value | 0.129 | **0.014** | **0.004** | **0.014** | **0.024** | **0.020** | 0.068 | **0.043** |

The screening results of exp\_z\_score (with autodock4, idock, rf\_score, vinardo as input) show as AUC, logAUC, BEDROC ( $\alpha=321.9$ , $\alpha=80.5$ , and $\alpha=20.0$ ), EF1%, EF5%, and EF10% on 51 targets. The line named DUD-E means the mean of 51 structures in DUD-E database. The line named AlphaFold means the mean of 51 AlphaFold structures. The line named AF – DUD-E means the mean of 51 AlphaFold structures minus the mean of 51 structures in DUD-E database. The line named *p*
value is the paired t-test for each index data of DUD-E (Holo) and AlphaFold structure screening results. The data ( $p < 0.05$ ) has been bold.

**Table 5.** Evaluate the protein structure of Holo, AlphaFold and Apo using consensus scoring methods.

| | AUC | logAUC | BEDROC($\alpha=321.9$) | BEDROC($\alpha=80.5$) | BEDROC($\alpha=20.0$) | EF1% | EF5% | EF10% |
| --- | --- | --- | --- | --- | --- | --- | --- | --- |
| DUD-E | 0.671 | 0.271 | 0.218 | 0.189 | 0.237 | 6.295 | 3.644 | 2.918 |
| AF | 0.641 | 0.241 | 0.169 | 0.147 | 0.197 | 5.161 | 3.128 | 2.473 |
| Apo | 0.627 | 0.223 | 0.103 | 0.110 | 0.170 | 3.223 | 2.624 | 2.223 |
| AF - DUD-E | -0.030 | -0.030 | -0.049 | -0.042 | -0.040 | -1.134 | -0.516 | -0.445 |
| AF - Apo | 0.014 | 0.018 | 0.066 | 0.037 | 0.026 | 1.938 | 0.504 | 0.249 |
| Apo - DUD-E | -0.044 | -0.049 | -0.115 | -0.079 | -0.067 | -3.072 | -1.020 | -0.694 |
| AF DUD-E $p$ value | 0.082 | 0.060 | 0.192 | 0.145 | 0.118 | 0.294 | 0.223 | 0.128 |
| AF Apo $p$ value | 0.398 | 0.243 | 0.119 | 0.136 | 0.233 | **0.050** | 0.251 | 0.364 |
| Apo DUD-E $p$ value | 0.052 | **0.020** | **0.021** | **0.019** | **0.029** | **0.011** | 0.064 | 0.055 |

The screening results of exp\_z\_score (with autodock4, idock, rf\_score, vinardo as input) show as AUC, logAUC, BEDROC ( $\alpha=321.9$ , $\alpha=80.5$ , and $\alpha=20.0$ ), EF1%, EF5%, and EF10% on 23 targets. The results of structures in DUD-E database, AlphaFold (AF) structures, Apo structures, AlphaFold minus DUD-E (AF - DUD-E), AlphaFold minus Apo (AF - Apo), and Apo minus DUD-E (Apo - DUD-E) are displayed line by line. The $p$ value is the paired t-test for each index data of DUD-E, AlphaFold, and Apo structure screening results, respectively. The data of $p < 0.05$ has been bold.

### Multi-stage screening combined with consensus scoring

In order to use the advantages of consensus scoring to improve the ability of virtual screening and reduce the computational resources consumed in the whole process of consensus scoring, we proposed a multi-stage screening combined with consensus scoring (Fig. 5a). It was worth noting that since our study above has performed virtual screening on 51 targets, and the scores of different scoring functions between all proteins and small molecules were known, multi-stage screening (plan A, B, C, D, E, F) did not actually perform virtual screening, but uses corresponding known data, and the time consumed was also a theoretical estimate calculated based on known data.

**Figure 5.**
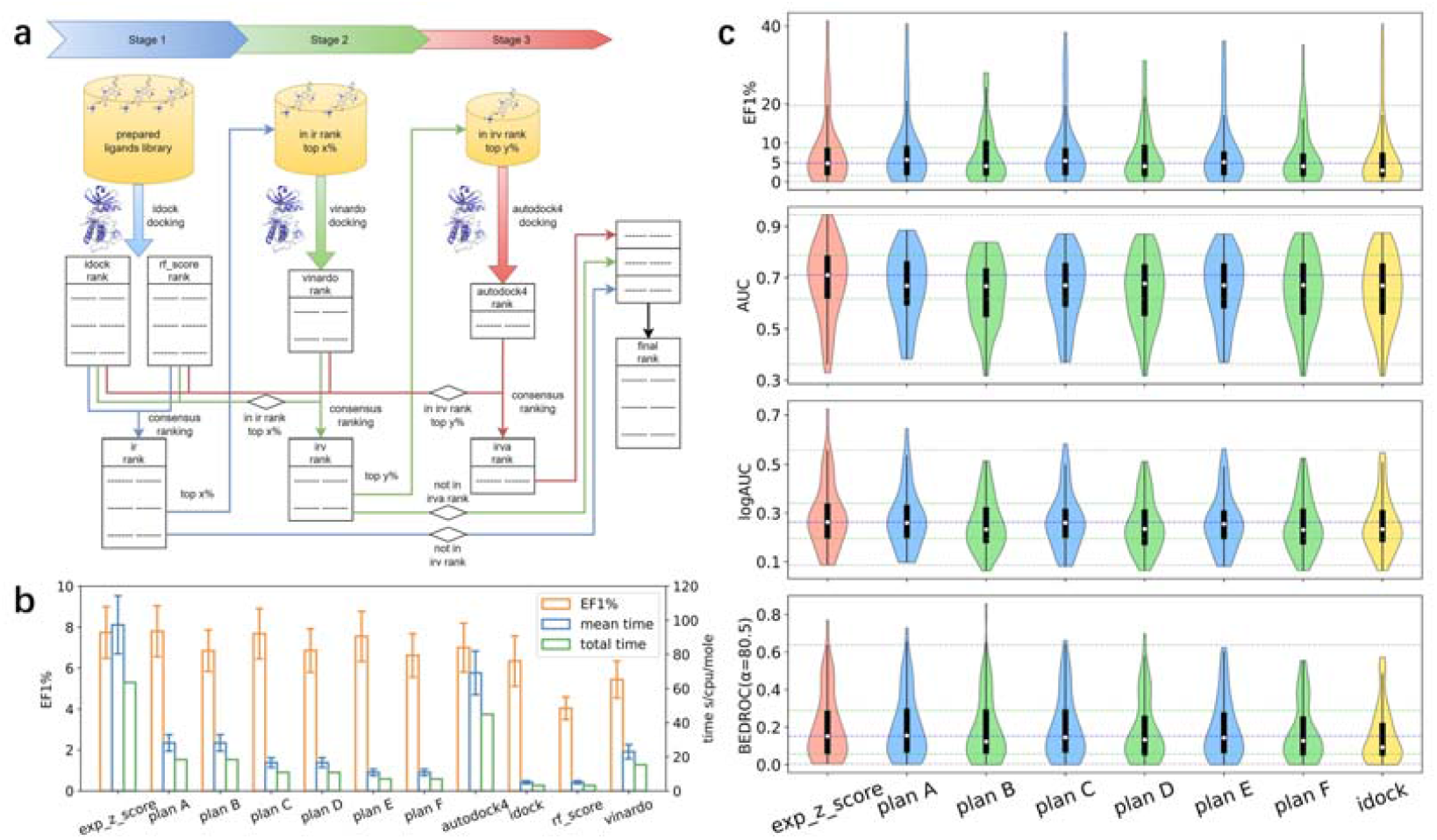
Multi-stage screening combined with consensus scoring. (a) Multi-stage screening flow chart combined with consensus scoring. Prepare small molecules, proteins and search space, and then start multi-stage screening. In the first stage, idock docking is used for virtual screening to obtain idock rank and rf_score rank, and then ir rank is obtained through the consensus scoring of idock rank and rf_score rank. The top x% of the ir rank are taken to enter the second stage. In the second stage, we use the vinardo scoring function to conduct virtual screening to obtain the vinardo rank, and then take the small molecules in the top x% of the ir rank in the idock rank and rf_score rank and all the small molecules in the vinardo rank for consensus scoring to obtain the irv rank. The top y% of the irv rank are taken to enter the third stage. In the third stage, the autodock4 scoring function is used to conduct virtual screening, and the autodock4 rank is obtained. Then the small molecules in idock rank, rf_score rank and vinardo rank that are y% before irv rank and all small molecules in autodock4 rank are scored by consensus to obtain irva rank. The final ranking is obtained by ranking the small molecules in irva rank, the small molecules in irv rank that are not in irva rank, and the small molecules in irv rank that are not in irv rank according to the original order within the ranking. (b) The results of 51 targets in DUD-E database for exp_z_score consensus scoring (all with autodock4, idock, rf_score, vinardo as input), multi-stage screening with consensus scoring (plan A, C, E), multi-stage screening without consensus scoring (plan B, D, F, the flow chart is shown in Supplementary Information Figure S1) and single scoring function (autodock4, idock, rf_score, vinardo). Including the mean ± s.e.m of EF1% (orange), the mean ± s.e.m time of 51 targets (blue, unit: second/cpu/molecule, abbreviated as s/cpu/mole in the figure), and the total time of 51 targets (green, unit: second/cpu/molecule). (c) The violin chart shows the screening results of exp_z_score consensus scoring (all with autodock4, idock, rf_score, vinardo as input), multi-stage screening with consensus scoring (plan A, C, E), and multi-stage screening without consensus scoring (plan B, D, F) for 51 targets, EF1%, AUC, logAUC and BEDROC (α=80.5) metrics are used to evaluate the screening effect. For plan A and plan B, take top 40% in the first stage to enter the second stage, and take top 50% in the second stage to enter the third stage. For plan C and plan D, take top 20% in the first stage to enter the second stage, and take top 50% in the second stage to enter the third stage. For plan E and plan F, take top 10% in the first stage to enter the second stage, and take top 50% in the second stage to enter the third stage.

On the crucial EF1% index for evaluating the early enrichment ability of virtual screening, the multi-stage screening plan A, C and E maintained a relatively high score (for the mean of EF1%, plan A = 7.80, plan C = 7.68, and plan E = 7.55) in comparison with the exp_z_score method (exp_z_score = 7.74, the *p* value for EF1% comparing between plan E and exp_z_score is 0.420). With the reduction of the time consumed for the multi-stage screening scheme, EF1% had a slight downward trend but no significant difference, and still keep a higher value. In addition, the EF1% of multi-stage screening with using consensus scoring (for plan E is 7.55) was higher than that without consensus scoring (for plan F is 6.62) (Fig. 5b and Table 6). These indicated that the multi-stage screening using consensus scoring would remarkably improve the virtual screening results.

**Table 6.**
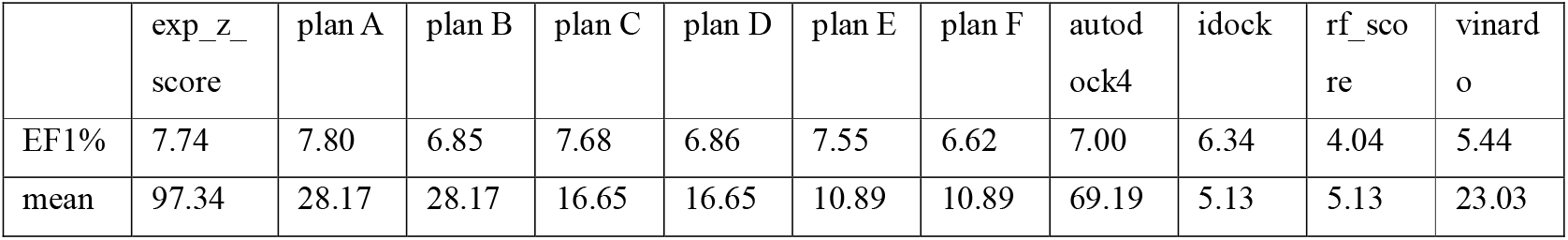
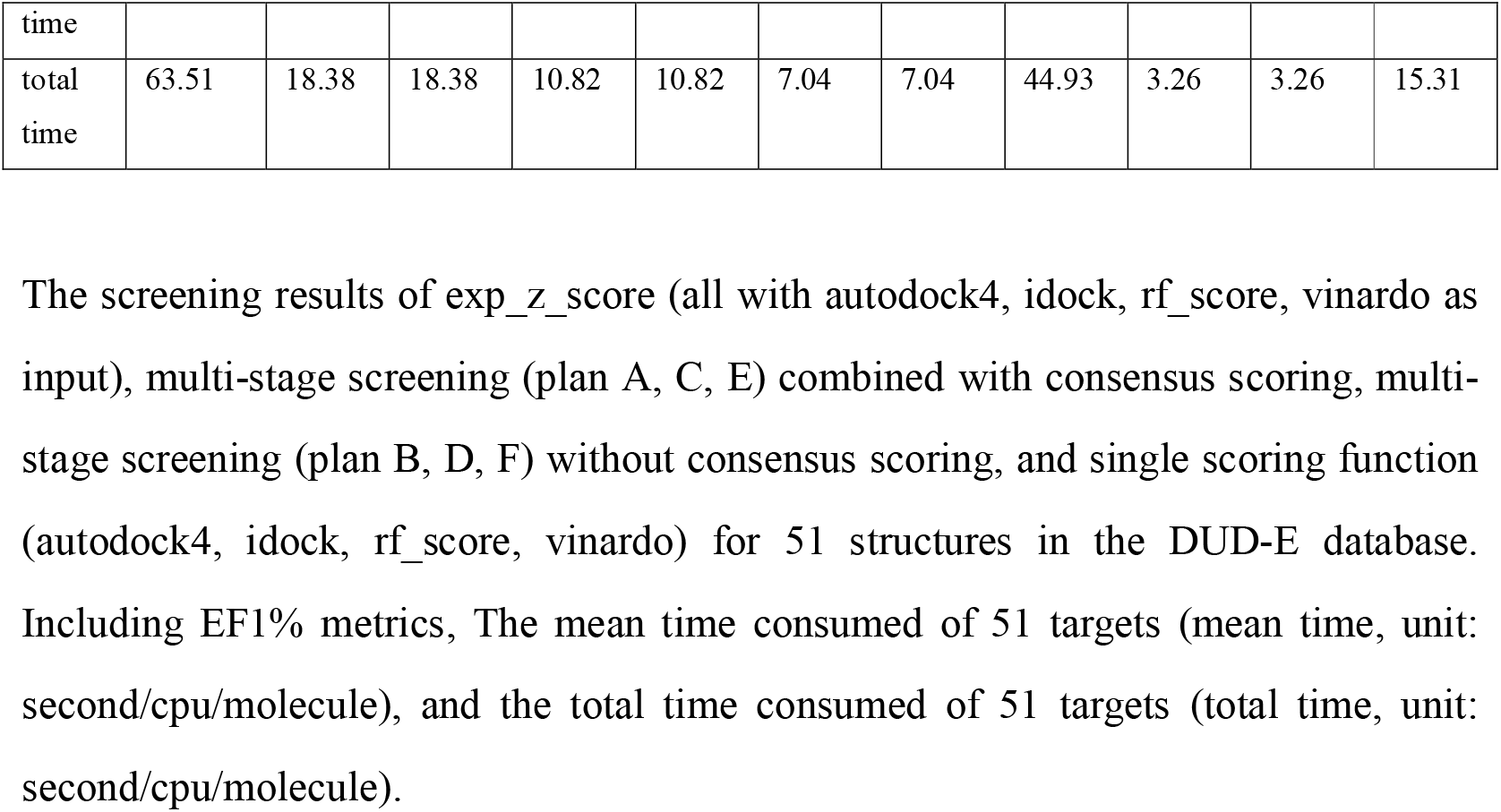
Early enrichment capacity and time consumed of multi-stage screening combined with consensus scoring.

BEDROC is an important metric of early enrichment capacity (α=80.5). We also observed that multi-stage screening plan A, C and E with consensus scoring (for the mean of BEDROC(α=80.5), plan A = 0.203, plan C = 0.197, plan E = 0.189) exhibited a high score of BEDROC(α=80.5) similar to exp_z_score (the mean of BEDROC(α=80.5) was 0.204, the *p* value for BEDROC(α=80.5) between plan E and exp_z_score was 0.032) (Fig. 5c), but a worse score in terms of AUC (for the mean of AUC, plan A =0.666, plan C = 0.659, plan E = 0.657, exp_z_score = 0.686. The *p* value for AUC between plan E and exp_z_score was 0.0005) and logAUC (for the mean of logAUC, plan A = 0.276, =plan C = 0.268, =plan E = 0.262, exp_z_score = 0.290. The *p* value for AUC between plan E and exp_z_score is 0.0002) metrics. These results demonstrated that multi-stage screening combined with consensus scoring had a great advantage in early enrichment ability which is of great significance for drug discovery.

There was no significant difference between Plan E and exp_z_score in the term of EF1% values (p=0.420), while the mean calculation speed of each small molecule of Plan E (7.04 second/cpu/molecule) was nearly 9 times faster than exp_z_score (63.51 second/cpu/molecule). Therefore, multi-stage screening combined with consensus scoring can use the advantages of consensus scoring to synthesize each single scoring function to improve the ability of virtual screening effect, and significantly reducing the computational resources consumed in the whole process of consensus scoring. The multi-stage screening combined with consensus scoring had higher early enrichment ability and less computational resource consumption, which was of great significance for large-scale drug screening.

### Relationship between scoring score and hit rate

In addition, we also analyzed the result of four single scoring functions and exp_z_score function for DUD-E structures and AlphaFold structures of 51 targets, and counted distributions and hit rates of small molecules.

To understand the relationship between the scoring and hit rate of small molecules in various scoring methods, we calculated the hit rate of active small molecules in the corresponding score segment of each scoring method. We found that the smaller the score of the autodock4, idock, and vinardo scoring functions, the more likely the small molecule was to combine with the protein. Moreover, as the score decreases, the hit rate tended to increase (Fig. 6a, b, e, f, i, j blue and green curves). While, according to the core algorithms of rf_score and exp_z_score functions, the higher values demonstrated that these small molecules were more likely to combine with the protein. With the increase of the score, the hit rate tended to increase in terms of rf_score and exp_z_score (Fig. 6c, d, g, h blue and green curves). For different methods, small molecules with better scores were more likely to be active small molecules and deserve more attention. This was also one of the bases for multi-stage screening to select the top small molecules for further screening in the next stage.

**Figure 6.**
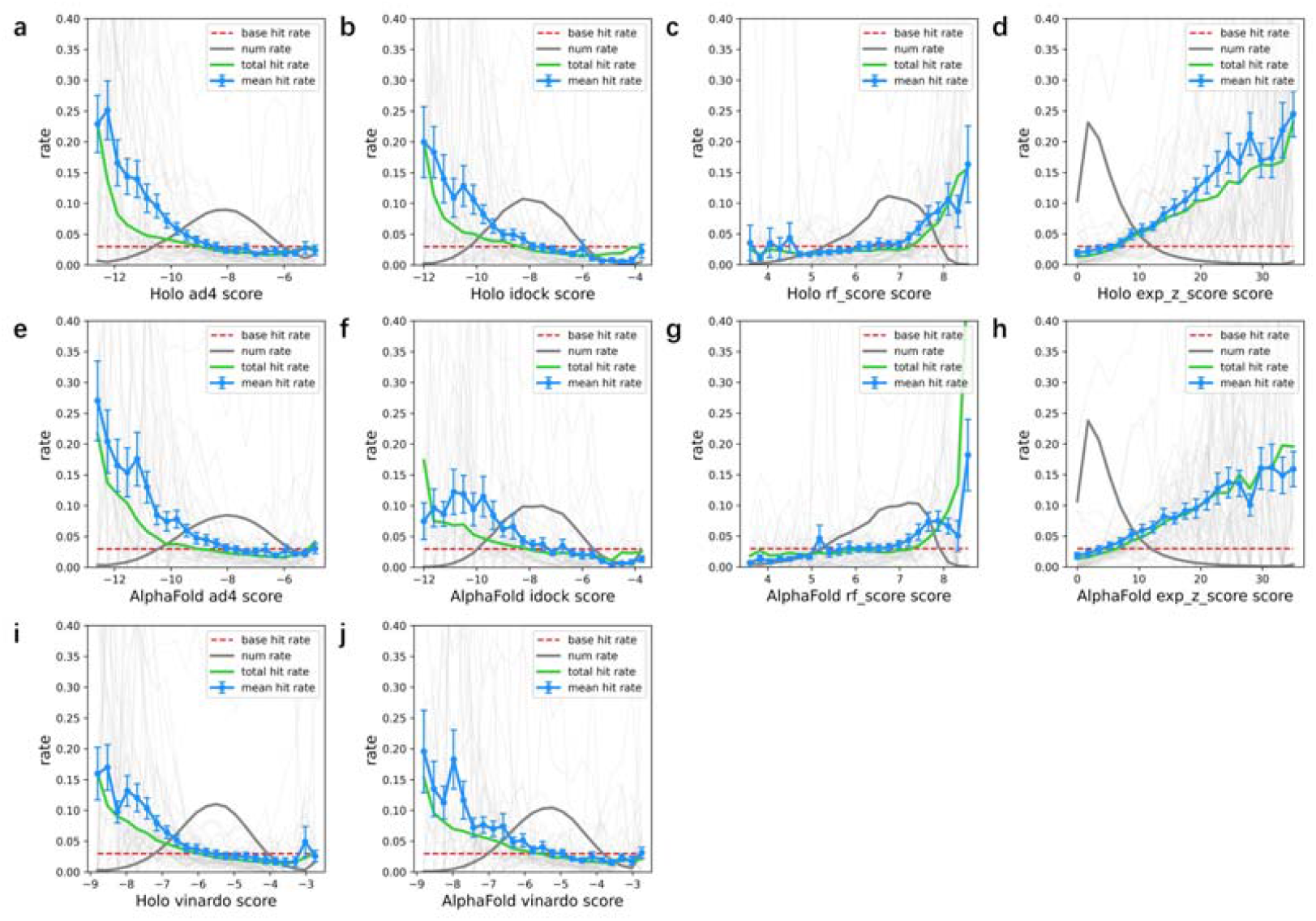
Relationship between scoring score and hit rate. (a-j) The hit rate for 51 targets according to the results of autodock4, idock, rf, vinardo and exp_z_score. The red dotted lines represent the base rate that the number of total active small molecules in 51 targets / the number of total small molecules in 51 targets, the base rate of each subgraph is about 3.0%. The gray dotted lines represent the number of small molecules. The green dotted lines represent the total hit rate of 51 targets. The blue dotted lines represent the mean ± s.e.m hit rate of 51 targets. The light gray dotted lines represent the hit rate of each target of 51 targets. The abscissas of a, b, c, e, f, g, i, j were divided into 22 segments according to the score, of which the end two segments respectively contain data that is greater than or equal to and less than the boundary, and the remaining 20 segments contain data that is left closed and right open within the segment. The abscissa of d and h were divided into 21 segments according to the score. One segment at the right end contains data that is greater than or equal to the boundary, and the other 20 segments contain data that is left closed and right open within the segment. The a, b, c, d, i are the screening results of structures in the DUD-E database, and the e, f, g, h, j are the screening results of AlphaFold structures.

Since the number of molecules in some fraction segments of high and low scores is small, the hit rate error corresponding to the fraction segment may be large. The curve of score and hit rate is of great significance for evaluating the scores given by different methods. It is helpful for researchers to set threshold values based on scores and give priority to small molecules that are more likely to be active compounds. For example, if the score of a small molecule on a protein is less than −9 by using the autodock4 scoring function for docking, the small molecule is more likely to be an active small molecule.

## Discussion

There is no doubt that the usage of AI/ML (intelligence and machine learning) is accelerating in drug discovery^30^. Researchers use machine learning methods, called MPxgb(AD), to identified Alzheimer’s disease-related genes which demonstrate good performance in experimental verification^31^. Also, virtual screening has been applied for developing new COVID-19 therapies from almost 4000 approved drugs^32^. Deep learning has also been employed in structure-based protein studies^33,34^. Here, to evaluate AlphaFold based on deep-learning for VS, we compare the performance of the protein structure of AlphaFold protein and DUD-E database, and apply a variety of docking software, multiple scoring functions and consensus scoring methods to perform VS.

For virtual screening, Holo structures show generally better performance than that of Apo structures^25,26^. Similar observations are also found in our results (Fig 2 and 4). Moreover, we find that in the test of 51 selected targets, the virtual screening effect of protein structures in DUD-E database is better than that of AlphaFold structures as a whole, but there are 18 targets (about 35%) whose AlphaFold structures are equivalent to that of DUD-E protein, and 13 targets (about 25%) whose AlphaFold structures are better than that of DUD-E. In the test of 23 selected targets, the virtual screening effect of protein structures in DUD-E database is slightly better than that of AlphaFold structure, and AlphaFold structures are somewhat better than that of Apo structure. Our result demonstrate that AlphaFold exhibits a relative worse performance compared to Holo stuctures for VS, which is consistent with recent findings when comparing with Holo PDB structures^8^. Our results have guiding significance for the selection of protein structures in virtual screening. Generally, when the Holo structures are available, the Holo structures are preferred. When the Holo structures are not available, the AlphaFold structures can be used preferentially. However, for specific targets, the selection of structure still needs to be careful.

At present, the AlphaFold protein structure database has more than 200 million entries, providing human proteome and proteome of 47 other organisms. Although AlphaFold is only used to predict the monomer protein structures, AlphaFold-Multimer can be used to predict multi-chain protein complexes^35^. In addition, ColabFold^36^, which is more user-friendly and easy to use, instead of installing the AlphaFold program and downloading the required large database locally. Combined with our conclusions, the super large-scale protein structure database and low-cost and fast accurate method from sequence to three-dimensional protein structure provided by AlphaFold are of excellent use value for virtual screening, and of great significance for speeding up drug discovery and reducing drug development costs.

We test 13 consensus scoring methods and find the exp_z_score is generally better than other methods, which shows to be higher with significant difference than ECR in terms of BEDROC(α=20.0) and EF10%. The advantage of exp_z_score consensus method may be that it used a more reasonable Z-score standardized method and exponential function that is more regular for virtual screening. The Z-score standardization method can standardize the scoring data group of the scoring function for the small molecule set to the data group with a mean value of 0 and a standard deviation of 1. Compared with the standardized method of Rank, the Z-score standardization method can better retain the difference information of the same scoring function for different small molecules. In addition, the Z-score standardization method is less affected by the maximum and minimum values than the AASS standardization method.

With the rapid expansion of the small molecules library available for virtual screening and the super extensive protein structure database provided by AlphaFold, the large-scale virtual screening will play an increasingly important role in drug discovery. To reduce the computing time of virtual screening, improve the computing efficiency, save computing resources, and be able to use a variety of different scoring functions and consensus scoring methods, we propose a multi-stage screening combining consensus scoring methods. The researchers have development a platform, VirtualFlow, for large-scale screening^24,37^. This platform has been successfully applied to drug discovery of SARS-CoV-2^38^ and broad-spectrum antivirals^39^. Our results showed that the mean of EF1% has no significant difference compared with consensus scoring using multi-stage screening, but its speed is about 9 times that of consensus scoring. Multi-stage screening combined with consensus scoring can flexibly adjust the number of small molecules screened at each stage to adjust the time and screening effect. It only uses a small amount of computing resources and has a high early enrichment capacity. Compared with the traditional multi-stage screening, the multi-stage screening combined with consensus scoring has great advantages in the early enrichment ability. Using multi-stage screening combined with consensus scoring for large-scale drug screening is of great significance for speeding up drug discovery and reducing drug development costs.

Alphafold has learned a protein structure energy function which does not need coevolution information to reach a high precision^40^. These significant findings reveal practical applications of AlphaFold in predicting protein structure when multiple sequence alignment is unavailable. Although AlphaFold could produce protein structure information with high-accuracy, there is a need to better harness AlphaFold for VS^7,8^. We optimize the AlphaFold structure by molecular dynamics in order to improve the virtual screening. This seems like a potential approach from the results of wee1 target shown in the Supplementary Information Table S19. But we can’t split the good structures from other structures for virtual screening without the active and decoy label. In other words, we can’t split the good structures from other structures by some simple indication such as the best score or the mean score. Nichols et al. also suggest that molecular dynamics could improve virtual screening results^41^. In addition, approaches based on machine learning would be a potential method to complement the use of AlphaFold2 for drug discovery. Researchers find that rescoring our docking poses with three machine learning-based scoring functions increased the accuracy of predictions^7^. Furthermore, post-modeling refinement strategies may be helpful to improve the chances of success^8^.

We use AutoSite^42^ to determine the search space, which has the advantage that we can flexibly describe the binding pocket structure of proteins. However, the multiple binding site structures given by AutoSite still need to be selected according to the position of eutectic small molecules relative to the protein structure. In addition, the combination pocket structure provided by AutoSite may be too large or too small in a few cases. To avoid the influence of human operation on the experimental results, we do not adjust the search space separately when the combined bag structures are too large or too small.

For our research, there are still some limitations. The number of targets used to compare Holo structures and AlphaFold structures is 51, and the number of targets used to compare Holo structure, Apo structure and AlphaFold structure is 23. Although some valuable data have been obtained from these targets, more targets can be used for testing to obtain more reliable data. In the treatment before the virtual screening of Holo structure, Apo structure and AlphaFold structure, we remove the non-protein structural parts from all structures. In the practical application of virtual screening, the non-protein components in Holo structures and Apo structures, such as water molecules, coenzymes, and metal ions, may be considered. The docking used in the study regards protein structure as rigid, while small molecules as flexible. Therefore, flexible docking^43^ reviewed by Spyrakis et al., that is, the amino acid residues in protein binding pockets are considered flexible and small molecules are considered flexible, which may be a method worth trying. The limitation of the curve drawn based on the relationship between scoring and hit rate lies in the high proportion of active small molecules in the DUD-E database and the small number of small molecules, which is quite different from the large-scale virtual screening. In the test of consensus scoring method, we only consider the scoring of small molecules by the scoring function of docking software, but do not consider docking conformation. Taking into account docking conformation for consensus scoring may be a method worth trying. In the multi-stage screening test, the number of stages divided is three, and only three groups of schemes are set according to the proportion of small molecules entering the next stage in each stage. What’s more, the small molecule set used for testing has fewer small molecules, which is several orders of magnitude less than the small molecule library of super large-scale virtual screening. Therefore, although our research can provide some guidance for super large-scale virtual screening, it is difficult to give a specific implementation plan for large-scale virtual screening. Further work is needed to develop a specific implementation plan for large-scale virtual screening.

## Methods

### Targets and structures

To evaluate the effectiveness of AlphaFold structure in virtual screening, we used the targets in the DUD-E database. The DUD-E database contains 102 protein targets, each of which includes protein structure (from the PDB database), eutectic small molecules to determine the docking software search space, active small molecules and decoy small molecules. It is widely utilized to assess the ability of virtual screening to distinguish active compounds from decoy compounds.

Considering the calculation time of virtual screening, the diversity of protein types and whether the AlphaFold structure can be used for virtual screening (structure availability and whether there is an obvious binding site), we selected 51 targets in the DUD-E database as our test targets representing eight types of protein targets namely protease, nuclear receiver, kinase, cytochrome P450, ion channel, GPCR, other enzymes and miscellaneous proteins (Fig. 1a). In total, more than 400,000 small molecules which interact with these 51 targets were used. The protein structures provided by DUD-E database corresponding to the 51 selected targets are used as the test targets of Holo structures.

The AlphaFold structures we used can be obtained from AlphaFold Protein Structure Database^44^ (https://alphafold.com/) by the uniprot accessions except HIVPR which is predicted by ColabFold. The uniprot accessions for the 51 targets are provided in the Supplementary Information Table S20. The 50 structures obtained from AlphaFold Protein Structure Database and the HIVPR structure predicted by ColabFold are used as the test targets of AlphaFold structures.

We tried to collect the Apo structures for the 51 selected targets in DUD-E database. However, some proteins cannot or are difficult to find their Apo structures in the PDB database and some structures have no apparent binding pocket. Only 23 of the 51 targets were found available Apo structures with binding pocket. They are used as the test targets of Apo structures. The information including the PDB ID and the resolution of 23 Apo structures are provided in the Supplementary Information Table S20.

### ColabFold

Milot Mira et al. developed ColabFold based on Google Colab in order to enable researchers without relevant hardware resources to use AlphaFold2^36^.

Because HIVPR and HIVRT are non-monomer proteins, their protein structures cannot be directly obtained in the AlphaFold Protein Structure Database, so ColabFold is used for structure prediction. The input sequences and configurations are provided as Supplementary Method in the Supplementary Information.

### Protein structure preparation

Each protein structure that is the target of the virtual screening test is processed as follows:

1. Use PDBFixer 1.8.1 (https://github.com/openmm/pdbfixer) to replace non-standard residues, remove heterologous substances (including water molecules, coenzymes, metal ions, etc.), and add missing atoms to obtain the receptor_fixed.pdb.
2. Use the script provided by AutoDock Vina to execute the following commands: prepare_receptor -r receptor_fixed.pdb -o receptor.pdbqt -A hydrogens The final obtained pdbqt format file is the prepared protein file, which can be directly used as the input of AutoDock Vina, Qvina2 and idock to perform docking.

### Compounds preparation

We get the active small molecule files actives_final.sdf.gz and decoy small molecule files decoys_final.sdf.gz corresponding to each target from DUD-E Database. Next, decompress and use openbabel 3.1.0^45^ (https://github.com/openbabel/openbabel) to perform split conversion to obtain the mol2 format file corresponding to each small molecule, and then use the script provided by AutoDock Vina for each small molecule to execute the following commands for hydrogenation and preparation of small molecule file formats:

prepare_ligand -l molecule.mol2 -o molecule.pdbqt -A hydrogens

The final obtained pdbqt format file is the prepared small molecule file, which can be directly used as the input of AutoDock Vina, Qvina2 and idock to perform docking.

### Search space

AutoSite 1.0.0^42^ is used to predict binding sites for each protein structure that is the target of the virtual screening test, and PyMOL (TM) Molecular Graphics System, Version 2.6.0a0 Open-Source^46^ is used to visually inspect and select binding sites.

For each protein structure in the DUD-E database, the binding site predicted by AutoSite is selected by the eutectic small molecule ligand.mol2 in the DUD-E database.

For the AlphaFold structure and Apo structure, align the AlphaFold structure and Apo structure to the structure in the DUD-E database. Then, select the binding site by referring to the position of eutectic small molecules in the DUD-E database.

After determining the protein structure binding site, use the PandasPdb module of biopandas 0.2.9 to calculate the spatial coordinate information of the binding site structure, including size and location, and then add 8 Å to each of the x, y, and z dimensions of the binding site structure size as the search space for AutoDock Vina, Qvina2, and idock.

### Docking

Three docking software AutoDock Vina 1.2.3, QuickVina 2.1 (Qvina2) and idock 2.2.3 are used. And five scoring functions including autodock4 and vinardo in AutoDock Vina, qvina scoring function, idock scoring function, and rf_scoring function in idock were employed. For each docking, the seed used is 20011204, and only the best docking position is considered as output. The exhaustiveness parameter used by AutoDock Vina and Qvina2 is 1, and the tasks parameter used by idock is 8. Neither flexible docking nor hydration docking is used.

The used autodock4 (ad4) scoring function and vinardo scoring function are the scoring functions provided by AutoDock Vina 1.2.3. When using the autodock4 scoring function and vinardo scoring function, perform docking respectively to obtain the scoring of a small molecule by autodock4 scoring function and the scoring of a small molecule by vinardo scoring function. Affinity map is also required when using the autodock4 scoring function. For each docking the required affinity map was calculated by the prepare_gpf.py script that AutoDock Vina provided.

The qvina scoring function used is QuickVina 2.1 scoring function.

The score function used by idock and rf_score is provided by idock 2.2.3. When idock 2.2.3 is executed, only one time of docking would be performed to obtain the score of idock and rf_score functions for a small molecule.

### Consensus scoring

For every target, to integrate different scoring functions, different scoring functions may need to be standardized. Common standardization methods include Rank, AASS (Average of auto-scaled scores), Z-score, etc. Only after the scores between different scoring functions are standardized can they be reasonably calculated and compared. According to the direction of scoring, the scoring function can be divided into two categories. The first category is that the larger the scoring value represents the stronger the binding ability of small molecules to protein. The second category is that the higher the score represents the weaker the binding ability of small molecules to proteins is. The valid scores of the first category are the scores greater than 0; The valid scores of the second category are the scores less than 0.

At the same time, another key to consensus scoring is how to process standardized scores to obtain the final consensus scoring. We selected four calculation methods: Summation (Sum), Best, Worst, and Summation after Exponentiation (Exp).

### Rank

Rank as the standardized score.

For the first category, the scores are sorted from the largest to the smallest. For the second category, the scores are sorted from the smallest to the largest. That is, the scores from two categories are ranked from good to bad. And then, use rank as standardized score for the two categories. The smaller the standardized score, the better.

### Z-score

Standardize score *S*_*ij*_ for small molecule *j* on scoring function *i* by Z-score to obtain the standardized score *Z*_*ij*°_

The mean of the effective scores on the scoring function *i* is 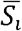, and the standard deviation of that is *σ*_*i*_.

The scores from the first category are standardized by the following formula:

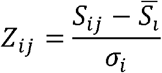

The scores from the second category are standardized by the following formula:

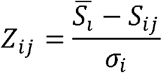

For the two categories, the larger the standardized score, the better.

### AASS

Standardize score *S*_*ij*_ for small molecule *j* on scoring function *i* by AASS to obtain the standardized score *A*_*ij*_.

For the first category and the scoring function *i*, the largest score is *Best*_*i*_, the smallest score is *Worst*_*i*_. For the second category and the scoring function *i*, the smallest score is *Best*_*i*_, the largest score is *Worst*_*i*_. That is, for the two categories, the best score is *Best*_*i*_, the worst is *Worst*_*i*_.

The scores from the two categories are standardized by the following formula:

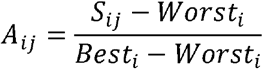

For the two categories, the larger the standardized score, the better.

### Sum

The standardized score for small molecule *j* on scoring function *i* is *V*_*ij*_. The consensus score *Sum*_*j*_ for small molecule *j* is calculated by the following formula:

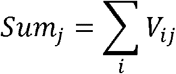

### Best

The consensus score *Best*_*j*_ for small molecule *j* is the best standardized score in every scoring function.

### Worst

The consensus score *Worst*_*j*_ for small molecule *j* is the worst standardized score in every scoring function.

### Exp

The standardized score for small molecule *j* on scoring function *i* is *V*_*ij*_. The consensus score *Exp*_*j*_ for small molecule *j* is calculated by the following formula:

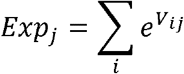

### Consensus function

According to the combination of standardization methods (3) and calculation methods (4), 12 consensus scoring methods were obtained. The combination and name are as follows:

|  | Rank | Z-score | AASS |
| --- | --- | --- | --- |
| Sum | sum_rank | sum_z_score | sum_aass |
| Best | best_rank | best_z_score | best_aass |
| Worst | worst_rank | worst_z_score | worst_aass |
| Exp | exp_rank | exp_z_score | exp_aass |

For example, the complete description of exp_z_score is as follows:

The mean of the effective scores on the scoring function *i* is 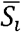, and the standard deviation of that is *σ*_*i*_. The standardized score for small molecule *j* on scoring function *i* is *Z*_*ij*_.

The scores from the first category are standardized by the following formula:

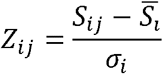

The scores from the second category are standardized by the following formula:

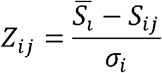

The final consensus score *Exp*_*j*_ for small molecule *j* is calculated by the following formula:

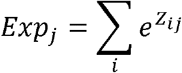

The larger the final consensus score, the more likely it is an active small molecule.

While exp_rank (ECR) has changed the method of Exp: The rank for small molecule *j* on scoring function *j* is *R*_*ij*_. The number of the effective scores on the scoring function *i* is *n*_*i*_. The consensus score *Exp*_*j*_ for small molecule *j* is calculated by the following formula:

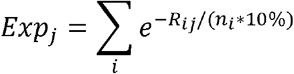

The larger the final consensus score, the more likely it is an active small molecule. In addition, Rank by Vote (RBV) consensus scoring method was tested.

### RBV

The threshold value is set to x=top10% and the ranking of small molecule *j* in the scoring function *i* is within the threshold value x, one vote will be obtained. The sum of votes of small molecule *j* on all scoring functions is the final score. Small molecules with the same number of votes are randomly ranked.

### Multi-stage virtual screening with consensus scoring

Although the virtual screening effect of the best consensus scoring obtained in the test is better than that of a single scoring function, consensus scoring is based on multiple scoring functions. The computing resources consumed in the whole process of consensus scoring are the sum of all the computing resources consumed by the involved single scoring function and docking software, which consumes more computing resources. In order to use the advantages of consensus scoring to improve the ability of virtual screening and reduce the computational resources consumed, we propose a multi-stage screening combined with consensus scoring.

Prepare small molecules, proteins and search space, and then start multi-stage screening. In the first stage, idock docking is used for virtual screening to obtain idock rank and rf_score rank, and then ir rank is obtained through the consensus scoring of idock rank and rf_score rank. The top x% of the ir rank are taken to enter the second stage. In the second stage, we use the vinardo scoring function to conduct virtual screening to obtain the vinardo rank, and then take the small molecules in the top x% of the ir rank in the idock rank and rf_score rank and all the small molecules in the vinardo rank for consensus scoring to obtain the irv rank. The top y% of the irv rank are taken to enter the third stage. In the third stage, the autodock4 scoring function is used to conduct virtual screening, and the autodock4 rank is obtained. Then the small molecules in idock rank, rf_score rank and vinardo rank that are y% before irv rank and all small molecules in autodock4 rank are scored by consensus to obtain irva rank. The final ranking is obtained by ranking the small molecules in irva rank, the small molecules in irv rank that are not in irva rank, and the small molecules in irv rank that are not in irv rank according to the original order within the ranking. The x and y parameters can be adjusted flexibly. Three groups of schemes are set, Plan A(x=40, y=50),Plan C(x=20,y=50),Plan E(x=10,y=50).

### Multi-stage virtual screening without consensus scoring

With the rapid expansion of the small molecules library that can be used for virtual screening, large-scale virtual screening is also increasingly widely used. Therefore, it is of great significance to improve the efficiency of virtual screening and reduce the computing resources consumed by virtual screening to save the cost of large-scale virtual screening and accelerate the drug screening process. Several studies have used multi-stage screening to improve the efficiency of virtual screening. Multi-stage screening uses rough but fast screening in the first stage, and more refined but slow screening in the second part of the stage, so as to give consideration to the early enrichment ability and computational efficiency of virtual screening.

Prepare small molecules, proteins and search space, and then start multi-stage screening. In the first stage, idock docking is used for virtual screening to obtain idock rank, the top x% of the idock rank are taken to enter the second stage. In the second stage, we use the vinardo scoring function to conduct virtual screening to obtain the vinardo rank, the top y% of the vinardo rank are taken to enter the third stage. In the third stage, the autodock4 scoring function is used to conduct virtual screening, and the autodock4 rank is obtained. The final ranking is obtained by ranking the small molecules in autodock4 rank, the small molecules in vinardo rank that are not in autodock4 rank, and the small molecules in idock rank that are not in vinardo rank according to the original order within the ranking. The x and y parameters can be adjusted flexibly. Three groups of schemes are set, Plan B(x=40, y=50), Plan D(x=20,y=50),Plan F(x=10,y=50).

### Metrics

We used AUC (area under the receiver operating characteristic curve), logAUC (area under the semilogarithmic receiver operating characteristic curve), EF1% (enrichment factor 1%), EF5%, EF10% and BEDROC (Boltzmann-enhanced discrimination of receiver operating characteristic, α=321.9, α=80.5, α=20.0) as benchmark metrics to assess the result. LogAUC is the area under the semilogarithmic ROC curve, and the early enrichment capacity has a greater weight in logAUC than AUC^11^. Enrichment factors (1%, 5%, and 10%) are used to truncate the assessment of early enrichment capacity under specific thresholds, which are closely related to the practical application of virtual screening. The BEDROC^47^ was proposed to consider all small molecules by adjusting α value for the weight of early enrichment capacity. The greater the early enrichment capacity, the larger α weight value.

### Hit rate

In a collection of small molecules, the number of active compounds in the collection is *actives*, the number of all compounds in the collection is *total*. *Hit rate* is calculated by the following formula:

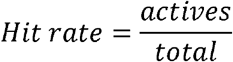

For a collection of small molecules in the top x% of a rank, the number of active compounds in the collection is *actives*(*x*%), the number of all compounds in the collection is *total*(*x*%). *Hit rate*(*x*%) is calculated by the following formula:

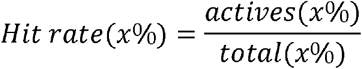

### Enrichment factor

For a collection of small molecules in the top x% of a rank, the enrichment factor x% (EFx%) is calculated by the following formula:

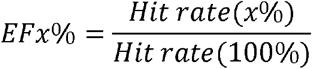

### BEDROC

BEDROC is defined as follows:

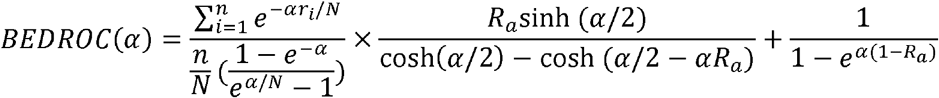

While the *n* is the number of active compounds, the *N* is the number of all compounds, the rate *Ra* = *n*/*N*, the *r*_*i*_ is the rank of *i* th active compound ^47^.

### Receiver Operating Characteristics (ROC) Curve and its area under the curve (AUC)

The Y-axis is the true positive rate (TPR, also named actives found rate) under a specific threshold. The X-axis is the false positive rate (FPR, also named decoys found rate) under a specific threshold.

TPR is calculated by the following formula:

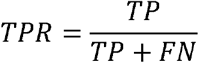

FPR is calculated by the following formula:

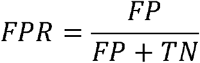

At a specific threshold, TP is the number of true positive compounds, FN is the number of false negative compounds, FP is the number of false positive compounds, and TN is the number of true negative compounds.

The area under the ROC curve is AUC, and its value range is [0,1]. The AUC corresponding to random screening is 0.5.

### Semilogarithmic Receiver Operating Characteristics (ROC) Curve and its area under the curve (logAUC)

After drawing ROC curve, let λ = 0.001. Points with abscissa less than λ are ignored and the abscissa of other points is performed logarithmic (*log*_10_) operation. That is, the point (*x*_*i*_, *y*_*i*_) in the ROC curve, the point (*log*_10_(*x*_*i*_),*y*_*i*_) in the semilogarithmic ROC.

The abscissa value range of Semilogarithmic ROC curve is [-3,0]. The area of semilogarithmic ROC curve is *area*. Let *logAUC* = *area*/3, and its value range is [0,1].

### Time

Use the Linux time command to count the time, and use user time + sys time as the time used for virtual screening.

### System

All docking calculations are performed in containers created by images built with Docker 20.10.17 software. Docker is an open source lightweight virtualization technology (https://www.docker.com/).

The host system is Linux 5.4.0-131 generic.

### CPU

Intel(R) Xeon(R) Gold 6238R CPU @ 2.20GHz.

### Figure

Violin chart, ROC curve, scatter plot, fitting line and histogram are drawn with matplotlib 3.6.1^48^.

The three-dimensional structure diagram of small molecules and protein is drawn with PyMOL (TM) Molecular Graphics System, Version 2.6.0a0 Open-Source^46^.

## Author contributions

**Yanfei Peng**: Conceptualization, Methodology, Software, Validation, Formal analysis, Investigation, Data Curation, Visualization, Writing - Original Draft, Writing - Review & Editing. **Xia Wu**: Resources, Data Curation, Writing - Review & Editing, Visualization. **Liang Lin:** Conceptualization, Writing - Review & Editing. **Zhiluo Deng**: Investigation, Data Curation, Writing - Review & Editing. **Limin Zhao**: Conceptualization, Investigation, Data Curation, Writing - Original Draft, Writing - Review & Editing. **Hao Ke**: Conceptualization, Investigation, Data Curation, Writing - Original Draft, Writing - Review & Editing.

## Data availability

Data will be made available on request.

