## supplementary for "Assessment of AlphaFold structures and optimization methods for virtual screening"

Supplementary Table 1. 51 targets selected from the DUD-E database with the class information and with the PDB ID and the resolution of 23 Apo structures.

| Target | Apo PDB ID | Resolution (Å) | Classes |
| --- | --- | --- | --- |
| ada | 1vfl | 1.80 | Other Enzymes |
| akt1 |  |  | Kinase |
| akt2 |  |  | Kinase |
| ampc | 3bls | 2.30 | Other Enzymes |
| aofb |  |  | Other Enzymes |
| comt | 4pyk | 2.22 | Other Enzymes |
| cp2c9 | 1og2 | 2.60 | Cytochrome P450 |
| cp3a4 |  |  | Cytochrome P450 |
| cxcr4 |  |  | GPCR |
| def | 1bs5 | 2.50 | Other Enzymes |
| fa7 |  |  | Protease |
| fabp4 | 3q6l | 1.40 | Miscellaneous |
| fak1 |  |  | Kinase |
| fgfr1 | 4uwy | 2.31 | Kinase |
| fkb1a | 2ppn | 0.92 | Other Enzymes |
| gcr |  |  | Nuclear |
| glcm |  |  | Other Enzymes |
| grik1 |  |  | Ion Channel |
| hivpr |  |  | Protease |
| hs90a | 1yes | 2.20 | Miscellaneous |
| hxk4 | 3idh | 2.14 | Other Enzymes |
| inha |  |  | Other Enzymes |
| jak2 | 4fvp | 2.01 | Kinase |
| kif11 |  |  | Miscellaneous |
| kith |  |  | Other Enzymes |
| mapk2 | 2oza | 2.70 | Kinase |
| mcr |  |  | Nuclear |
| mk01 | 4iz7 | 1.80 | Kinase |
| mk10 | 4kkg | 2.40 | Kinase |
| nos1 | 1zvi | 2.00 | Other Enzymes |
| pa2ga | 1bbc | 2.20 | Other Enzymes |
| plk1 |  |  | Kinase |
| pnph | 1m73 | 2.30 | Other Enzymes |
| ptn1 | 2hnp | 2.85 | Other Enzymes |
| pur2 | 1mej | 2.00 | Other Enzymes |
| pygm |  |  | Other Enzymes |
| pyrd |  |  | Other Enzymes |
| rock1 |  |  | Kinase |
| rxra |  |  | Nuclear |
| sahh |  |  | Other Enzymes |
| thb |  |  | Nuclear |
| tryb1 |  |  | Protease |
| tysy | 3tms | 2.10 | Other Enzymes |
| wee1 |  |  | Kinase |
| xiap |  |  | Miscellaneous |
| casp3 |  |  | Protease |
| gria2 | 1fto | 2.00 | Ion Channel |
| ppard | 2gwx | 2.30 | Nuclear |
| bace1 | 2zhv | 1.85 | Protease |
| adrb1 |  |  | GPCR |
| adrb2 |  |  | GPCR |

Supplementary Table 2. The virtual screening results by autodock4 scorinig function of 51 targets based on the protein structures in DUD-E databases.

| Target | Active | Total | AUC | logAUC | BEDROC(α=321.9) | BEDROC(α=80.5) | BEDROC(α=20.0) | EF1% | EF5% | EF10% |
| --- | --- | --- | --- | --- | --- | --- | --- | --- | --- | --- |
| ada | 259 | 5716 | 0.266 | 0.057 | 0.001 | 0.010 | 0.017 | 0.00 | 0.23 | 0.15 |
| adrb1 | 451 | 16259 | 0.690 | 0.311 | 0.405 | 0.276 | 0.291 | 11.57 | 5.11 | 3.51 |
| adrb2 | 444 | 15602 | 0.709 | 0.332 | 0.487 | 0.319 | 0.318 | 13.52 | 5.68 | 3.74 |
| akt1 | 422 | 16931 | 0.647 | 0.220 | 0.113 | 0.088 | 0.129 | 3.80 | 1.99 | 1.87 |
| akt2 | 189 | 7108 | 0.688 | 0.244 | 0.067 | 0.106 | 0.171 | 3.71 | 2.75 | 2.54 |
| ampc | 62 | 2952 | 0.566 | 0.162 | 0.003 | 0.024 | 0.073 | 0.00 | 1.62 | 1.13 |
| aofb | 168 | 7079 | 0.566 | 0.181 | 0.019 | 0.040 | 0.097 | 1.81 | 1.55 | 1.49 |
| bace1 | 481 | 18613 | 0.752 | 0.304 | 0.149 | 0.166 | 0.254 | 6.03 | 4.70 | 3.58 |
| casp3 | 347 | 11068 | 0.643 | 0.249 | 0.217 | 0.159 | 0.206 | 5.51 | 3.63 | 2.77 |
| comt | 86 | 3994 | 0.473 | 0.116 | 0.000 | 0.000 | 0.017 | 0.00 | 0.00 | 0.23 |
| cp2c9 | 183 | 7730 | 0.595 | 0.227 | 0.176 | 0.136 | 0.178 | 7.13 | 2.95 | 2.35 |
| cp3a4 | 357 | 12254 | 0.483 | 0.144 | 0.090 | 0.057 | 0.073 | 2.53 | 0.95 | 1.01 |
| cxcr4 | 122 | 3522 | 0.830 | 0.467 | 0.662 | 0.479 | 0.533 | 14.85 | 9.51 | 6.81 |
| def | 160 | 5866 | 0.418 | 0.116 | 0.099 | 0.053 | 0.050 | 2.53 | 0.75 | 0.63 |
| fa7 | 185 | 6437 | 0.763 | 0.302 | 0.151 | 0.150 | 0.240 | 5.44 | 3.79 | 3.73 |
| fabp4 | 57 | 2899 | 0.814 | 0.439 | 0.299 | 0.398 | 0.457 | 19.98 | 9.54 | 5.10 |
| fak1 | 114 | 5486 | 0.575 | 0.163 | 0.000 | 0.004 | 0.056 | 0.00 | 0.35 | 1.32 |
| fgfr1 | 241 | 730 | 0.550 | 0.172 | 0.578 | 0.477 | 0.426 | 1.73 | 1.43 | 1.20 |
| fkb1a | 271 | 6098 | 0.767 | 0.287 | 0.146 | 0.144 | 0.232 | 3.00 | 3.33 | 3.29 |
| gcr | 562 | 15678 | 0.621 | 0.221 | 0.147 | 0.123 | 0.165 | 3.58 | 2.85 | 2.08 |
| glcm | 310 | 4123 | 0.746 | 0.275 | 0.064 | 0.177 | 0.278 | 1.95 | 3.29 | 3.07 |
| gria2 | 294 | 12310 | 0.640 | 0.219 | 0.065 | 0.067 | 0.137 | 2.04 | 2.45 | 2.04 |
| grik1 | 150 | 6743 | 0.586 | 0.206 | 0.230 | 0.114 | 0.126 | 6.04 | 1.73 | 1.60 |
| hivpr | 1378 | 37214 | 0.663 | 0.250 | 0.201 | 0.165 | 0.209 | 4.79 | 3.32 | 2.69 |
| hs90a | 125 | 5054 | 0.364 | 0.084 | 0.000 | 0.000 | 0.009 | 0.00 | 0.00 | 0.08 |
| hxk4 | 127 | 4917 | 0.585 | 0.171 | 0.005 | 0.018 | 0.063 | 0.79 | 0.63 | 0.95 |
| inha | 71 | 2380 | 0.699 | 0.353 | 0.495 | 0.360 | 0.376 | 14.57 | 6.20 | 4.65 |
| jak2 | 153 | 6716 | 0.532 | 0.160 | 0.030 | 0.035 | 0.078 | 1.31 | 1.05 | 1.24 |
| kif11 | 196 | 7084 | 0.778 | 0.357 | 0.326 | 0.270 | 0.337 | 11.36 | 6.13 | 4.39 |
| kith | 132 | 2968 | 0.460 | 0.167 | 0.097 | 0.122 | 0.172 | 2.33 | 3.19 | 2.20 |
| mapk2 | 205 | 6420 | 0.706 | 0.267 | 0.188 | 0.143 | 0.216 | 4.89 | 3.32 | 3.37 |
| mcr | 190 | 5425 | 0.651 | 0.291 | 0.418 | 0.301 | 0.287 | 11.63 | 4.64 | 3.32 |
| mk01 | 138 | 4744 | 0.721 | 0.255 | 0.170 | 0.088 | 0.154 | 2.93 | 2.18 | 2.39 |
| mk10 | 185 | 6861 | 0.546 | 0.167 | 0.022 | 0.046 | 0.087 | 2.18 | 1.19 | 1.35 |
| nos1 | 234 | 8283 | 0.560 | 0.177 | 0.030 | 0.037 | 0.087 | 1.73 | 1.03 | 1.50 |
| pa2ga | 127 | 5333 | 0.651 | 0.206 | 0.001 | 0.029 | 0.105 | 0.00 | 2.05 | 1.65 |
| plk1 | 152 | 6974 | 0.556 | 0.213 | 0.143 | 0.127 | 0.173 | 5.32 | 3.56 | 2.24 |
| pnph | 233 | 7230 | 0.694 | 0.277 | 0.205 | 0.162 | 0.243 | 5.17 | 4.30 | 3.43 |
| ppard | 285 | 13439 | 0.737 | 0.291 | 0.114 | 0.128 | 0.227 | 5.28 | 4.50 | 3.27 |
| ptn1 | 225 | 7628 | 0.852 | 0.416 | 0.358 | 0.336 | 0.413 | 13.83 | 7.03 | 5.34 |
| pur2 | 199 | 2913 | 0.903 | 0.724 | 0.990 | 0.933 | 0.872 | 14.13 | 13.12 | 8.45 |
| pygm | 112 | 4136 | 0.331 | 0.072 | 0.000 | 0.000 | 0.008 | 0.00 | 0.00 | 0.27 |
| pyrd | 134 | 6755 | 0.639 | 0.226 | 0.016 | 0.060 | 0.145 | 3.01 | 2.69 | 2.17 |
| rock1 | 202 | 6570 | 0.702 | 0.257 | 0.411 | 0.180 | 0.164 | 6.50 | 2.38 | 1.93 |
| rxra | 161 | 7845 | 0.845 | 0.478 | 0.333 | 0.375 | 0.520 | 19.37 | 10.69 | 7.02 |
| sahh | 190 | 3662 | 0.229 | 0.052 | 0.000 | 0.002 | 0.013 | 0.00 | 0.11 | 0.21 |
| thb | 168 | 7793 | 0.765 | 0.303 | 0.027 | 0.103 | 0.243 | 3.01 | 4.65 | 3.75 |
| tryb1 | 169 | 7820 | 0.753 | 0.362 | 0.350 | 0.272 | 0.350 | 13.05 | 6.86 | 4.62 |
| tysy | 308 | 7152 | 0.760 | 0.412 | 0.770 | 0.531 | 0.470 | 14.72 | 7.74 | 4.90 |
| wee1 | 136 | 6324 | 0.937 | 0.758 | 0.970 | 0.824 | 0.781 | 45.02 | 15.30 | 7.87 |
| xiap | 129 | 5312 | 0.879 | 0.611 | 0.860 | 0.668 | 0.650 | 33.41 | 11.65 | 7.21 |

Supplementary Table 3. The virtual screening results by idock scorinig function of 51 targets based on the protein structures in DUD-E databases.

| Target | Active | Total | AUC | logAUC | BEDROC(α=321.9) | BEDROC(α=80.5) | BEDROC(α=20.0) | EF1% | EF5% | EF10% |
| --- | --- | --- | --- | --- | --- | --- | --- | --- | --- | --- |
| ada | 262 | 5734 | 0.410 | 0.107 | 0.002 | 0.020 | 0.049 | 0.00 | 0.54 | 0.76 |
| adrb1 | 458 | 16416 | 0.645 | 0.207 | 0.087 | 0.061 | 0.105 | 2.19 | 1.44 | 1.64 |
| adrb2 | 447 | 15702 | 0.678 | 0.235 | 0.073 | 0.077 | 0.157 | 2.24 | 2.73 | 2.35 |
| akt1 | 423 | 16999 | 0.708 | 0.242 | 0.065 | 0.066 | 0.140 | 2.85 | 2.27 | 2.32 |
| akt2 | 190 | 7142 | 0.776 | 0.335 | 0.311 | 0.213 | 0.291 | 7.41 | 5.26 | 3.95 |
| ampc | 62 | 2964 | 0.534 | 0.160 | 0.103 | 0.052 | 0.083 | 1.65 | 1.62 | 1.29 |
| aofb | 168 | 7099 | 0.669 | 0.234 | 0.028 | 0.050 | 0.150 | 1.21 | 2.51 | 2.62 |
| bace1 | 485 | 18706 | 0.675 | 0.248 | 0.145 | 0.118 | 0.183 | 4.54 | 3.30 | 2.66 |
| casp3 | 350 | 11172 | 0.554 | 0.158 | 0.007 | 0.023 | 0.061 | 0.86 | 0.80 | 0.97 |
| comt | 86 | 4012 | 0.495 | 0.118 | 0.000 | 0.000 | 0.011 | 0.00 | 0.00 | 0.12 |
| cp2c9 | 183 | 7757 | 0.613 | 0.213 | 0.085 | 0.073 | 0.139 | 3.30 | 2.30 | 2.24 |
| cp3a4 | 363 | 12303 | 0.562 | 0.186 | 0.090 | 0.081 | 0.121 | 3.31 | 1.98 | 1.85 |
| cxcr4 | 122 | 3536 | 0.681 | 0.212 | 0.032 | 0.068 | 0.116 | 2.48 | 1.32 | 1.72 |
| def | 161 | 5899 | 0.382 | 0.090 | 0.000 | 0.003 | 0.016 | 0.00 | 0.37 | 0.19 |
| fa7 | 185 | 6487 | 0.828 | 0.361 | 0.258 | 0.229 | 0.318 | 7.67 | 5.30 | 4.55 |
| fabp4 | 57 | 2912 | 0.818 | 0.534 | 0.718 | 0.564 | 0.508 | 40.52 | 8.46 | 5.44 |
| fak1 | 114 | 5516 | 0.588 | 0.194 | 0.045 | 0.049 | 0.116 | 1.76 | 2.29 | 1.76 |
| fgfr1 | 242 | 736 | 0.550 | 0.165 | 0.377 | 0.277 | 0.374 | 0.43 | 1.10 | 1.33 |
| fkb1a | 273 | 6105 | 0.707 | 0.253 | 0.261 | 0.191 | 0.201 | 5.13 | 3.23 | 2.24 |
| gcr | 563 | 15748 | 0.559 | 0.248 | 0.363 | 0.270 | 0.247 | 10.51 | 3.95 | 2.70 |
| glcm | 313 | 4150 | 0.458 | 0.107 | 0.023 | 0.021 | 0.027 | 0.32 | 0.38 | 0.26 |
| gria2 | 297 | 12358 | 0.728 | 0.319 | 0.193 | 0.165 | 0.292 | 5.75 | 5.73 | 4.51 |
| grik1 | 152 | 6769 | 0.605 | 0.234 | 0.106 | 0.099 | 0.181 | 4.65 | 3.69 | 2.83 |
| hivpr | 1395 | 37673 | 0.704 | 0.247 | 0.086 | 0.096 | 0.171 | 2.44 | 2.50 | 2.47 |
| hs90a | 125 | 5067 | 0.317 | 0.064 | 0.000 | 0.000 | 0.002 | 0.00 | 0.00 | 0.00 |
| hxk4 | 127 | 4930 | 0.533 | 0.140 | 0.088 | 0.034 | 0.030 | 1.58 | 0.32 | 0.24 |
| inha | 71 | 2389 | 0.618 | 0.254 | 0.076 | 0.137 | 0.247 | 2.93 | 4.81 | 4.10 |
| jak2 | 153 | 6743 | 0.755 | 0.349 | 0.509 | 0.283 | 0.303 | 13.81 | 5.49 | 3.73 |
| kif11 | 197 | 7109 | 0.820 | 0.400 | 0.514 | 0.327 | 0.378 | 13.21 | 6.20 | 5.08 |
| kith | 132 | 2998 | 0.421 | 0.204 | 0.740 | 0.357 | 0.211 | 12.53 | 2.90 | 1.90 |
| mapk2 | 206 | 6450 | 0.773 | 0.324 | 0.226 | 0.209 | 0.289 | 5.87 | 5.06 | 3.74 |
| mcr | 193 | 5433 | 0.565 | 0.228 | 0.535 | 0.231 | 0.186 | 7.82 | 2.60 | 2.07 |
| mk01 | 139 | 4767 | 0.818 | 0.325 | 0.004 | 0.057 | 0.253 | 0.00 | 3.75 | 4.90 |
| mk10 | 186 | 6900 | 0.688 | 0.239 | 0.053 | 0.093 | 0.162 | 3.23 | 2.80 | 2.42 |
| nos1 | 234 | 8307 | 0.575 | 0.175 | 0.025 | 0.037 | 0.081 | 1.28 | 1.11 | 1.33 |
| pa2ga | 127 | 5343 | 0.505 | 0.127 | 0.000 | 0.001 | 0.020 | 0.00 | 0.16 | 0.24 |
| plk1 | 155 | 7034 | 0.601 | 0.199 | 0.050 | 0.058 | 0.114 | 2.59 | 1.81 | 1.94 |
| pnph | 233 | 7249 | 0.874 | 0.410 | 0.400 | 0.295 | 0.389 | 9.94 | 6.53 | 5.07 |
| ppard | 288 | 13520 | 0.636 | 0.209 | 0.022 | 0.043 | 0.113 | 1.74 | 2.01 | 1.81 |
| ptn1 | 225 | 7658 | 0.801 | 0.436 | 0.704 | 0.472 | 0.436 | 20.15 | 7.57 | 4.76 |
| pur2 | 201 | 2926 | 0.803 | 0.306 | 0.056 | 0.168 | 0.301 | 1.00 | 4.19 | 3.49 |
| pygm | 114 | 4159 | 0.467 | 0.121 | 0.000 | 0.009 | 0.032 | 0.00 | 0.35 | 0.44 |
| pyrd | 134 | 6782 | 0.698 | 0.248 | 0.162 | 0.108 | 0.158 | 4.53 | 3.14 | 2.24 |
| rock1 | 203 | 6580 | 0.682 | 0.219 | 0.116 | 0.084 | 0.121 | 3.49 | 1.67 | 1.82 |
| rxra | 162 | 7869 | 0.815 | 0.508 | 0.740 | 0.516 | 0.520 | 29.27 | 9.76 | 5.99 |
| sahh | 190 | 3673 | 0.514 | 0.136 | 0.042 | 0.026 | 0.046 | 0.54 | 0.32 | 0.63 |
| thb | 168 | 7821 | 0.819 | 0.489 | 0.679 | 0.495 | 0.506 | 25.07 | 9.29 | 5.95 |
| tryb1 | 171 | 7884 | 0.646 | 0.203 | 0.024 | 0.040 | 0.102 | 1.18 | 1.52 | 1.76 |
| tysy | 311 | 7194 | 0.756 | 0.304 | 0.155 | 0.217 | 0.281 | 5.54 | 3.93 | 3.51 |
| wee1 | 137 | 6371 | 0.867 | 0.548 | 0.871 | 0.572 | 0.535 | 33.96 | 9.65 | 5.99 |
| xiap | 129 | 5342 | 0.701 | 0.294 | 0.176 | 0.211 | 0.264 | 10.94 | 4.96 | 3.33 |

Supplementary Table 4. The virtual screening results by rf_score scorinig function of 51 targets based on the protein structures in DUD-E databases.

| Target | Active | Total | AUC | logAUC | BEDROC(α=321.9) | BEDROC(α=80.5) | BEDROC(α=20.0) | EF1% | EF5% | EF10% |
| --- | --- | --- | --- | --- | --- | --- | --- | --- | --- | --- |
| ada | 262 | 5734 | 0.450 | 0.129 | 0.039 | 0.057 | 0.075 | 1.15 | 0.99 | 0.92 |
| adrb1 | 458 | 16416 | 0.591 | 0.205 | 0.113 | 0.090 | 0.141 | 3.06 | 2.49 | 2.10 |
| adrb2 | 447 | 15702 | 0.618 | 0.216 | 0.049 | 0.082 | 0.156 | 2.24 | 2.86 | 2.28 |
| akt1 | 423 | 16999 | 0.700 | 0.229 | 0.035 | 0.056 | 0.117 | 1.90 | 1.66 | 1.75 |
| akt2 | 190 | 7142 | 0.773 | 0.299 | 0.104 | 0.117 | 0.218 | 4.76 | 3.26 | 3.26 |
| ampc | 62 | 2964 | 0.485 | 0.144 | 0.075 | 0.049 | 0.079 | 1.65 | 1.94 | 1.13 |
| aofb | 168 | 7099 | 0.508 | 0.149 | 0.012 | 0.029 | 0.069 | 0.60 | 1.07 | 1.19 |
| bace1 | 485 | 18706 | 0.694 | 0.259 | 0.087 | 0.119 | 0.200 | 3.71 | 3.67 | 2.87 |
| casp3 | 350 | 11172 | 0.471 | 0.121 | 0.015 | 0.014 | 0.036 | 0.29 | 0.34 | 0.57 |
| comt | 86 | 4012 | 0.402 | 0.083 | 0.000 | 0.000 | 0.001 | 0.00 | 0.00 | 0.00 |
| cp2c9 | 183 | 7757 | 0.590 | 0.219 | 0.165 | 0.131 | 0.168 | 5.50 | 3.40 | 2.08 |
| cp3a4 | 363 | 12303 | 0.609 | 0.237 | 0.157 | 0.148 | 0.205 | 5.51 | 3.75 | 2.76 |
| cxcr4 | 122 | 3536 | 0.845 | 0.309 | 0.004 | 0.039 | 0.184 | 0.00 | 1.48 | 3.20 |
| def | 161 | 5899 | 0.367 | 0.091 | 0.000 | 0.000 | 0.015 | 0.00 | 0.00 | 0.25 |
| fa7 | 185 | 6487 | 0.827 | 0.360 | 0.108 | 0.212 | 0.339 | 6.03 | 6.28 | 4.87 |
| fabp4 | 57 | 2912 | 0.640 | 0.310 | 0.333 | 0.288 | 0.312 | 12.33 | 5.99 | 3.51 |
| fak1 | 114 | 5516 | 0.766 | 0.330 | 0.171 | 0.190 | 0.289 | 7.92 | 5.81 | 4.04 |
| fgfr1 | 242 | 736 | 0.510 | 0.145 | 0.337 | 0.366 | 0.316 | 1.30 | 1.01 | 0.87 |
| fkb1a | 273 | 6105 | 0.618 | 0.192 | 0.047 | 0.051 | 0.111 | 0.73 | 1.39 | 1.65 |
| gcr | 563 | 15748 | 0.674 | 0.221 | 0.092 | 0.074 | 0.127 | 2.14 | 1.85 | 1.67 |
| glcm | 313 | 4150 | 0.537 | 0.158 | 0.001 | 0.015 | 0.077 | 0.00 | 0.32 | 0.83 |
| gria2 | 297 | 12358 | 0.625 | 0.200 | 0.059 | 0.061 | 0.103 | 3.04 | 1.62 | 1.55 |
| grik1 | 152 | 6769 | 0.521 | 0.160 | 0.036 | 0.049 | 0.085 | 1.99 | 1.32 | 1.45 |
| hivpr | 1395 | 37673 | 0.737 | 0.363 | 0.602 | 0.415 | 0.384 | 13.57 | 6.54 | 4.19 |
| hs90a | 125 | 5067 | 0.509 | 0.193 | 0.244 | 0.151 | 0.159 | 6.49 | 3.20 | 1.92 |
| hxk4 | 127 | 4930 | 0.575 | 0.158 | 0.001 | 0.009 | 0.045 | 0.00 | 0.47 | 0.79 |
| inha | 71 | 2389 | 0.565 | 0.157 | 0.035 | 0.043 | 0.053 | 2.93 | 0.57 | 0.71 |
| jak2 | 153 | 6743 | 0.800 | 0.357 | 0.150 | 0.188 | 0.331 | 6.58 | 6.41 | 5.03 |
| kif11 | 197 | 7109 | 0.758 | 0.335 | 0.268 | 0.219 | 0.307 | 8.64 | 5.39 | 4.63 |
| kith | 132 | 2998 | 0.350 | 0.112 | 0.000 | 0.019 | 0.076 | 0.00 | 1.22 | 0.99 |
| mapk2 | 206 | 6450 | 0.687 | 0.263 | 0.204 | 0.180 | 0.223 | 6.36 | 4.08 | 2.67 |
| mcr | 193 | 5433 | 0.721 | 0.254 | 0.103 | 0.138 | 0.179 | 4.17 | 2.29 | 2.38 |
| mk01 | 139 | 4767 | 0.737 | 0.249 | 0.001 | 0.026 | 0.135 | 0.00 | 1.87 | 2.52 |
| mk10 | 186 | 6900 | 0.751 | 0.332 | 0.144 | 0.195 | 0.322 | 5.38 | 6.24 | 4.52 |
| nos1 | 234 | 8307 | 0.593 | 0.208 | 0.102 | 0.079 | 0.144 | 2.99 | 2.40 | 2.44 |
| pa2ga | 127 | 5343 | 0.629 | 0.241 | 0.112 | 0.094 | 0.184 | 3.18 | 2.52 | 3.31 |
| plk1 | 155 | 7034 | 0.703 | 0.278 | 0.095 | 0.127 | 0.225 | 6.48 | 4.01 | 3.42 |
| pnph | 233 | 7249 | 0.640 | 0.230 | 0.121 | 0.138 | 0.186 | 3.46 | 3.18 | 2.36 |
| ppard | 288 | 13520 | 0.622 | 0.252 | 0.208 | 0.159 | 0.209 | 8.00 | 4.17 | 2.71 |
| ptn1 | 225 | 7658 | 0.749 | 0.403 | 0.599 | 0.450 | 0.419 | 17.91 | 7.31 | 4.54 |
| pur2 | 201 | 2926 | 0.661 | 0.171 | 0.000 | 0.001 | 0.029 | 0.00 | 0.00 | 0.20 |
| pygm | 114 | 4159 | 0.370 | 0.088 | 0.064 | 0.021 | 0.021 | 0.89 | 0.18 | 0.26 |
| pyrd | 134 | 6782 | 0.575 | 0.176 | 0.019 | 0.045 | 0.094 | 2.27 | 1.64 | 1.49 |
| rock1 | 203 | 6580 | 0.707 | 0.261 | 0.097 | 0.142 | 0.214 | 3.49 | 3.94 | 2.96 |
| rxra | 162 | 7869 | 0.683 | 0.247 | 0.091 | 0.091 | 0.173 | 4.36 | 3.71 | 2.60 |
| sahh | 190 | 3673 | 0.521 | 0.134 | 0.000 | 0.001 | 0.032 | 0.00 | 0.11 | 0.42 |
| thb | 168 | 7821 | 0.662 | 0.295 | 0.314 | 0.225 | 0.265 | 11.34 | 4.88 | 3.39 |
| tryb1 | 171 | 7884 | 0.684 | 0.232 | 0.023 | 0.049 | 0.137 | 1.77 | 2.22 | 2.52 |
| tysy | 311 | 7194 | 0.542 | 0.184 | 0.065 | 0.119 | 0.160 | 2.61 | 2.64 | 1.93 |
| wee1 | 137 | 6371 | 0.790 | 0.364 | 0.209 | 0.216 | 0.344 | 11.07 | 7.02 | 5.11 |
| xiap | 129 | 5342 | 0.733 | 0.308 | 0.098 | 0.145 | 0.274 | 6.25 | 5.43 | 4.34 |

Supplementary Table 5. The virtual screening results by vinardo scorinig function of 51 targets based on the protein structures in DUD-E databases.

| Target | Active | Total | AUC | logAUC | BEDROC(α=321.9) | BEDROC(α=80.5) | BEDROC(α=20.0) | EF1% | EF5% | EF10% |
| --- | --- | --- | --- | --- | --- | --- | --- | --- | --- | --- |
| ada | 262 | 5734 | 0.445 | 0.110 | 0.031 | 0.024 | 0.037 | 0.77 | 0.54 | 0.53 |
| adrb1 | 458 | 16416 | 0.630 | 0.228 | 0.105 | 0.110 | 0.171 | 3.72 | 2.80 | 2.32 |
| adrb2 | 447 | 15702 | 0.679 | 0.249 | 0.079 | 0.104 | 0.186 | 3.13 | 3.31 | 2.62 |
| akt1 | 423 | 16999 | 0.499 | 0.146 | 0.055 | 0.038 | 0.067 | 1.66 | 0.99 | 0.97 |
| akt2 | 190 | 7142 | 0.683 | 0.223 | 0.014 | 0.054 | 0.121 | 1.59 | 1.90 | 1.58 |
| ampc | 62 | 2964 | 0.458 | 0.112 | 0.000 | 0.006 | 0.026 | 0.00 | 0.32 | 0.32 |
| aofb | 168 | 7099 | 0.720 | 0.267 | 0.093 | 0.103 | 0.192 | 4.23 | 3.10 | 3.16 |
| bace1 | 485 | 18706 | 0.680 | 0.248 | 0.083 | 0.096 | 0.181 | 2.68 | 3.34 | 2.85 |
| casp3 | 350 | 11172 | 0.569 | 0.175 | 0.030 | 0.043 | 0.090 | 1.44 | 1.32 | 1.31 |
| comt | 86 | 4012 | 0.530 | 0.153 | 0.011 | 0.039 | 0.076 | 1.17 | 1.40 | 1.16 |
| cp2c9 | 183 | 7757 | 0.609 | 0.222 | 0.128 | 0.119 | 0.168 | 4.95 | 2.96 | 2.46 |
| cp3a4 | 363 | 12303 | 0.521 | 0.148 | 0.011 | 0.031 | 0.063 | 1.10 | 0.88 | 0.91 |
| cxcr4 | 122 | 3536 | 0.680 | 0.205 | 0.002 | 0.031 | 0.086 | 0.00 | 1.32 | 1.07 |
| def | 161 | 5899 | 0.636 | 0.217 | 0.026 | 0.061 | 0.130 | 2.53 | 2.24 | 1.80 |
| fa7 | 185 | 6487 | 0.801 | 0.348 | 0.294 | 0.207 | 0.307 | 6.57 | 5.09 | 4.49 |
| fabp4 | 57 | 2912 | 0.875 | 0.498 | 0.395 | 0.413 | 0.538 | 19.38 | 10.57 | 7.20 |
| fak1 | 114 | 5516 | 0.631 | 0.222 | 0.057 | 0.068 | 0.144 | 0.88 | 2.64 | 2.20 |
| fgfr1 | 242 | 736 | 0.538 | 0.165 | 0.455 | 0.454 | 0.404 | 1.74 | 1.27 | 1.21 |
| fkb1a | 273 | 6105 | 0.629 | 0.192 | 0.015 | 0.046 | 0.104 | 1.10 | 1.32 | 1.54 |
| gcr | 563 | 15748 | 0.446 | 0.182 | 0.274 | 0.197 | 0.186 | 5.70 | 3.13 | 2.03 |
| glcm | 313 | 4150 | 0.652 | 0.192 | 0.000 | 0.018 | 0.088 | 0.00 | 0.64 | 1.05 |
| gria2 | 297 | 12358 | 0.731 | 0.282 | 0.205 | 0.121 | 0.195 | 4.74 | 2.83 | 3.13 |
| grik1 | 152 | 6769 | 0.723 | 0.313 | 0.202 | 0.216 | 0.274 | 11.30 | 5.01 | 3.49 |
| hivpr | 1395 | 37673 | 0.659 | 0.219 | 0.088 | 0.081 | 0.138 | 2.23 | 2.01 | 1.90 |
| hs90a | 125 | 5067 | 0.322 | 0.063 | 0.000 | 0.000 | 0.002 | 0.00 | 0.00 | 0.08 |
| hxk4 | 127 | 4930 | 0.580 | 0.147 | 0.000 | 0.000 | 0.023 | 0.00 | 0.00 | 0.39 |
| inha | 71 | 2389 | 0.744 | 0.311 | 0.161 | 0.202 | 0.285 | 7.31 | 5.37 | 3.68 |
| jak2 | 153 | 6743 | 0.701 | 0.275 | 0.094 | 0.145 | 0.227 | 5.26 | 4.58 | 3.07 |
| kif11 | 197 | 7109 | 0.774 | 0.420 | 0.677 | 0.456 | 0.427 | 19.82 | 7.32 | 4.83 |
| kith | 132 | 2998 | 0.447 | 0.137 | 0.014 | 0.040 | 0.089 | 0.78 | 1.22 | 1.29 |
| mapk2 | 206 | 6450 | 0.759 | 0.324 | 0.371 | 0.270 | 0.291 | 9.78 | 4.86 | 3.54 |
| mcr | 193 | 5433 | 0.488 | 0.147 | 0.214 | 0.101 | 0.078 | 3.65 | 1.25 | 0.73 |
| mk01 | 139 | 4767 | 0.709 | 0.251 | 0.003 | 0.034 | 0.160 | 0.00 | 3.31 | 2.95 |
| mk10 | 186 | 6900 | 0.683 | 0.241 | 0.044 | 0.086 | 0.165 | 2.69 | 2.69 | 2.42 |
| nos1 | 234 | 8307 | 0.582 | 0.179 | 0.062 | 0.051 | 0.082 | 1.28 | 1.20 | 1.15 |
| pa2ga | 127 | 5343 | 0.650 | 0.290 | 0.290 | 0.232 | 0.286 | 10.32 | 5.99 | 3.62 |
| plk1 | 155 | 7034 | 0.563 | 0.165 | 0.002 | 0.019 | 0.074 | 0.00 | 0.91 | 1.42 |
| pnph | 233 | 7249 | 0.788 | 0.360 | 0.326 | 0.299 | 0.360 | 9.94 | 6.45 | 4.43 |
| ppard | 288 | 13520 | 0.636 | 0.222 | 0.029 | 0.049 | 0.139 | 1.74 | 2.29 | 2.50 |
| ptn1 | 225 | 7658 | 0.806 | 0.431 | 0.522 | 0.440 | 0.459 | 15.67 | 8.46 | 5.25 |
| pur2 | 201 | 2926 | 0.742 | 0.333 | 0.539 | 0.476 | 0.409 | 7.03 | 5.58 | 3.79 |
| pygm | 114 | 4159 | 0.519 | 0.168 | 0.020 | 0.028 | 0.095 | 0.89 | 0.88 | 1.76 |
| pyrd | 134 | 6782 | 0.733 | 0.267 | 0.029 | 0.076 | 0.176 | 3.02 | 2.84 | 2.91 |
| rock1 | 203 | 6580 | 0.681 | 0.243 | 0.178 | 0.126 | 0.178 | 3.49 | 3.15 | 2.36 |
| rxra | 162 | 7869 | 0.752 | 0.443 | 0.728 | 0.465 | 0.451 | 26.16 | 8.28 | 5.19 |
| sahh | 190 | 3673 | 0.486 | 0.138 | 0.007 | 0.058 | 0.091 | 0.00 | 1.27 | 1.05 |
| thb | 168 | 7821 | 0.828 | 0.479 | 0.565 | 0.452 | 0.519 | 22.68 | 10.83 | 6.01 |
| tryb1 | 171 | 7884 | 0.604 | 0.223 | 0.119 | 0.111 | 0.161 | 5.32 | 2.93 | 2.16 |
| tysy | 311 | 7194 | 0.744 | 0.318 | 0.313 | 0.230 | 0.306 | 4.56 | 4.45 | 4.02 |
| wee1 | 137 | 6371 | 0.668 | 0.338 | 0.537 | 0.353 | 0.322 | 19.19 | 5.85 | 3.29 |
| xiap | 129 | 5342 | 0.575 | 0.258 | 0.286 | 0.256 | 0.253 | 14.06 | 4.50 | 2.79 |

Supplementary Table 6. The virtual screening results by autodock4 scorinig function of 51 targets based on the protein structures of AlphaFold.

| Target | Active | Total | AUC | logAUC | BEDROC(α=321.9) | BEDROC(α=80.5) | BEDROC(α=20.0) | EF1% | EF5% | EF10% |
| --- | --- | --- | --- | --- | --- | --- | --- | --- | --- | --- |
| ada | 262 | 5721 | 0.231 | 0.069 | 0.156 | 0.052 | 0.048 | 1.15 | 0.61 | 0.57 |
| adrb1 | 451 | 16277 | 0.631 | 0.253 | 0.280 | 0.182 | 0.214 | 6.91 | 3.91 | 2.73 |
| adrb2 | 444 | 15601 | 0.704 | 0.300 | 0.354 | 0.245 | 0.264 | 10.36 | 4.59 | 3.18 |
| akt1 | 418 | 16947 | 0.721 | 0.300 | 0.209 | 0.183 | 0.262 | 6.24 | 4.55 | 3.71 |
| akt2 | 189 | 7105 | 0.768 | 0.354 | 0.333 | 0.295 | 0.347 | 12.18 | 6.14 | 4.50 |
| ampc | 60 | 2946 | 0.585 | 0.170 | 0.000 | 0.014 | 0.064 | 0.00 | 1.34 | 1.00 |
| aofb | 168 | 7091 | 0.581 | 0.193 | 0.068 | 0.055 | 0.115 | 1.81 | 1.55 | 2.02 |
| bace1 | 484 | 18593 | 0.756 | 0.362 | 0.416 | 0.315 | 0.355 | 12.46 | 6.70 | 4.15 |
| casp3 | 348 | 11084 | 0.707 | 0.315 | 0.217 | 0.231 | 0.314 | 6.95 | 5.98 | 4.17 |
| comt | 86 | 4000 | 0.386 | 0.087 | 0.000 | 0.000 | 0.005 | 0.00 | 0.00 | 0.12 |
| cp2c9 | 183 | 7724 | 0.575 | 0.211 | 0.203 | 0.139 | 0.150 | 6.03 | 2.73 | 1.75 |
| cp3a4 | 359 | 12223 | 0.496 | 0.140 | 0.047 | 0.029 | 0.057 | 1.12 | 0.72 | 0.92 |
| cxcr4 | 120 | 3520 | 0.716 | 0.274 | 0.252 | 0.163 | 0.205 | 5.03 | 2.83 | 2.83 |
| def | 160 | 5866 | 0.723 | 0.282 | 0.150 | 0.129 | 0.221 | 4.42 | 3.50 | 3.50 |
| fa7 | 183 | 6451 | 0.715 | 0.290 | 0.195 | 0.174 | 0.248 | 7.16 | 4.60 | 3.39 |
| fabp4 | 57 | 2897 | 0.598 | 0.180 | 0.000 | 0.009 | 0.077 | 0.00 | 1.06 | 1.76 |
| fak1 | 114 | 5505 | 0.564 | 0.160 | 0.010 | 0.022 | 0.059 | 0.88 | 0.88 | 0.97 |
| fgfr1 | 242 | 730 | 0.556 | 0.167 | 0.249 | 0.314 | 0.391 | 0.43 | 1.34 | 1.20 |
| fkb1a | 273 | 6094 | 0.778 | 0.347 | 0.449 | 0.360 | 0.351 | 10.05 | 5.43 | 3.92 |
| gcr | 563 | 15706 | 0.482 | 0.156 | 0.076 | 0.083 | 0.115 | 2.49 | 1.95 | 1.51 |
| glcm | 311 | 4133 | 0.562 | 0.174 | 0.079 | 0.123 | 0.149 | 1.62 | 1.81 | 1.54 |
| gria2 | 293 | 12313 | 0.537 | 0.174 | 0.044 | 0.046 | 0.097 | 1.71 | 1.71 | 1.60 |
| grik1 | 152 | 6752 | 0.722 | 0.263 | 0.056 | 0.075 | 0.177 | 3.32 | 2.90 | 3.03 |
| hivpr | 1380 | 37284 | 0.802 | 0.417 | 0.590 | 0.464 | 0.455 | 14.16 | 7.78 | 5.01 |
| hs90a | 125 | 5044 | 0.287 | 0.071 | 0.002 | 0.017 | 0.025 | 0.00 | 0.48 | 0.24 |
| hxk4 | 127 | 4913 | 0.595 | 0.178 | 0.000 | 0.013 | 0.072 | 0.00 | 0.63 | 1.26 |
| inha | 70 | 2375 | 0.723 | 0.313 | 0.212 | 0.225 | 0.301 | 8.85 | 5.75 | 3.72 |
| jak2 | 153 | 6715 | 0.547 | 0.179 | 0.114 | 0.071 | 0.108 | 3.28 | 2.23 | 1.50 |
| kif11 | 197 | 7075 | 0.592 | 0.182 | 0.004 | 0.019 | 0.084 | 0.51 | 1.22 | 1.37 |
| kith | 130 | 2981 | 0.662 | 0.266 | 0.239 | 0.178 | 0.264 | 3.16 | 4.00 | 3.62 |
| mapk2 | 205 | 6420 | 0.744 | 0.286 | 0.144 | 0.143 | 0.238 | 4.40 | 3.80 | 3.85 |
| mcr | 193 | 5420 | 0.594 | 0.254 | 0.544 | 0.303 | 0.247 | 10.92 | 4.25 | 2.54 |
| mk01 | 139 | 4742 | 0.673 | 0.203 | 0.006 | 0.024 | 0.066 | 0.73 | 0.72 | 0.86 |
| mk10 | 183 | 6861 | 0.674 | 0.247 | 0.133 | 0.113 | 0.181 | 3.31 | 3.06 | 2.68 |
| nos1 | 232 | 8275 | 0.523 | 0.156 | 0.002 | 0.012 | 0.061 | 0.43 | 0.78 | 0.95 |
| pa2ga | 127 | 5329 | 0.487 | 0.156 | 0.093 | 0.080 | 0.092 | 3.96 | 1.42 | 1.03 |
| plk1 | 153 | 6996 | 0.553 | 0.181 | 0.013 | 0.035 | 0.107 | 0.66 | 1.57 | 1.90 |
| pnph | 232 | 7226 | 0.495 | 0.162 | 0.023 | 0.052 | 0.112 | 0.87 | 2.16 | 1.64 |
| ppard | 287 | 13432 | 0.637 | 0.240 | 0.119 | 0.112 | 0.180 | 4.89 | 3.35 | 2.65 |
| ptn1 | 222 | 7597 | 0.850 | 0.455 | 0.687 | 0.432 | 0.454 | 18.71 | 7.86 | 5.86 |
| pur2 | 199 | 2903 | 0.645 | 0.248 | 0.019 | 0.119 | 0.258 | 1.01 | 3.22 | 3.52 |
| pygm | 114 | 4142 | 0.592 | 0.194 | 0.127 | 0.110 | 0.127 | 5.32 | 2.28 | 1.58 |
| pyrd | 134 | 6768 | 0.619 | 0.201 | 0.015 | 0.043 | 0.107 | 2.26 | 1.34 | 1.72 |
| rock1 | 203 | 6552 | 0.661 | 0.237 | 0.421 | 0.187 | 0.153 | 7.45 | 2.47 | 1.43 |
| rxra | 162 | 7852 | 0.814 | 0.431 | 0.379 | 0.316 | 0.439 | 14.91 | 9.03 | 6.05 |
| sahh | 190 | 3661 | 0.192 | 0.043 | 0.000 | 0.000 | 0.011 | 0.00 | 0.00 | 0.26 |
| thb | 167 | 7778 | 0.744 | 0.271 | 0.006 | 0.056 | 0.186 | 1.21 | 3.12 | 3.18 |
| tryb1 | 170 | 7841 | 0.766 | 0.413 | 0.585 | 0.401 | 0.417 | 19.51 | 8.35 | 4.65 |
| tysy | 310 | 7145 | 0.622 | 0.244 | 0.243 | 0.223 | 0.232 | 5.84 | 3.29 | 2.68 |
| wee1 | 137 | 6329 | 0.896 | 0.599 | 0.787 | 0.618 | 0.637 | 30.80 | 12.43 | 7.24 |
| xiap | 128 | 5324 | 0.626 | 0.196 | 0.001 | 0.024 | 0.098 | 0.00 | 1.72 | 1.56 |

Supplementary Table 7. The virtual screening results by idock scorinig function of 51 targets based on the protein structures of AlphaFold.

| Target | Active | Total | AUC | logAUC | BEDROC(α=321.9) | BEDROC(α=80.5) | BEDROC(α=20.0) | EF1% | EF5% | EF10% |
| --- | --- | --- | --- | --- | --- | --- | --- | --- | --- | --- |
| ada | 262 | 5734 | 0.452 | 0.122 | 0.052 | 0.045 | 0.061 | 0.77 | 1.07 | 0.76 |
| adrb1 | 458 | 16416 | 0.626 | 0.202 | 0.052 | 0.064 | 0.118 | 2.19 | 1.79 | 1.79 |
| adrb2 | 447 | 15702 | 0.643 | 0.218 | 0.082 | 0.078 | 0.137 | 2.46 | 2.28 | 2.01 |
| akt1 | 423 | 16999 | 0.712 | 0.280 | 0.147 | 0.165 | 0.230 | 6.66 | 4.31 | 3.05 |
| akt2 | 190 | 7142 | 0.749 | 0.321 | 0.141 | 0.206 | 0.305 | 6.35 | 5.58 | 4.37 |
| ampc | 62 | 2964 | 0.474 | 0.118 | 0.000 | 0.001 | 0.025 | 0.00 | 0.00 | 0.65 |
| aofb | 168 | 7099 | 0.693 | 0.266 | 0.109 | 0.137 | 0.210 | 4.83 | 4.06 | 2.98 |
| bace1 | 485 | 18706 | 0.681 | 0.229 | 0.043 | 0.055 | 0.135 | 1.86 | 2.23 | 2.21 |
| casp3 | 350 | 11172 | 0.640 | 0.236 | 0.083 | 0.126 | 0.194 | 3.16 | 3.43 | 2.60 |
| comt | 86 | 4012 | 0.481 | 0.115 | 0.000 | 0.000 | 0.009 | 0.00 | 0.00 | 0.00 |
| cp2c9 | 183 | 7757 | 0.617 | 0.213 | 0.084 | 0.086 | 0.147 | 2.75 | 2.52 | 2.24 |
| cp3a4 | 363 | 12303 | 0.565 | 0.197 | 0.172 | 0.106 | 0.134 | 4.68 | 2.31 | 1.82 |
| cxcr4 | 122 | 3536 | 0.657 | 0.184 | 0.000 | 0.009 | 0.065 | 0.00 | 0.99 | 1.07 |
| def | 161 | 5899 | 0.455 | 0.123 | 0.001 | 0.007 | 0.043 | 0.00 | 0.62 | 0.87 |
| fa7 | 185 | 6487 | 0.733 | 0.282 | 0.191 | 0.147 | 0.225 | 4.38 | 4.22 | 3.35 |
| fabp4 | 57 | 2912 | 0.775 | 0.401 | 0.360 | 0.349 | 0.405 | 19.38 | 8.10 | 4.74 |
| fak1 | 114 | 5516 | 0.631 | 0.192 | 0.086 | 0.051 | 0.081 | 1.76 | 1.23 | 0.97 |
| fgfr1 | 242 | 736 | 0.548 | 0.169 | 0.393 | 0.408 | 0.415 | 0.87 | 1.35 | 1.21 |
| fkb1a | 273 | 6105 | 0.692 | 0.250 | 0.272 | 0.201 | 0.193 | 6.97 | 2.79 | 2.09 |
| gcr | 563 | 15748 | 0.426 | 0.131 | 0.106 | 0.078 | 0.094 | 2.49 | 1.53 | 1.16 |
| glcm | 313 | 4150 | 0.337 | 0.069 | 0.000 | 0.001 | 0.008 | 0.00 | 0.13 | 0.10 |
| gria2 | 297 | 12358 | 0.544 | 0.234 | 0.337 | 0.209 | 0.212 | 9.47 | 3.71 | 2.59 |
| grik1 | 152 | 6769 | 0.717 | 0.257 | 0.045 | 0.061 | 0.153 | 3.99 | 1.58 | 3.03 |
| hivpr | 1395 | 37673 | 0.590 | 0.183 | 0.044 | 0.055 | 0.105 | 1.22 | 1.56 | 1.52 |
| hs90a | 125 | 5067 | 0.353 | 0.080 | 0.000 | 0.000 | 0.008 | 0.00 | 0.00 | 0.24 |
| hxk4 | 127 | 4930 | 0.520 | 0.126 | 0.000 | 0.005 | 0.015 | 0.00 | 0.16 | 0.24 |
| inha | 71 | 2389 | 0.637 | 0.301 | 0.314 | 0.297 | 0.324 | 10.24 | 6.50 | 3.39 |
| jak2 | 153 | 6743 | 0.657 | 0.226 | 0.110 | 0.091 | 0.143 | 3.29 | 2.88 | 1.83 |
| kif11 | 197 | 7109 | 0.737 | 0.280 | 0.102 | 0.134 | 0.217 | 5.08 | 3.96 | 2.95 |
| kith | 132 | 2998 | 0.712 | 0.346 | 0.408 | 0.339 | 0.398 | 10.18 | 5.79 | 5.24 |
| mapk2 | 206 | 6450 | 0.799 | 0.351 | 0.304 | 0.263 | 0.328 | 8.81 | 5.74 | 3.98 |
| mcr | 193 | 5433 | 0.559 | 0.180 | 0.263 | 0.119 | 0.106 | 3.13 | 1.45 | 1.09 |
| mk01 | 139 | 4767 | 0.798 | 0.294 | 0.023 | 0.051 | 0.188 | 0.73 | 2.16 | 3.31 |
| mk10 | 186 | 6900 | 0.674 | 0.216 | 0.047 | 0.055 | 0.116 | 1.61 | 1.61 | 1.61 |
| nos1 | 234 | 8307 | 0.541 | 0.163 | 0.010 | 0.037 | 0.077 | 0.86 | 1.11 | 1.07 |
| pa2ga | 127 | 5343 | 0.449 | 0.107 | 0.000 | 0.000 | 0.016 | 0.00 | 0.00 | 0.32 |
| plk1 | 155 | 7034 | 0.613 | 0.196 | 0.041 | 0.041 | 0.104 | 1.30 | 1.81 | 1.87 |
| pnph | 233 | 7249 | 0.776 | 0.320 | 0.283 | 0.220 | 0.273 | 7.78 | 4.13 | 3.78 |
| ppard | 288 | 13520 | 0.762 | 0.267 | 0.073 | 0.058 | 0.152 | 2.09 | 2.08 | 2.85 |
| ptn1 | 225 | 7658 | 0.756 | 0.376 | 0.532 | 0.375 | 0.368 | 16.57 | 6.06 | 4.27 |
| pur2 | 201 | 2926 | 0.807 | 0.265 | 0.000 | 0.015 | 0.138 | 0.00 | 0.70 | 1.94 |
| pygm | 114 | 4159 | 0.625 | 0.198 | 0.007 | 0.046 | 0.111 | 0.00 | 1.76 | 1.67 |
| pyrd | 134 | 6782 | 0.684 | 0.250 | 0.068 | 0.089 | 0.177 | 3.78 | 3.28 | 2.84 |
| rock1 | 203 | 6580 | 0.703 | 0.228 | 0.076 | 0.060 | 0.127 | 1.50 | 1.97 | 2.07 |
| rxra | 162 | 7869 | 0.819 | 0.482 | 0.619 | 0.448 | 0.500 | 23.66 | 9.52 | 6.24 |
| sahh | 190 | 3673 | 0.440 | 0.115 | 0.000 | 0.009 | 0.045 | 0.00 | 0.53 | 0.63 |
| thb | 168 | 7821 | 0.796 | 0.378 | 0.269 | 0.278 | 0.362 | 14.32 | 6.67 | 4.82 |
| tryb1 | 171 | 7884 | 0.756 | 0.261 | 0.001 | 0.035 | 0.153 | 0.00 | 2.46 | 2.81 |
| tysy | 311 | 7194 | 0.765 | 0.291 | 0.046 | 0.107 | 0.241 | 1.30 | 3.67 | 3.67 |
| wee1 | 137 | 6371 | 0.762 | 0.240 | 0.009 | 0.013 | 0.083 | 0.74 | 0.44 | 1.61 |
| xiap | 129 | 5342 | 0.424 | 0.095 | 0.000 | 0.000 | 0.005 | 0.00 | 0.00 | 0.00 |

Supplementary Table 8. The virtual screening results by rf_score scorinig function of 51 targets based on the protein structures of AlphaFold.

| Target | Active | Total | AUC | logAUC | BEDROC(α=321.9) | BEDROC(α=80.5) | BEDROC(α=20.0) | EF1% | EF5% | EF10% |
| --- | --- | --- | --- | --- | --- | --- | --- | --- | --- | --- |
| ada | 262 | 5734 | 0.485 | 0.146 | 0.091 | 0.089 | 0.099 | 1.92 | 1.61 | 1.15 |
| adrb1 | 458 | 16416 | 0.611 | 0.209 | 0.065 | 0.073 | 0.138 | 1.97 | 2.32 | 2.12 |
| adrb2 | 447 | 15702 | 0.608 | 0.199 | 0.072 | 0.048 | 0.111 | 1.34 | 1.83 | 1.81 |
| akt1 | 423 | 16999 | 0.770 | 0.296 | 0.112 | 0.114 | 0.211 | 4.28 | 3.36 | 3.24 |
| akt2 | 190 | 7142 | 0.804 | 0.321 | 0.082 | 0.128 | 0.250 | 5.29 | 4.00 | 3.74 |
| ampc | 62 | 2964 | 0.460 | 0.116 | 0.000 | 0.002 | 0.026 | 0.00 | 0.32 | 0.48 |
| aofb | 168 | 7099 | 0.586 | 0.181 | 0.037 | 0.044 | 0.092 | 1.21 | 1.67 | 1.43 |
| bace1 | 485 | 18706 | 0.710 | 0.315 | 0.268 | 0.240 | 0.300 | 9.08 | 5.82 | 3.80 |
| casp3 | 350 | 11172 | 0.541 | 0.148 | 0.003 | 0.015 | 0.054 | 0.58 | 0.80 | 0.97 |
| comt | 86 | 4012 | 0.337 | 0.065 | 0.000 | 0.000 | 0.000 | 0.00 | 0.00 | 0.00 |
| cp2c9 | 183 | 7757 | 0.590 | 0.209 | 0.083 | 0.097 | 0.149 | 3.85 | 2.85 | 2.08 |
| cp3a4 | 363 | 12303 | 0.596 | 0.217 | 0.123 | 0.106 | 0.168 | 3.58 | 2.92 | 2.31 |
| cxcr4 | 122 | 3536 | 0.814 | 0.282 | 0.005 | 0.033 | 0.152 | 0.00 | 1.65 | 2.71 |
| def | 161 | 5899 | 0.351 | 0.083 | 0.000 | 0.004 | 0.027 | 0.00 | 0.50 | 0.56 |
| fa7 | 185 | 6487 | 0.732 | 0.308 | 0.234 | 0.242 | 0.280 | 10.96 | 4.87 | 3.57 |
| fabp4 | 57 | 2912 | 0.571 | 0.192 | 0.101 | 0.063 | 0.125 | 3.52 | 2.11 | 2.28 |
| fak1 | 114 | 5516 | 0.795 | 0.328 | 0.155 | 0.164 | 0.262 | 7.92 | 4.57 | 4.22 |
| fgfr1 | 242 | 736 | 0.511 | 0.137 | 0.103 | 0.226 | 0.247 | 0.43 | 0.84 | 0.62 |
| fkb1a | 273 | 6105 | 0.621 | 0.201 | 0.043 | 0.065 | 0.132 | 1.10 | 1.83 | 1.72 |
| gcr | 563 | 15748 | 0.675 | 0.238 | 0.032 | 0.082 | 0.185 | 1.60 | 3.31 | 2.81 |
| glcm | 313 | 4150 | 0.450 | 0.108 | 0.000 | 0.000 | 0.018 | 0.00 | 0.00 | 0.19 |
| gria2 | 297 | 12358 | 0.545 | 0.150 | 0.000 | 0.001 | 0.040 | 0.00 | 0.27 | 0.74 |
| grik1 | 152 | 6769 | 0.498 | 0.134 | 0.000 | 0.002 | 0.044 | 0.00 | 0.26 | 1.05 |
| hivpr | 1395 | 37673 | 0.651 | 0.308 | 0.707 | 0.390 | 0.308 | 13.57 | 4.92 | 3.25 |
| hs90a | 125 | 5067 | 0.481 | 0.144 | 0.052 | 0.044 | 0.064 | 1.62 | 0.96 | 0.80 |
| hxk4 | 127 | 4930 | 0.599 | 0.180 | 0.004 | 0.021 | 0.081 | 0.79 | 0.95 | 1.50 |
| inha | 71 | 2389 | 0.552 | 0.155 | 0.013 | 0.035 | 0.063 | 1.46 | 1.13 | 0.85 |
| jak2 | 153 | 6743 | 0.727 | 0.313 | 0.196 | 0.190 | 0.279 | 7.89 | 5.23 | 3.86 |
| kif11 | 197 | 7109 | 0.653 | 0.204 | 0.001 | 0.021 | 0.089 | 0.00 | 1.22 | 1.27 |
| kith | 132 | 2998 | 0.443 | 0.110 | 0.000 | 0.003 | 0.035 | 0.00 | 0.46 | 0.68 |
| mapk2 | 206 | 6450 | 0.681 | 0.254 | 0.101 | 0.139 | 0.217 | 3.91 | 3.89 | 2.91 |
| mcr | 193 | 5433 | 0.653 | 0.204 | 0.010 | 0.029 | 0.114 | 0.52 | 1.45 | 2.02 |
| mk01 | 139 | 4767 | 0.678 | 0.230 | 0.128 | 0.086 | 0.136 | 2.92 | 2.31 | 1.87 |
| mk10 | 186 | 6900 | 0.746 | 0.323 | 0.169 | 0.171 | 0.297 | 5.38 | 5.59 | 4.68 |
| nos1 | 234 | 8307 | 0.583 | 0.203 | 0.041 | 0.062 | 0.144 | 2.14 | 2.91 | 2.44 |
| pa2ga | 127 | 5343 | 0.409 | 0.113 | 0.023 | 0.035 | 0.051 | 1.59 | 0.63 | 0.63 |
| plk1 | 155 | 7034 | 0.699 | 0.255 | 0.030 | 0.068 | 0.173 | 1.94 | 2.59 | 3.03 |
| pnph | 233 | 7249 | 0.688 | 0.305 | 0.369 | 0.272 | 0.301 | 9.51 | 4.98 | 3.74 |
| ppard | 288 | 13520 | 0.691 | 0.290 | 0.171 | 0.161 | 0.251 | 7.30 | 5.14 | 3.51 |
| ptn1 | 225 | 7658 | 0.696 | 0.324 | 0.414 | 0.312 | 0.319 | 12.99 | 5.61 | 3.65 |
| pur2 | 201 | 2926 | 0.593 | 0.146 | 0.000 | 0.000 | 0.022 | 0.00 | 0.00 | 0.25 |
| pygm | 114 | 4159 | 0.531 | 0.141 | 0.000 | 0.004 | 0.036 | 0.00 | 0.35 | 0.53 |
| pyrd | 134 | 6782 | 0.632 | 0.191 | 0.001 | 0.014 | 0.078 | 0.00 | 0.75 | 1.42 |
| rock1 | 203 | 6580 | 0.730 | 0.282 | 0.182 | 0.183 | 0.234 | 5.98 | 4.33 | 3.05 |
| rxra | 162 | 7869 | 0.636 | 0.211 | 0.047 | 0.072 | 0.124 | 4.36 | 1.98 | 1.98 |
| sahh | 190 | 3673 | 0.447 | 0.112 | 0.000 | 0.001 | 0.030 | 0.00 | 0.00 | 0.58 |
| thb | 168 | 7821 | 0.622 | 0.214 | 0.098 | 0.075 | 0.139 | 2.39 | 2.50 | 2.26 |
| tryb1 | 171 | 7884 | 0.696 | 0.266 | 0.044 | 0.109 | 0.218 | 4.73 | 4.56 | 3.28 |
| tysy | 311 | 7194 | 0.504 | 0.156 | 0.020 | 0.049 | 0.108 | 0.65 | 1.74 | 1.61 |
| wee1 | 137 | 6371 | 0.738 | 0.237 | 0.026 | 0.018 | 0.089 | 0.74 | 0.73 | 1.53 |
| xiap | 129 | 5342 | 0.631 | 0.172 | 0.000 | 0.001 | 0.031 | 0.00 | 0.31 | 0.39 |

Supplementary Table 9. The virtual screening results by vinardo scorinig function of 51 targets based on the protein structures of AlphaFold.

| Target | Active | Total | AUC | logAUC | BEDROC(α=321.9) | BEDROC(α=80.5) | BEDROC(α=20.0) | EF1% | EF5% | EF10% |
| --- | --- | --- | --- | --- | --- | --- | --- | --- | --- | --- |
| ada | 262 | 5734 | 0.483 | 0.125 | 0.011 | 0.016 | 0.039 | 0.38 | 0.54 | 0.57 |
| adrb1 | 458 | 16416 | 0.619 | 0.212 | 0.071 | 0.093 | 0.141 | 3.28 | 2.49 | 1.86 |
| adrb2 | 447 | 15702 | 0.616 | 0.208 | 0.086 | 0.075 | 0.130 | 2.68 | 2.10 | 1.99 |
| akt1 | 423 | 16999 | 0.640 | 0.229 | 0.056 | 0.088 | 0.173 | 3.09 | 3.31 | 2.58 |
| akt2 | 190 | 7142 | 0.723 | 0.260 | 0.036 | 0.067 | 0.180 | 0.53 | 2.84 | 2.79 |
| ampc | 62 | 2964 | 0.565 | 0.165 | 0.000 | 0.006 | 0.059 | 0.00 | 0.32 | 1.45 |
| aofb | 168 | 7099 | 0.712 | 0.287 | 0.213 | 0.175 | 0.233 | 6.64 | 4.54 | 2.92 |
| bace1 | 485 | 18706 | 0.655 | 0.229 | 0.070 | 0.079 | 0.153 | 2.68 | 2.68 | 2.39 |
| casp3 | 350 | 11172 | 0.675 | 0.287 | 0.317 | 0.250 | 0.260 | 10.93 | 4.35 | 3.09 |
| comt | 86 | 4012 | 0.534 | 0.146 | 0.000 | 0.001 | 0.046 | 0.00 | 0.00 | 1.05 |
| cp2c9 | 183 | 7757 | 0.592 | 0.210 | 0.189 | 0.117 | 0.134 | 6.61 | 2.19 | 1.80 |
| cp3a4 | 363 | 12303 | 0.516 | 0.147 | 0.007 | 0.032 | 0.066 | 0.28 | 0.94 | 1.02 |
| cxcr4 | 122 | 3536 | 0.517 | 0.137 | 0.000 | 0.006 | 0.043 | 0.00 | 1.15 | 0.74 |
| def | 161 | 5899 | 0.623 | 0.253 | 0.333 | 0.221 | 0.219 | 8.84 | 4.11 | 2.43 |
| fa7 | 185 | 6487 | 0.738 | 0.285 | 0.117 | 0.122 | 0.230 | 2.74 | 4.33 | 3.35 |
| fabp4 | 57 | 2912 | 0.831 | 0.440 | 0.589 | 0.351 | 0.399 | 19.38 | 6.69 | 5.44 |
| fak1 | 114 | 5516 | 0.496 | 0.162 | 0.064 | 0.065 | 0.101 | 2.64 | 2.29 | 1.32 |
| fgfr1 | 242 | 736 | 0.539 | 0.159 | 0.444 | 0.432 | 0.370 | 1.30 | 1.10 | 1.08 |
| fkb1a | 273 | 6105 | 0.623 | 0.195 | 0.018 | 0.040 | 0.113 | 0.73 | 1.61 | 1.72 |
| gcr | 563 | 15748 | 0.392 | 0.099 | 0.055 | 0.031 | 0.033 | 1.25 | 0.39 | 0.43 |
| glcm | 313 | 4150 | 0.482 | 0.113 | 0.000 | 0.006 | 0.019 | 0.00 | 0.13 | 0.10 |
| gria2 | 297 | 12358 | 0.529 | 0.199 | 0.183 | 0.147 | 0.182 | 5.41 | 3.37 | 2.39 |
| grik1 | 152 | 6769 | 0.660 | 0.308 | 0.396 | 0.254 | 0.291 | 12.63 | 5.53 | 3.69 |
| hivpr | 1395 | 37673 | 0.611 | 0.195 | 0.067 | 0.068 | 0.121 | 1.94 | 1.91 | 1.76 |
| hs90a | 125 | 5067 | 0.318 | 0.073 | 0.005 | 0.016 | 0.017 | 0.81 | 0.32 | 0.16 |
| hxk4 | 127 | 4930 | 0.577 | 0.156 | 0.026 | 0.024 | 0.046 | 1.58 | 0.47 | 0.71 |
| inha | 71 | 2389 | 0.679 | 0.264 | 0.097 | 0.212 | 0.239 | 8.78 | 4.24 | 2.26 |
| jak2 | 153 | 6743 | 0.640 | 0.209 | 0.059 | 0.055 | 0.117 | 2.63 | 1.96 | 1.83 |
| kif11 | 197 | 7109 | 0.645 | 0.215 | 0.027 | 0.061 | 0.141 | 1.02 | 2.13 | 2.39 |
| kith | 132 | 2998 | 0.681 | 0.283 | 0.327 | 0.248 | 0.288 | 7.05 | 4.57 | 3.42 |
| mapk2 | 206 | 6450 | 0.803 | 0.391 | 0.381 | 0.355 | 0.411 | 12.72 | 7.78 | 5.00 |
| mcr | 193 | 5433 | 0.485 | 0.121 | 0.052 | 0.032 | 0.038 | 1.04 | 0.42 | 0.36 |
| mk01 | 139 | 4767 | 0.766 | 0.293 | 0.047 | 0.102 | 0.228 | 2.19 | 4.03 | 3.39 |
| mk10 | 186 | 6900 | 0.718 | 0.244 | 0.015 | 0.046 | 0.141 | 1.08 | 1.94 | 2.31 |
| nos1 | 234 | 8307 | 0.586 | 0.189 | 0.017 | 0.041 | 0.107 | 0.86 | 1.71 | 1.88 |
| pa2ga | 127 | 5343 | 0.530 | 0.143 | 0.009 | 0.024 | 0.048 | 1.59 | 0.47 | 0.71 |
| plk1 | 155 | 7034 | 0.582 | 0.172 | 0.022 | 0.033 | 0.078 | 1.30 | 1.16 | 1.42 |
| pnph | 233 | 7249 | 0.587 | 0.210 | 0.224 | 0.135 | 0.162 | 4.75 | 3.01 | 2.11 |
| ppard | 288 | 13520 | 0.778 | 0.301 | 0.051 | 0.091 | 0.220 | 4.17 | 3.68 | 3.85 |
| ptn1 | 225 | 7658 | 0.756 | 0.356 | 0.582 | 0.322 | 0.330 | 13.44 | 5.43 | 3.96 |
| pur2 | 201 | 2926 | 0.766 | 0.276 | 0.015 | 0.055 | 0.207 | 0.50 | 1.99 | 2.79 |
| pygm | 114 | 4159 | 0.545 | 0.181 | 0.044 | 0.043 | 0.114 | 0.89 | 2.11 | 1.93 |
| pyrd | 134 | 6782 | 0.635 | 0.208 | 0.051 | 0.063 | 0.118 | 3.02 | 1.94 | 1.87 |
| rock1 | 203 | 6580 | 0.614 | 0.174 | 0.000 | 0.010 | 0.061 | 0.00 | 0.79 | 0.99 |
| rxra | 162 | 7869 | 0.713 | 0.352 | 0.543 | 0.317 | 0.333 | 17.44 | 6.18 | 4.14 |
| sahh | 190 | 3673 | 0.369 | 0.085 | 0.001 | 0.012 | 0.023 | 0.00 | 0.21 | 0.32 |
| thb | 168 | 7821 | 0.731 | 0.333 | 0.322 | 0.226 | 0.304 | 10.15 | 5.83 | 4.17 |
| tryb1 | 171 | 7884 | 0.656 | 0.226 | 0.056 | 0.070 | 0.136 | 2.96 | 2.11 | 1.99 |
| tysy | 311 | 7194 | 0.764 | 0.318 | 0.227 | 0.218 | 0.307 | 4.56 | 4.90 | 4.02 |
| wee1 | 137 | 6371 | 0.607 | 0.160 | 0.000 | 0.001 | 0.031 | 0.00 | 0.15 | 0.66 |
| xiap | 129 | 5342 | 0.419 | 0.107 | 0.000 | 0.005 | 0.034 | 0.00 | 0.62 | 0.78 |

Supplementary Table 10. The virtual screening results by autodock4 scorinig function of 23 targets based on the Apo protein structures.

| Target | Active | Total | AUC | logAUC | BEDROC(α=321.9) | BEDROC(α=80.5) | BEDROC(α=20.0) | EF1% | EF5% | EF10% |
| --- | --- | --- | --- | --- | --- | --- | --- | --- | --- | --- |
| ada | 262 | 5707 | 0.214 | 0.047 | 0.000 | 0.004 | 0.015 | 0.00 | 0.23 | 0.19 |
| ampc | 62 | 2950 | 0.597 | 0.185 | 0.032 | 0.038 | 0.093 | 1.64 | 1.29 | 1.61 |
| bace1 | 480 | 18576 | 0.614 | 0.208 | 0.100 | 0.079 | 0.129 | 2.93 | 2.09 | 1.94 |
| comt | 86 | 4005 | 0.482 | 0.113 | 0.000 | 0.000 | 0.008 | 0.00 | 0.00 | 0.12 |
| cp2c9 | 181 | 7711 | 0.555 | 0.209 | 0.230 | 0.130 | 0.160 | 4.43 | 2.55 | 2.15 |
| def | 161 | 5862 | 0.643 | 0.231 | 0.168 | 0.096 | 0.152 | 3.14 | 2.24 | 2.24 |
| fabp4 | 57 | 2896 | 0.826 | 0.455 | 0.434 | 0.415 | 0.462 | 27.22 | 8.82 | 5.45 |
| fgfr1 | 241 | 732 | 0.532 | 0.164 | 0.436 | 0.439 | 0.405 | 1.30 | 1.35 | 1.17 |
| fkb1a | 270 | 6082 | 0.784 | 0.335 | 0.348 | 0.291 | 0.333 | 6.76 | 5.34 | 3.93 |
| gria2 | 296 | 12305 | 0.602 | 0.183 | 0.007 | 0.024 | 0.080 | 0.68 | 1.08 | 1.39 |
| hs90a | 125 | 5057 | 0.319 | 0.069 | 0.000 | 0.000 | 0.006 | 0.00 | 0.00 | 0.16 |
| hxk4 | 127 | 4886 | 0.690 | 0.262 | 0.077 | 0.111 | 0.211 | 3.21 | 3.94 | 3.15 |
| jak2 | 151 | 6717 | 0.403 | 0.095 | 0.000 | 0.002 | 0.020 | 0.00 | 0.27 | 0.40 |
| mapk2 | 206 | 6419 | 0.570 | 0.164 | 0.010 | 0.025 | 0.068 | 0.49 | 0.78 | 1.17 |
| mk01 | 137 | 4749 | 0.723 | 0.261 | 0.055 | 0.090 | 0.168 | 4.43 | 2.19 | 2.63 |
| mk10 | 185 | 6879 | 0.572 | 0.166 | 0.027 | 0.025 | 0.074 | 0.55 | 1.19 | 1.19 |
| nos1 | 232 | 8274 | 0.536 | 0.169 | 0.013 | 0.039 | 0.088 | 1.30 | 1.21 | 1.38 |
| pa2ga | 127 | 5302 | 0.464 | 0.126 | 0.033 | 0.029 | 0.053 | 1.58 | 0.79 | 0.87 |
| pnph | 233 | 7220 | 0.478 | 0.126 | 0.003 | 0.016 | 0.041 | 0.43 | 0.69 | 0.60 |
| ppard | 287 | 13448 | 0.696 | 0.298 | 0.210 | 0.194 | 0.262 | 7.34 | 5.16 | 3.31 |
| ptn1 | 225 | 7626 | 0.760 | 0.300 | 0.118 | 0.141 | 0.249 | 4.46 | 4.18 | 3.83 |
| pur2 | 199 | 2910 | 0.634 | 0.298 | 0.400 | 0.384 | 0.383 | 6.56 | 4.44 | 4.12 |
| tysy | 311 | 7140 | 0.642 | 0.272 | 0.141 | 0.231 | 0.303 | 4.20 | 5.27 | 3.67 |

Supplementary Table 11. The virtual screening results by idock scorinig function of 23 targets based on the Apo protein structures.

| Target | Active | Total | AUC | logAUC | BEDROC(α=321.9) | BEDROC(α=80.5) | BEDROC(α=20.0) | EF1% | EF5% | EF10% |
| --- | --- | --- | --- | --- | --- | --- | --- | --- | --- | --- |
| ada | 262 | 5734 | 0.426 | 0.105 | 0.001 | 0.007 | 0.028 | 0.00 | 0.23 | 0.34 |
| ampc | 62 | 2964 | 0.467 | 0.125 | 0.000 | 0.006 | 0.045 | 0.00 | 1.29 | 0.65 |
| bace1 | 485 | 18706 | 0.580 | 0.186 | 0.010 | 0.047 | 0.115 | 1.24 | 2.02 | 1.82 |
| comt | 86 | 4012 | 0.475 | 0.108 | 0.000 | 0.000 | 0.009 | 0.00 | 0.00 | 0.12 |
| cp2c9 | 183 | 7757 | 0.600 | 0.189 | 0.006 | 0.025 | 0.096 | 1.10 | 1.42 | 1.75 |
| def | 161 | 5899 | 0.311 | 0.072 | 0.000 | 0.000 | 0.011 | 0.00 | 0.00 | 0.31 |
| fabp4 | 57 | 2912 | 0.847 | 0.482 | 0.319 | 0.402 | 0.537 | 17.62 | 10.92 | 6.85 |
| fgfr1 | 242 | 736 | 0.584 | 0.179 | 0.711 | 0.479 | 0.400 | 1.74 | 1.27 | 1.04 |
| fkb1a | 273 | 6105 | 0.664 | 0.208 | 0.012 | 0.058 | 0.130 | 0.37 | 2.35 | 1.65 |
| gria2 | 297 | 12358 | 0.655 | 0.225 | 0.041 | 0.055 | 0.135 | 1.69 | 2.02 | 2.29 |
| hs90a | 125 | 5067 | 0.375 | 0.087 | 0.000 | 0.001 | 0.012 | 0.00 | 0.16 | 0.24 |
| hxk4 | 127 | 4930 | 0.591 | 0.191 | 0.067 | 0.053 | 0.103 | 1.58 | 1.74 | 1.26 |
| jak2 | 153 | 6743 | 0.618 | 0.190 | 0.100 | 0.069 | 0.087 | 3.29 | 1.31 | 1.05 |
| mapk2 | 206 | 6450 | 0.774 | 0.311 | 0.199 | 0.213 | 0.266 | 6.36 | 4.57 | 3.30 |
| mk01 | 139 | 4767 | 0.804 | 0.342 | 0.193 | 0.153 | 0.302 | 4.38 | 5.19 | 5.19 |
| mk10 | 186 | 6900 | 0.631 | 0.199 | 0.063 | 0.060 | 0.109 | 2.69 | 1.83 | 1.83 |
| nos1 | 234 | 8307 | 0.593 | 0.189 | 0.045 | 0.041 | 0.099 | 1.28 | 1.28 | 1.75 |
| pa2ga | 127 | 5343 | 0.351 | 0.072 | 0.000 | 0.000 | 0.004 | 0.00 | 0.00 | 0.08 |
| pnph | 233 | 7249 | 0.666 | 0.196 | 0.029 | 0.052 | 0.079 | 1.30 | 1.12 | 0.95 |
| ppard | 288 | 13520 | 0.785 | 0.294 | 0.120 | 0.102 | 0.195 | 4.17 | 3.54 | 3.02 |
| ptn1 | 225 | 7658 | 0.658 | 0.217 | 0.079 | 0.062 | 0.121 | 1.34 | 1.78 | 1.78 |
| pur2 | 201 | 2926 | 0.659 | 0.176 | 0.000 | 0.003 | 0.047 | 0.00 | 0.10 | 0.60 |
| tysy | 311 | 7194 | 0.764 | 0.298 | 0.137 | 0.149 | 0.259 | 2.93 | 4.25 | 3.47 |

Supplementary Table 12. The virtual screening results by rf_score scorinig function of 23 targets based on the Apo protein structures.

| Target | Active | Total | AUC | logAUC | BEDROC(α=321.9) | BEDROC(α=80.5) | BEDROC(α=20.0) | EF1% | EF5% | EF10% |
| --- | --- | --- | --- | --- | --- | --- | --- | --- | --- | --- |
| ada | 262 | 5734 | 0.466 | 0.126 | 0.001 | 0.020 | 0.058 | 0.00 | 0.99 | 0.80 |
| ampc | 62 | 2964 | 0.462 | 0.113 | 0.000 | 0.002 | 0.025 | 0.00 | 0.32 | 0.48 |
| bace1 | 485 | 18706 | 0.620 | 0.185 | 0.001 | 0.012 | 0.068 | 0.00 | 0.70 | 1.20 |
| comt | 86 | 4012 | 0.393 | 0.081 | 0.000 | 0.000 | 0.001 | 0.00 | 0.00 | 0.00 |
| cp2c9 | 183 | 7757 | 0.576 | 0.223 | 0.238 | 0.153 | 0.176 | 7.16 | 3.18 | 2.08 |
| def | 161 | 5899 | 0.302 | 0.067 | 0.000 | 0.004 | 0.016 | 0.00 | 0.25 | 0.19 |
| fabp4 | 57 | 2912 | 0.625 | 0.232 | 0.159 | 0.106 | 0.164 | 5.28 | 2.47 | 2.81 |
| fgfr1 | 242 | 736 | 0.550 | 0.169 | 0.684 | 0.433 | 0.401 | 1.30 | 1.18 | 1.29 |
| fkb1a | 273 | 6105 | 0.565 | 0.174 | 0.013 | 0.047 | 0.111 | 0.37 | 1.76 | 1.58 |
| gria2 | 297 | 12358 | 0.594 | 0.188 | 0.008 | 0.023 | 0.102 | 0.34 | 1.89 | 1.95 |
| hs90a | 125 | 5067 | 0.527 | 0.135 | 0.000 | 0.006 | 0.035 | 0.00 | 0.48 | 0.72 |
| hxk4 | 127 | 4930 | 0.685 | 0.221 | 0.001 | 0.031 | 0.116 | 0.00 | 2.05 | 1.89 |
| jak2 | 153 | 6743 | 0.709 | 0.288 | 0.088 | 0.121 | 0.241 | 3.29 | 4.58 | 3.66 |
| mapk2 | 206 | 6450 | 0.647 | 0.212 | 0.056 | 0.051 | 0.124 | 0.98 | 1.94 | 1.99 |
| mk01 | 139 | 4767 | 0.741 | 0.286 | 0.119 | 0.105 | 0.213 | 2.92 | 3.31 | 3.39 |
| mk10 | 186 | 6900 | 0.726 | 0.302 | 0.185 | 0.153 | 0.265 | 4.84 | 4.73 | 4.30 |
| nos1 | 234 | 8307 | 0.603 | 0.214 | 0.010 | 0.060 | 0.157 | 0.86 | 2.74 | 2.69 |
| pa2ga | 127 | 5343 | 0.358 | 0.118 | 0.018 | 0.044 | 0.091 | 0.79 | 1.73 | 1.34 |
| pnph | 233 | 7249 | 0.571 | 0.166 | 0.051 | 0.054 | 0.082 | 1.73 | 1.20 | 1.20 |
| ppard | 288 | 13520 | 0.777 | 0.365 | 0.317 | 0.270 | 0.336 | 13.56 | 6.04 | 4.44 |
| ptn1 | 225 | 7658 | 0.682 | 0.321 | 0.274 | 0.291 | 0.339 | 10.75 | 6.77 | 3.96 |
| pur2 | 201 | 2926 | 0.623 | 0.163 | 0.000 | 0.007 | 0.054 | 0.00 | 0.20 | 0.85 |
| tysy | 311 | 7194 | 0.554 | 0.194 | 0.062 | 0.126 | 0.178 | 1.95 | 3.22 | 2.16 |

Supplementary Table 13. The virtual screening results by vinardo scorinig function of 23 targets based on the Apo protein structures.

| Target | Active | Total | AUC | logAUC | BEDROC(α=321.9) | BEDROC(α=80.5) | BEDROC(α=20.0) | EF1% | EF5% | EF10% |
| --- | --- | --- | --- | --- | --- | --- | --- | --- | --- | --- |
| ada | 262 | 5734 | 0.470 | 0.124 | 0.039 | 0.033 | 0.052 | 0.77 | 0.69 | 0.69 |
| ampc | 62 | 2964 | 0.530 | 0.150 | 0.000 | 0.010 | 0.067 | 0.00 | 1.94 | 1.13 |
| bace1 | 485 | 18706 | 0.581 | 0.198 | 0.042 | 0.078 | 0.139 | 2.27 | 2.56 | 1.96 |
| comt | 86 | 4012 | 0.642 | 0.200 | 0.000 | 0.022 | 0.105 | 0.00 | 1.87 | 1.75 |
| cp2c9 | 183 | 7757 | 0.628 | 0.213 | 0.064 | 0.081 | 0.133 | 2.75 | 2.30 | 1.86 |
| def | 161 | 5899 | 0.485 | 0.173 | 0.178 | 0.150 | 0.141 | 6.32 | 2.62 | 1.43 |
| fabp4 | 57 | 2912 | 0.887 | 0.432 | 0.114 | 0.205 | 0.442 | 7.05 | 9.51 | 7.20 |
| fgfr1 | 242 | 736 | 0.575 | 0.181 | 0.418 | 0.484 | 0.444 | 1.30 | 1.44 | 1.29 |
| fkb1a | 273 | 6105 | 0.592 | 0.187 | 0.011 | 0.055 | 0.115 | 0.37 | 1.61 | 1.50 |
| gria2 | 297 | 12358 | 0.652 | 0.215 | 0.040 | 0.061 | 0.126 | 2.03 | 2.09 | 1.82 |
| hs90a | 125 | 5067 | 0.398 | 0.095 | 0.000 | 0.001 | 0.018 | 0.00 | 0.32 | 0.32 |
| hxk4 | 127 | 4930 | 0.551 | 0.153 | 0.059 | 0.028 | 0.061 | 0.79 | 0.79 | 1.18 |
| jak2 | 153 | 6743 | 0.637 | 0.192 | 0.010 | 0.025 | 0.079 | 0.66 | 1.31 | 1.31 |
| mapk2 | 206 | 6450 | 0.721 | 0.290 | 0.114 | 0.175 | 0.259 | 6.36 | 4.67 | 3.35 |
| mk01 | 139 | 4767 | 0.759 | 0.350 | 0.234 | 0.248 | 0.364 | 8.76 | 7.49 | 4.83 |
| mk10 | 186 | 6900 | 0.706 | 0.230 | 0.009 | 0.045 | 0.124 | 1.08 | 2.04 | 1.94 |
| nos1 | 234 | 8307 | 0.600 | 0.174 | 0.001 | 0.016 | 0.065 | 0.00 | 1.11 | 1.07 |
| pa2ga | 127 | 5343 | 0.466 | 0.121 | 0.008 | 0.019 | 0.043 | 0.79 | 0.79 | 0.63 |
| pnph | 233 | 7249 | 0.579 | 0.169 | 0.016 | 0.030 | 0.074 | 0.86 | 0.95 | 1.07 |
| ppard | 288 | 13520 | 0.578 | 0.215 | 0.065 | 0.084 | 0.161 | 3.48 | 2.99 | 2.50 |
| ptn1 | 225 | 7658 | 0.711 | 0.256 | 0.072 | 0.084 | 0.173 | 2.24 | 2.58 | 2.58 |
| pur2 | 201 | 2926 | 0.721 | 0.252 | 0.104 | 0.125 | 0.201 | 2.01 | 1.89 | 2.54 |
| tysy | 311 | 7194 | 0.730 | 0.268 | 0.105 | 0.129 | 0.214 | 3.26 | 3.16 | 2.86 |

Supplementary Table 14. The average values for virtual screening results by different functions of 51 targets based on the protein structures in DUD-E databases. The function with prefix “irva_” means the consensun scoring method using idock, rf_score, vinardo and autodock4. The function with prefix “irv_” means the consensun scoring method using idock, rf_score and vinardo. The function with prefix “ir_” means the consensun scoring method using idock and rf_score.

| Function | AUC | logAUC | BEDROC(α=321.9) | BEDROC(α=80.5) | BEDROC(α=20.0) | EF1% | EF5% | EF10% |
| --- | --- | --- | --- | --- | --- | --- | --- | --- |
| autodock4 | 0.645 | 0.270 | 0.229 | 0.194 | 0.235 | 7.00 | 3.92 | 2.90 |
| idock | 0.647 | 0.252 | 0.204 | 0.156 | 0.199 | 6.34 | 3.23 | 2.57 |
| vinardo | 0.639 | 0.245 | 0.172 | 0.152 | 0.198 | 5.44 | 3.28 | 2.51 |
| rf_score | 0.625 | 0.228 | 0.118 | 0.116 | 0.172 | 4.04 | 2.87 | 2.34 |
| irva_exp_z_score | 0.686 | 0.290 | 0.248 | 0.204 | 0.253 | 7.74 | 4.23 | 3.20 |
| irva_best_z_score | 0.687 | 0.285 | 0.222 | 0.193 | 0.246 | 7.04 | 4.16 | 3.10 |
| irva_exp_rank | 0.689 | 0.284 | 0.247 | 0.193 | 0.244 | 7.42 | 4.05 | 3.10 |
| irva_sum_z_score | 0.676 | 0.282 | 0.250 | 0.197 | 0.243 | 7.70 | 4.06 | 3.07 |
| irva_best_rank | 0.688 | 0.280 | 0.187 | 0.180 | 0.243 | 6.53 | 4.14 | 3.12 |
| irva_exp_aass | 0.671 | 0.279 | 0.248 | 0.195 | 0.240 | 7.62 | 3.98 | 3.02 |
| irva_sum_aass | 0.669 | 0.278 | 0.248 | 0.194 | 0.238 | 7.51 | 3.95 | 2.97 |
| irva_best_aass | 0.678 | 0.272 | 0.192 | 0.173 | 0.230 | 6.36 | 3.89 | 3.00 |
| irva_sum_rank | 0.673 | 0.272 | 0.234 | 0.179 | 0.227 | 6.78 | 3.72 | 2.90 |
| irva_rbv | 0.659 | 0.260 | 0.211 | 0.171 | 0.223 | 6.74 | 3.64 | 3.00 |
| irva_worst_aass | 0.633 | 0.250 | 0.204 | 0.164 | 0.206 | 6.19 | 3.43 | 2.61 |
| irva_worst_z_score | 0.634 | 0.249 | 0.223 | 0.166 | 0.204 | 6.37 | 3.36 | 2.53 |
| irva_worst_rank | 0.635 | 0.249 | 0.223 | 0.164 | 0.202 | 6.26 | 3.30 | 2.51 |
| irv_exp_z_score | 0.673 | 0.268 | 0.202 | 0.168 | 0.218 | 6.47 | 3.65 | 2.81 |
| irv_best_z_score | 0.674 | 0.266 | 0.188 | 0.164 | 0.217 | 6.25 | 3.59 | 2.78 |
| irv_best_rank | 0.676 | 0.266 | 0.168 | 0.160 | 0.218 | 6.06 | 3.68 | 2.82 |
| irv_exp_aass | 0.662 | 0.262 | 0.200 | 0.164 | 0.211 | 6.42 | 3.55 | 2.71 |
| irv_sum_z_score | 0.665 | 0.261 | 0.198 | 0.162 | 0.211 | 6.13 | 3.50 | 2.72 |
| irv_exp_rank | 0.676 | 0.260 | 0.192 | 0.159 | 0.214 | 5.83 | 3.56 | 2.84 |
| irv_best_aass | 0.666 | 0.260 | 0.174 | 0.156 | 0.211 | 6.00 | 3.60 | 2.77 |
| irv_sum_aass | 0.660 | 0.260 | 0.199 | 0.162 | 0.209 | 6.25 | 3.48 | 2.69 |
| irv_sum_rank | 0.663 | 0.254 | 0.187 | 0.149 | 0.199 | 5.58 | 3.23 | 2.56 |
| irv_worst_z_score | 0.633 | 0.238 | 0.177 | 0.141 | 0.183 | 5.18 | 3.02 | 2.34 |
| irv_worst_rank | 0.634 | 0.238 | 0.178 | 0.141 | 0.183 | 5.26 | 3.01 | 2.32 |
| irv_rbv | 0.633 | 0.237 | 0.179 | 0.146 | 0.203 | 5.23 | 3.28 | 2.68 |
| irv_worst_aass | 0.633 | 0.236 | 0.164 | 0.134 | 0.181 | 5.09 | 2.97 | 2.35 |
| ir_exp_z_score | 0.656 | 0.256 | 0.197 | 0.154 | 0.203 | 6.07 | 3.36 | 2.67 |
| ir_best_rank | 0.658 | 0.255 | 0.170 | 0.151 | 0.203 | 5.96 | 3.38 | 2.65 |
| ir_best_z_score | 0.656 | 0.255 | 0.187 | 0.149 | 0.200 | 5.89 | 3.33 | 2.63 |
| ir_exp_aass | 0.651 | 0.253 | 0.198 | 0.156 | 0.202 | 5.99 | 3.35 | 2.59 |
| ir_sum_aass | 0.650 | 0.252 | 0.198 | 0.155 | 0.201 | 5.99 | 3.30 | 2.58 |
| ir_sum_z_score | 0.652 | 0.252 | 0.198 | 0.155 | 0.200 | 5.87 | 3.31 | 2.59 |
| ir_best_aass | 0.652 | 0.251 | 0.166 | 0.145 | 0.199 | 5.59 | 3.34 | 2.65 |
| ir_exp_rank | 0.658 | 0.250 | 0.187 | 0.148 | 0.201 | 5.60 | 3.38 | 2.67 |
| ir_sum_rank | 0.651 | 0.248 | 0.186 | 0.146 | 0.192 | 5.53 | 3.19 | 2.50 |
| ir_worst_z_score | 0.636 | 0.239 | 0.170 | 0.139 | 0.184 | 5.44 | 3.10 | 2.39 |
| ir_worst_rank | 0.635 | 0.239 | 0.180 | 0.142 | 0.184 | 5.40 | 3.08 | 2.37 |
| ir_worst_aass | 0.633 | 0.237 | 0.189 | 0.140 | 0.180 | 5.43 | 2.90 | 2.33 |
| ir_rbv | 0.610 | 0.220 | 0.193 | 0.146 | 0.199 | 5.59 | 3.33 | 2.57 |

Supplementary Table 15. The average values for virtual screening results by different functions of 51 targets based on the protein structures of AlphaFold. The function with prefix “irva_” means the consensun scoring method using idock, rf_score, vinardo and autodock4. The function with prefix “irv_” means the consensun scoring method using idock, rf_score and vinardo. The function with prefix “ir_” means the consensun scoring method using idock and rf_score.

| Function | AUC | logAUC | BEDROC(α=321.9) | BEDROC(α=80.5) | BEDROC(α=20.0) | EF1% | EF5% | EF10% |
| --- | --- | --- | --- | --- | --- | --- | --- | --- |
| autodock4 | 0.622 | 0.240 | 0.183 | 0.150 | 0.196 | 5.28 | 3.24 | 2.52 |
| idock | 0.635 | 0.228 | 0.126 | 0.114 | 0.166 | 3.98 | 2.62 | 2.23 |
| vinardo | 0.614 | 0.217 | 0.128 | 0.109 | 0.157 | 3.89 | 2.53 | 2.11 |
| rf_score | 0.609 | 0.208 | 0.087 | 0.086 | 0.140 | 2.96 | 2.25 | 1.98 |
| irva_exp_z_score | 0.665 | 0.256 | 0.164 | 0.151 | 0.208 | 5.52 | 3.55 | 2.74 |
| irva_exp_rank | 0.667 | 0.252 | 0.152 | 0.140 | 0.202 | 4.80 | 3.42 | 2.72 |
| irva_best_z_score | 0.664 | 0.251 | 0.164 | 0.141 | 0.197 | 4.85 | 3.32 | 2.59 |
| irva_best_rank | 0.665 | 0.251 | 0.147 | 0.136 | 0.197 | 4.83 | 3.27 | 2.65 |
| irva_sum_z_score | 0.660 | 0.249 | 0.144 | 0.139 | 0.198 | 4.94 | 3.33 | 2.64 |
| irva_exp_aass | 0.653 | 0.248 | 0.147 | 0.141 | 0.199 | 5.08 | 3.38 | 2.64 |
| irva_sum_aass | 0.653 | 0.247 | 0.145 | 0.139 | 0.197 | 4.96 | 3.30 | 2.60 |
| irva_best_aass | 0.659 | 0.243 | 0.136 | 0.121 | 0.183 | 4.41 | 3.01 | 2.52 |
| irva_sum_rank | 0.655 | 0.243 | 0.145 | 0.130 | 0.186 | 4.44 | 3.07 | 2.48 |
| irva_rbv | 0.631 | 0.231 | 0.135 | 0.127 | 0.185 | 4.28 | 3.00 | 2.59 |
| irva_worst_rank | 0.615 | 0.222 | 0.141 | 0.119 | 0.165 | 4.20 | 2.69 | 2.14 |
| irva_worst_aass | 0.616 | 0.219 | 0.125 | 0.112 | 0.159 | 3.88 | 2.65 | 2.14 |
| irva_worst_z_score | 0.614 | 0.219 | 0.127 | 0.112 | 0.160 | 3.77 | 2.60 | 2.11 |
| irv_exp_z_score | 0.657 | 0.239 | 0.124 | 0.118 | 0.177 | 3.94 | 2.92 | 2.40 |
| irv_best_rank | 0.657 | 0.238 | 0.121 | 0.115 | 0.174 | 4.11 | 2.88 | 2.37 |
| irv_best_z_score | 0.657 | 0.236 | 0.125 | 0.114 | 0.171 | 3.80 | 2.82 | 2.29 |
| irv_best_aass | 0.652 | 0.235 | 0.119 | 0.111 | 0.171 | 4.02 | 2.79 | 2.37 |
| irv_exp_aass | 0.647 | 0.234 | 0.116 | 0.113 | 0.173 | 3.89 | 2.90 | 2.35 |
| irv_sum_z_score | 0.653 | 0.234 | 0.112 | 0.110 | 0.171 | 3.70 | 2.82 | 2.31 |
| irv_exp_rank | 0.658 | 0.234 | 0.119 | 0.112 | 0.176 | 3.69 | 2.94 | 2.40 |
| irv_sum_aass | 0.647 | 0.233 | 0.114 | 0.111 | 0.171 | 3.88 | 2.85 | 2.33 |
| irv_sum_rank | 0.648 | 0.230 | 0.116 | 0.105 | 0.163 | 3.50 | 2.63 | 2.24 |
| irv_worst_rank | 0.617 | 0.214 | 0.109 | 0.099 | 0.149 | 3.56 | 2.38 | 1.99 |
| irv_rbv | 0.607 | 0.212 | 0.107 | 0.108 | 0.169 | 3.76 | 2.74 | 2.29 |
| irv_worst_z_score | 0.616 | 0.212 | 0.097 | 0.093 | 0.145 | 3.18 | 2.28 | 1.96 |
| irv_worst_aass | 0.612 | 0.208 | 0.098 | 0.088 | 0.138 | 2.95 | 2.17 | 1.91 |
| ir_exp_z_score | 0.645 | 0.232 | 0.118 | 0.116 | 0.171 | 3.85 | 2.82 | 2.31 |
| ir_best_rank | 0.645 | 0.230 | 0.110 | 0.108 | 0.166 | 3.80 | 2.75 | 2.25 |
| ir_exp_aass | 0.637 | 0.230 | 0.128 | 0.117 | 0.170 | 4.04 | 2.84 | 2.27 |
| ir_best_z_score | 0.645 | 0.230 | 0.113 | 0.110 | 0.166 | 3.49 | 2.68 | 2.22 |
| ir_sum_z_score | 0.642 | 0.230 | 0.122 | 0.114 | 0.169 | 3.81 | 2.80 | 2.27 |
| ir_sum_aass | 0.637 | 0.230 | 0.128 | 0.116 | 0.169 | 3.98 | 2.85 | 2.27 |
| ir_exp_rank | 0.646 | 0.228 | 0.122 | 0.111 | 0.169 | 4.07 | 2.82 | 2.32 |
| ir_sum_rank | 0.641 | 0.227 | 0.122 | 0.108 | 0.161 | 3.98 | 2.67 | 2.19 |
| ir_best_aass | 0.641 | 0.227 | 0.098 | 0.099 | 0.160 | 3.56 | 2.60 | 2.25 |
| ir_worst_rank | 0.623 | 0.218 | 0.119 | 0.105 | 0.155 | 3.79 | 2.55 | 2.09 |
| ir_worst_z_score | 0.622 | 0.217 | 0.107 | 0.099 | 0.153 | 3.45 | 2.52 | 2.06 |
| ir_worst_aass | 0.613 | 0.212 | 0.107 | 0.096 | 0.146 | 3.34 | 2.32 | 1.98 |
| ir_rbv | 0.587 | 0.202 | 0.117 | 0.110 | 0.169 | 3.68 | 2.86 | 2.23 |

Supplementary Table 16. The average values for virtual screening results by different functions of 23 targets based on the Apo protein structures. The function with prefix “irva_” means the consensun scoring method using idock, rf_score, vinardo and autodock4. The function with prefix “irv_” means the consensun scoring method using idock, rf_score and vinardo. The function with prefix “ir_” means the consensun scoring method using idock and rf_score.

| Function | AUC | logAUC | BEDROC(α=321.9) | BEDROC(α=80.5) | BEDROC(α=20.0) | EF1% | EF5% | EF10% |
| --- | --- | --- | --- | --- | --- | --- | --- | --- |
| vinardo | 0.617 | 0.210 | 0.074 | 0.095 | 0.157 | 2.31 | 2.47 | 2.04 |
| autodock4 | 0.580 | 0.206 | 0.124 | 0.122 | 0.164 | 3.59 | 2.40 | 2.03 |
| idock | 0.603 | 0.202 | 0.093 | 0.089 | 0.139 | 2.31 | 2.10 | 1.80 |
| rf_score | 0.581 | 0.198 | 0.099 | 0.092 | 0.146 | 2.44 | 2.25 | 1.96 |
| irva_best_aass | 0.632 | 0.226 | 0.102 | 0.112 | 0.178 | 2.91 | 2.69 | 2.36 |
| irva_exp_rank | 0.635 | 0.223 | 0.107 | 0.115 | 0.173 | 3.62 | 2.70 | 2.24 |
| irva_exp_z_score | 0.627 | 0.223 | 0.103 | 0.110 | 0.170 | 3.22 | 2.62 | 2.22 |
| irva_best_rank | 0.636 | 0.221 | 0.100 | 0.104 | 0.164 | 2.60 | 2.39 | 2.17 |
| irva_best_z_score | 0.634 | 0.221 | 0.096 | 0.104 | 0.162 | 3.43 | 2.42 | 2.11 |
| irva_exp_aass | 0.617 | 0.219 | 0.107 | 0.113 | 0.169 | 3.22 | 2.56 | 2.15 |
| irva_sum_z_score | 0.617 | 0.219 | 0.111 | 0.114 | 0.169 | 3.50 | 2.59 | 2.17 |
| irva_sum_rank | 0.617 | 0.218 | 0.108 | 0.115 | 0.166 | 3.88 | 2.57 | 2.06 |
| irva_sum_aass | 0.615 | 0.218 | 0.108 | 0.113 | 0.168 | 3.28 | 2.56 | 2.15 |
| irva_rbv | 0.603 | 0.212 | 0.114 | 0.109 | 0.161 | 3.19 | 2.39 | 2.16 |
| irva_worst_z_score | 0.587 | 0.205 | 0.130 | 0.114 | 0.156 | 3.67 | 2.42 | 1.94 |
| irva_worst_rank | 0.588 | 0.205 | 0.118 | 0.113 | 0.156 | 3.65 | 2.43 | 1.95 |
| irva_worst_aass | 0.581 | 0.200 | 0.109 | 0.105 | 0.148 | 3.41 | 2.19 | 1.86 |
| irv_best_aass | 0.626 | 0.220 | 0.094 | 0.102 | 0.167 | 2.95 | 2.59 | 2.24 |
| irv_exp_z_score | 0.625 | 0.216 | 0.086 | 0.098 | 0.159 | 2.78 | 2.45 | 2.09 |
| irv_exp_rank | 0.629 | 0.215 | 0.106 | 0.105 | 0.164 | 2.83 | 2.56 | 2.13 |
| irv_best_rank | 0.629 | 0.215 | 0.091 | 0.094 | 0.155 | 2.52 | 2.27 | 2.14 |
| irv_exp_aass | 0.619 | 0.215 | 0.097 | 0.101 | 0.160 | 2.81 | 2.54 | 2.06 |
| irv_sum_rank | 0.620 | 0.215 | 0.104 | 0.103 | 0.157 | 2.96 | 2.38 | 2.04 |
| irv_sum_z_score | 0.620 | 0.214 | 0.092 | 0.099 | 0.159 | 2.68 | 2.48 | 2.03 |
| irv_sum_aass | 0.618 | 0.214 | 0.098 | 0.100 | 0.158 | 2.68 | 2.54 | 2.04 |
| irv_best_z_score | 0.626 | 0.214 | 0.078 | 0.092 | 0.153 | 2.19 | 2.29 | 2.09 |
| irv_worst_rank | 0.597 | 0.204 | 0.112 | 0.103 | 0.149 | 3.17 | 2.17 | 1.91 |
| irv_worst_z_score | 0.597 | 0.204 | 0.122 | 0.105 | 0.149 | 3.00 | 2.22 | 1.86 |
| irv_rbv | 0.588 | 0.201 | 0.090 | 0.103 | 0.154 | 3.08 | 2.36 | 1.90 |
| irv_worst_aass | 0.591 | 0.198 | 0.083 | 0.090 | 0.139 | 2.53 | 2.02 | 1.78 |
| ir_best_aass | 0.609 | 0.213 | 0.104 | 0.102 | 0.160 | 2.90 | 2.48 | 2.16 |
| ir_best_rank | 0.610 | 0.208 | 0.102 | 0.093 | 0.149 | 2.68 | 2.28 | 2.05 |
| ir_exp_z_score | 0.606 | 0.206 | 0.102 | 0.092 | 0.146 | 2.40 | 2.22 | 1.97 |
| ir_best_z_score | 0.608 | 0.206 | 0.089 | 0.086 | 0.145 | 2.23 | 2.09 | 2.01 |
| ir_exp_aass | 0.601 | 0.205 | 0.096 | 0.097 | 0.147 | 2.93 | 2.28 | 1.89 |
| ir_sum_z_score | 0.603 | 0.204 | 0.095 | 0.095 | 0.146 | 2.86 | 2.28 | 1.90 |
| ir_sum_rank | 0.602 | 0.204 | 0.099 | 0.093 | 0.145 | 2.90 | 2.27 | 1.88 |
| ir_sum_aass | 0.600 | 0.204 | 0.097 | 0.096 | 0.146 | 2.92 | 2.26 | 1.88 |
| ir_exp_rank | 0.609 | 0.203 | 0.097 | 0.093 | 0.151 | 2.78 | 2.28 | 1.99 |
| ir_worst_z_score | 0.588 | 0.199 | 0.093 | 0.098 | 0.143 | 3.06 | 2.24 | 1.80 |
| ir_worst_rank | 0.588 | 0.198 | 0.101 | 0.092 | 0.141 | 2.86 | 2.19 | 1.74 |
| ir_worst_aass | 0.577 | 0.189 | 0.095 | 0.082 | 0.129 | 2.20 | 1.89 | 1.66 |
| ir_rbv | 0.569 | 0.187 | 0.089 | 0.090 | 0.145 | 2.68 | 2.26 | 1.80 |

Supplementary Table 17. The virtual screening results by exp_z_score consensus scoring method (using idock, rf_score, vinardo and autodock4) of 51 targets based on the protein structures in DUD-E databases.

| Target | Active | Total | AUC | logAUC | BEDROC(α=321.9) | BEDROC(α=80.5) | BEDROC(α=20.0) | EF1% | EF5% | EF10% |
| --- | --- | --- | --- | --- | --- | --- | --- | --- | --- | --- |
| ada | 262 | 5734 | 0.329 | 0.086 | 0.013 | 0.015 | 0.046 | 0.38 | 0.69 | 0.80 |
| adrb1 | 458 | 16416 | 0.678 | 0.270 | 0.187 | 0.162 | 0.232 | 5.90 | 4.33 | 3.08 |
| adrb2 | 447 | 15702 | 0.697 | 0.298 | 0.248 | 0.222 | 0.281 | 8.28 | 5.19 | 3.51 |
| akt1 | 423 | 16999 | 0.681 | 0.224 | 0.124 | 0.084 | 0.116 | 3.80 | 1.80 | 1.47 |
| akt2 | 190 | 7142 | 0.786 | 0.315 | 0.130 | 0.147 | 0.250 | 4.24 | 4.42 | 3.63 |
| ampc | 62 | 2964 | 0.511 | 0.143 | 0.048 | 0.036 | 0.062 | 1.65 | 0.97 | 0.97 |
| aofb | 168 | 7099 | 0.640 | 0.223 | 0.029 | 0.051 | 0.144 | 1.81 | 2.27 | 2.68 |
| bace1 | 485 | 18705 | 0.786 | 0.320 | 0.156 | 0.158 | 0.259 | 6.81 | 4.29 | 3.88 |
| casp3 | 350 | 11172 | 0.615 | 0.198 | 0.040 | 0.062 | 0.117 | 2.01 | 1.83 | 1.71 |
| comt | 86 | 4012 | 0.463 | 0.111 | 0.000 | 0.004 | 0.022 | 0.00 | 0.23 | 0.47 |
| cp2c9 | 183 | 7757 | 0.625 | 0.248 | 0.220 | 0.152 | 0.206 | 6.06 | 3.72 | 2.84 |
| cp3a4 | 363 | 12303 | 0.558 | 0.182 | 0.029 | 0.050 | 0.112 | 1.65 | 1.65 | 1.71 |
| cxcr4 | 122 | 3522 | 0.833 | 0.369 | 0.497 | 0.276 | 0.327 | 8.25 | 4.92 | 3.85 |
| def | 161 | 5899 | 0.489 | 0.132 | 0.025 | 0.029 | 0.049 | 1.26 | 0.50 | 0.75 |
| fa7 | 185 | 6487 | 0.883 | 0.435 | 0.330 | 0.332 | 0.441 | 11.51 | 8.23 | 5.74 |
| fabp4 | 57 | 2912 | 0.813 | 0.502 | 0.580 | 0.524 | 0.497 | 35.23 | 9.16 | 5.09 |
| fak1 | 114 | 5516 | 0.703 | 0.259 | 0.080 | 0.077 | 0.177 | 2.64 | 2.64 | 3.07 |
| fgfr1 | 242 | 736 | 0.551 | 0.171 | 0.377 | 0.432 | 0.424 | 1.30 | 1.35 | 1.29 |
| fkb1a | 273 | 6105 | 0.733 | 0.261 | 0.139 | 0.124 | 0.190 | 2.20 | 2.20 | 2.57 |
| gcr | 563 | 15748 | 0.631 | 0.261 | 0.297 | 0.223 | 0.246 | 6.77 | 4.16 | 2.95 |
| glcm | 313 | 4150 | 0.620 | 0.191 | 0.006 | 0.035 | 0.121 | 0.32 | 0.90 | 1.53 |
| gria2 | 297 | 12356 | 0.737 | 0.297 | 0.188 | 0.135 | 0.235 | 4.74 | 4.18 | 3.71 |
| grik1 | 152 | 6769 | 0.639 | 0.260 | 0.159 | 0.161 | 0.224 | 6.65 | 4.48 | 2.96 |
| hivpr | 1395 | 37673 | 0.741 | 0.287 | 0.157 | 0.158 | 0.236 | 4.17 | 3.60 | 3.28 |
| hs90a | 125 | 5067 | 0.363 | 0.092 | 0.001 | 0.009 | 0.030 | 0.00 | 0.48 | 0.48 |
| hxk4 | 127 | 4930 | 0.566 | 0.147 | 0.009 | 0.018 | 0.033 | 0.79 | 0.32 | 0.47 |
| inha | 71 | 2389 | 0.710 | 0.318 | 0.188 | 0.263 | 0.332 | 8.78 | 6.22 | 4.38 |
| jak2 | 153 | 6743 | 0.762 | 0.335 | 0.410 | 0.232 | 0.276 | 9.87 | 4.58 | 3.79 |
| kif11 | 197 | 7109 | 0.843 | 0.469 | 0.598 | 0.455 | 0.494 | 19.31 | 8.84 | 6.25 |
| kith | 132 | 2998 | 0.430 | 0.191 | 0.392 | 0.257 | 0.208 | 7.05 | 3.20 | 2.20 |
| mapk2 | 206 | 6450 | 0.759 | 0.345 | 0.358 | 0.302 | 0.344 | 9.30 | 6.13 | 4.13 |
| mcr | 193 | 5432 | 0.638 | 0.262 | 0.558 | 0.257 | 0.216 | 8.86 | 2.80 | 2.54 |
| mk01 | 139 | 4767 | 0.789 | 0.301 | 0.081 | 0.072 | 0.205 | 1.46 | 2.88 | 3.46 |
| mk10 | 186 | 6900 | 0.728 | 0.263 | 0.066 | 0.098 | 0.184 | 3.23 | 3.55 | 2.47 |
| nos1 | 234 | 8307 | 0.587 | 0.187 | 0.030 | 0.051 | 0.098 | 1.28 | 1.54 | 1.41 |
| pa2ga | 127 | 5343 | 0.687 | 0.240 | 0.013 | 0.051 | 0.154 | 1.59 | 2.84 | 2.84 |
| plk1 | 155 | 7034 | 0.636 | 0.230 | 0.060 | 0.098 | 0.173 | 3.89 | 3.36 | 2.58 |
| pnph | 233 | 7249 | 0.808 | 0.374 | 0.464 | 0.343 | 0.358 | 12.10 | 6.19 | 4.13 |
| ppard | 288 | 13520 | 0.735 | 0.267 | 0.039 | 0.067 | 0.174 | 2.43 | 2.99 | 2.81 |
| ptn1 | 225 | 7658 | 0.881 | 0.530 | 0.687 | 0.545 | 0.577 | 22.84 | 10.78 | 6.85 |
| pur2 | 201 | 2926 | 0.917 | 0.471 | 0.515 | 0.507 | 0.576 | 8.53 | 7.28 | 6.43 |
| pygm | 114 | 4154 | 0.413 | 0.101 | 0.001 | 0.011 | 0.026 | 0.00 | 0.53 | 0.35 |
| pyrd | 134 | 6782 | 0.734 | 0.261 | 0.072 | 0.084 | 0.165 | 4.53 | 2.84 | 2.69 |
| rock1 | 203 | 6580 | 0.722 | 0.278 | 0.403 | 0.216 | 0.209 | 7.98 | 3.25 | 2.32 |
| rxra | 162 | 7869 | 0.889 | 0.549 | 0.695 | 0.507 | 0.574 | 26.16 | 11.25 | 7.29 |
| sahh | 190 | 3673 | 0.439 | 0.107 | 0.000 | 0.009 | 0.032 | 0.00 | 0.32 | 0.42 |
| thb | 168 | 7821 | 0.849 | 0.510 | 0.674 | 0.498 | 0.530 | 26.86 | 10.24 | 6.31 |
| tryb1 | 171 | 7884 | 0.771 | 0.306 | 0.142 | 0.135 | 0.233 | 5.91 | 4.10 | 3.69 |
| tysy | 311 | 7194 | 0.820 | 0.416 | 0.590 | 0.437 | 0.455 | 11.73 | 6.96 | 5.47 |
| wee1 | 137 | 6371 | 0.945 | 0.725 | 0.954 | 0.771 | 0.766 | 41.34 | 15.21 | 8.32 |
| xiap | 129 | 5342 | 0.823 | 0.465 | 0.578 | 0.465 | 0.489 | 21.10 | 9.15 | 5.66 |

Supplementary Table 18. The virtual screening results by exp_z_score consensus scoring method (using idock, rf_score, vinardo and autodock4) of 51 targets based on the protein structures of AlphaFold.

| Target | Active | Total | AUC | logAUC | BEDROC(α=321.9) | BEDROC(α=80.5) | BEDROC(α=20.0) | EF1% | EF5% | EF10% |
| --- | --- | --- | --- | --- | --- | --- | --- | --- | --- | --- |
| ada | 262 | 5734 | 0.394 | 0.115 | 0.177 | 0.079 | 0.068 | 2.30 | 0.99 | 0.69 |
| adrb1 | 458 | 16416 | 0.649 | 0.242 | 0.111 | 0.132 | 0.190 | 5.03 | 3.19 | 2.51 |
| adrb2 | 447 | 15702 | 0.687 | 0.264 | 0.149 | 0.145 | 0.213 | 4.70 | 3.71 | 2.89 |
| akt1 | 423 | 16999 | 0.758 | 0.319 | 0.157 | 0.216 | 0.284 | 8.08 | 5.44 | 3.64 |
| akt2 | 190 | 7142 | 0.792 | 0.360 | 0.164 | 0.253 | 0.358 | 8.47 | 7.05 | 4.63 |
| ampc | 62 | 2964 | 0.527 | 0.136 | 0.000 | 0.006 | 0.033 | 0.00 | 0.32 | 0.65 |
| aofb | 168 | 7099 | 0.700 | 0.278 | 0.146 | 0.155 | 0.233 | 6.64 | 4.30 | 3.34 |
| bace1 | 485 | 18706 | 0.790 | 0.351 | 0.217 | 0.208 | 0.326 | 7.22 | 6.11 | 4.68 |
| casp3 | 350 | 11172 | 0.718 | 0.309 | 0.205 | 0.239 | 0.304 | 8.63 | 5.38 | 3.91 |
| comt | 86 | 4012 | 0.432 | 0.096 | 0.000 | 0.000 | 0.002 | 0.00 | 0.00 | 0.00 |
| cp2c9 | 183 | 7757 | 0.618 | 0.239 | 0.158 | 0.160 | 0.195 | 8.26 | 3.72 | 2.35 |
| cp3a4 | 363 | 12303 | 0.562 | 0.182 | 0.048 | 0.058 | 0.106 | 3.03 | 1.60 | 1.71 |
| cxcr4 | 122 | 3536 | 0.722 | 0.231 | 0.113 | 0.091 | 0.119 | 2.48 | 1.81 | 1.31 |
| def | 161 | 5899 | 0.615 | 0.231 | 0.159 | 0.129 | 0.186 | 5.05 | 3.24 | 2.61 |
| fa7 | 185 | 6487 | 0.804 | 0.352 | 0.254 | 0.233 | 0.324 | 7.12 | 5.95 | 4.27 |
| fabp4 | 57 | 2912 | 0.734 | 0.358 | 0.288 | 0.294 | 0.359 | 17.62 | 7.75 | 4.39 |
| fak1 | 114 | 5515 | 0.695 | 0.231 | 0.087 | 0.055 | 0.121 | 2.64 | 2.29 | 1.84 |
| fgfr1 | 242 | 736 | 0.555 | 0.169 | 0.421 | 0.368 | 0.397 | 1.30 | 1.10 | 1.29 |
| fkb1a | 273 | 6105 | 0.747 | 0.292 | 0.258 | 0.234 | 0.252 | 7.33 | 3.67 | 3.01 |
| gcr | 563 | 15674 | 0.504 | 0.154 | 0.104 | 0.082 | 0.098 | 2.50 | 1.56 | 1.23 |
| glcm | 313 | 4150 | 0.434 | 0.104 | 0.000 | 0.002 | 0.027 | 0.00 | 0.06 | 0.32 |
| gria2 | 296 | 12327 | 0.558 | 0.227 | 0.185 | 0.161 | 0.203 | 6.43 | 3.79 | 2.50 |
| grik1 | 152 | 6767 | 0.765 | 0.274 | 0.033 | 0.077 | 0.179 | 2.66 | 3.16 | 2.96 |
| hivpr | 1395 | 37673 | 0.755 | 0.334 | 0.351 | 0.283 | 0.329 | 8.76 | 5.34 | 4.20 |
| hs90a | 125 | 5067 | 0.323 | 0.070 | 0.000 | 0.000 | 0.008 | 0.00 | 0.00 | 0.08 |
| hxk4 | 127 | 4930 | 0.577 | 0.154 | 0.002 | 0.012 | 0.041 | 0.00 | 0.47 | 0.71 |
| inha | 71 | 2389 | 0.671 | 0.310 | 0.188 | 0.290 | 0.340 | 10.24 | 6.50 | 3.96 |
| jak2 | 153 | 6743 | 0.681 | 0.245 | 0.088 | 0.094 | 0.167 | 4.60 | 3.01 | 2.48 |
| kif11 | 197 | 7109 | 0.712 | 0.250 | 0.011 | 0.054 | 0.161 | 1.52 | 2.24 | 2.80 |
| kith | 130 | 2824 | 0.737 | 0.338 | 0.333 | 0.314 | 0.374 | 6.98 | 5.70 | 4.85 |
| mapk2 | 206 | 6450 | 0.780 | 0.382 | 0.474 | 0.374 | 0.397 | 14.19 | 7.29 | 4.76 |
| mcr | 193 | 5377 | 0.595 | 0.202 | 0.273 | 0.139 | 0.139 | 4.73 | 2.18 | 1.71 |
| mk01 | 139 | 4767 | 0.769 | 0.280 | 0.014 | 0.054 | 0.183 | 2.19 | 2.59 | 3.10 |
| mk10 | 186 | 6900 | 0.767 | 0.285 | 0.097 | 0.106 | 0.201 | 4.30 | 3.55 | 3.28 |
| nos1 | 234 | 8307 | 0.566 | 0.176 | 0.005 | 0.024 | 0.082 | 0.86 | 1.03 | 1.54 |
| pa2ga | 127 | 5343 | 0.469 | 0.128 | 0.006 | 0.017 | 0.054 | 0.79 | 1.26 | 0.95 |
| plk1 | 155 | 7034 | 0.652 | 0.211 | 0.001 | 0.024 | 0.106 | 0.00 | 1.68 | 2.07 |
| pnph | 233 | 7249 | 0.714 | 0.301 | 0.268 | 0.231 | 0.283 | 7.78 | 4.98 | 3.70 |
| ppard | 288 | 13520 | 0.781 | 0.313 | 0.087 | 0.130 | 0.250 | 5.56 | 4.72 | 3.92 |
| ptn1 | 225 | 7658 | 0.828 | 0.440 | 0.756 | 0.466 | 0.437 | 18.36 | 7.31 | 4.94 |
| pur2 | 201 | 2926 | 0.775 | 0.246 | 0.000 | 0.008 | 0.112 | 0.00 | 0.40 | 1.70 |
| pygm | 114 | 4159 | 0.606 | 0.200 | 0.056 | 0.075 | 0.122 | 1.78 | 2.11 | 1.67 |
| pyrd | 134 | 6782 | 0.688 | 0.238 | 0.028 | 0.063 | 0.151 | 2.27 | 2.99 | 2.39 |
| rock1 | 203 | 6580 | 0.714 | 0.262 | 0.327 | 0.169 | 0.189 | 6.48 | 3.15 | 2.51 |
| rxra | 162 | 7869 | 0.873 | 0.485 | 0.564 | 0.404 | 0.495 | 20.55 | 10.14 | 6.43 |
| sahh | 190 | 3673 | 0.349 | 0.076 | 0.000 | 0.000 | 0.007 | 0.00 | 0.00 | 0.11 |
| thb | 168 | 7805 | 0.780 | 0.368 | 0.268 | 0.244 | 0.360 | 11.32 | 7.39 | 5.24 |
| tryb1 | 171 | 7883 | 0.805 | 0.352 | 0.168 | 0.178 | 0.312 | 7.68 | 5.85 | 4.86 |
| tysy | 311 | 7194 | 0.751 | 0.307 | 0.222 | 0.218 | 0.288 | 4.56 | 4.64 | 3.54 |
| wee1 | 137 | 6371 | 0.871 | 0.442 | 0.326 | 0.339 | 0.430 | 18.45 | 8.19 | 5.55 |
| xiap | 129 | 5336 | 0.542 | 0.132 | 0.000 | 0.000 | 0.011 | 0.00 | 0.00 | 0.16 |

Supplementary Table 19. The virtual screening results by exp_z_score consensus scoring method (using idock, rf_score, vinardo and autodock4) of 23 targets based on the Apo protein structures.

| Target | Active | Total | AUC | logAUC | BEDROC(α=321.9) | BEDROC(α=80.5) | BEDROC(α=20.0) | EF1% | EF5% | EF10% |
| --- | --- | --- | --- | --- | --- | --- | --- | --- | --- | --- |
| ada | 262 | 5734 | 0.346 | 0.087 | 0.000 | 0.009 | 0.038 | 0.00 | 0.54 | 0.65 |
| ampc | 62 | 2964 | 0.520 | 0.144 | 0.005 | 0.018 | 0.053 | 1.65 | 0.65 | 0.81 |
| bace1 | 485 | 18706 | 0.636 | 0.208 | 0.019 | 0.053 | 0.127 | 1.65 | 2.15 | 2.02 |
| comt | 86 | 4012 | 0.527 | 0.122 | 0.000 | 0.000 | 0.006 | 0.00 | 0.00 | 0.00 |
| cp2c9 | 183 | 7757 | 0.599 | 0.221 | 0.129 | 0.110 | 0.166 | 4.95 | 2.96 | 2.41 |
| def | 161 | 5899 | 0.485 | 0.160 | 0.111 | 0.071 | 0.111 | 1.90 | 1.99 | 1.56 |
| fabp4 | 57 | 2911 | 0.852 | 0.520 | 0.611 | 0.486 | 0.536 | 26.42 | 10.57 | 6.32 |
| fgfr1 | 242 | 736 | 0.591 | 0.186 | 0.582 | 0.488 | 0.436 | 1.74 | 1.27 | 1.25 |
| fkb1a | 273 | 6105 | 0.709 | 0.251 | 0.040 | 0.114 | 0.192 | 1.10 | 2.42 | 2.68 |
| gria2 | 297 | 12356 | 0.658 | 0.217 | 0.044 | 0.041 | 0.115 | 1.69 | 2.02 | 1.89 |
| hs90a | 125 | 5067 | 0.364 | 0.079 | 0.000 | 0.000 | 0.006 | 0.00 | 0.00 | 0.24 |
| hxk4 | 127 | 4930 | 0.693 | 0.227 | 0.068 | 0.048 | 0.116 | 1.58 | 1.74 | 1.97 |
| jak2 | 153 | 6743 | 0.624 | 0.189 | 0.039 | 0.050 | 0.084 | 1.97 | 1.44 | 1.11 |
| mapk2 | 206 | 6450 | 0.717 | 0.276 | 0.157 | 0.186 | 0.221 | 7.34 | 3.70 | 2.72 |
| mk01 | 139 | 4767 | 0.803 | 0.369 | 0.203 | 0.206 | 0.372 | 5.11 | 7.64 | 5.76 |
| mk10 | 186 | 6900 | 0.711 | 0.239 | 0.048 | 0.069 | 0.136 | 2.69 | 2.04 | 2.10 |
| nos1 | 234 | 8307 | 0.591 | 0.184 | 0.001 | 0.021 | 0.087 | 0.00 | 1.20 | 1.45 |
| pa2ga | 127 | 5343 | 0.403 | 0.101 | 0.000 | 0.009 | 0.030 | 0.00 | 0.47 | 0.47 |
| pnph | 233 | 7249 | 0.575 | 0.161 | 0.032 | 0.049 | 0.071 | 1.73 | 0.95 | 0.90 |
| ppard | 288 | 13520 | 0.803 | 0.333 | 0.104 | 0.150 | 0.277 | 5.91 | 5.07 | 4.27 |
| ptn1 | 225 | 7658 | 0.751 | 0.308 | 0.098 | 0.144 | 0.279 | 3.58 | 5.26 | 4.18 |
| pur2 | 201 | 2926 | 0.719 | 0.238 | 0.014 | 0.051 | 0.172 | 0.50 | 1.70 | 2.29 |
| tysy | 311 | 7194 | 0.750 | 0.303 | 0.065 | 0.146 | 0.286 | 2.61 | 4.57 | 4.09 |

Supplementary Table 20. Wee1 target with 137 active compounds and 6371 compounds in total was generated 100 snapshots in 100ps (one snapshot every 1 ps) by OpenMM (a software for molecular dynamics) using the AlphaFold structure with a docked small molecule in the pocket. The results of virtual screening using idock as follows. The best value for EF1% is 31.00 from the 93ps snapshot. But we can’t split the good structures from other structures for virtual screening without the active and decoy label. In other words, we can’t split the good structures from other structures by some simple indication such as the best score or the average score.

| Target | AUC | logAUC | BEDROC(α=321.9) | BEDROC(α=80.5) | BEDROC(α=20.0) | EF1% | EF5% | EF10% |
| --- | --- | --- | --- | --- | --- | --- | --- | --- |
| AlphaFold | 0.762 | 0.240 | 0.009 | 0.013 | 0.083 | 0.74 | 0.44 | 1.61 |
| 1ps | 0.790 | 0.332 | 0.162 | 0.189 | 0.282 | 8.12 | 5.26 | 4.09 |
| 2ps | 0.735 | 0.278 | 0.035 | 0.097 | 0.212 | 2.21 | 4.39 | 3.21 |
| 3ps | 0.543 | 0.130 | 0.000 | 0.001 | 0.021 | 0.00 | 0.15 | 0.51 |
| 4ps | 0.655 | 0.186 | 0.000 | 0.009 | 0.047 | 0.00 | 0.44 | 0.80 |
| 5ps | 0.510 | 0.124 | 0.000 | 0.000 | 0.016 | 0.00 | 0.00 | 0.37 |
| 6ps | 0.808 | 0.314 | 0.019 | 0.077 | 0.234 | 0.74 | 4.53 | 3.65 |
| 7ps | 0.775 | 0.280 | 0.001 | 0.035 | 0.176 | 0.00 | 2.63 | 3.65 |
| 8ps | 0.685 | 0.209 | 0.000 | 0.004 | 0.075 | 0.00 | 0.88 | 1.61 |
| 9ps | 0.719 | 0.213 | 0.000 | 0.001 | 0.053 | 0.00 | 0.00 | 0.95 |
| 10ps | 0.750 | 0.237 | 0.000 | 0.011 | 0.092 | 0.00 | 1.17 | 1.75 |
| 11ps | 0.547 | 0.172 | 0.007 | 0.033 | 0.104 | 0.74 | 1.75 | 2.04 |
| 12ps | 0.736 | 0.236 | 0.041 | 0.022 | 0.087 | 0.74 | 0.44 | 1.68 |
| 13ps | 0.695 | 0.206 | 0.000 | 0.004 | 0.064 | 0.00 | 0.73 | 1.17 |
| 14ps | 0.674 | 0.201 | 0.015 | 0.015 | 0.064 | 0.74 | 0.44 | 1.24 |
| 15ps | 0.766 | 0.280 | 0.048 | 0.090 | 0.190 | 2.95 | 3.95 | 2.70 |
| 16ps | 0.743 | 0.254 | 0.001 | 0.029 | 0.138 | 0.00 | 2.34 | 2.34 |
| 17ps | 0.650 | 0.199 | 0.005 | 0.021 | 0.087 | 0.74 | 1.17 | 1.53 |
| 18ps | 0.674 | 0.198 | 0.002 | 0.008 | 0.059 | 0.74 | 0.29 | 1.17 |
| 19ps | 0.734 | 0.248 | 0.002 | 0.022 | 0.128 | 0.00 | 1.90 | 2.41 |
| 20ps | 0.766 | 0.251 | 0.004 | 0.017 | 0.103 | 0.74 | 1.17 | 2.12 |
| 21ps | 0.721 | 0.248 | 0.018 | 0.049 | 0.146 | 2.21 | 2.34 | 2.70 |
| 22ps | 0.675 | 0.202 | 0.001 | 0.013 | 0.076 | 0.00 | 1.17 | 1.39 |
| 23ps | 0.748 | 0.278 | 0.051 | 0.080 | 0.200 | 2.95 | 3.95 | 3.29 |
| 24ps | 0.704 | 0.237 | 0.014 | 0.023 | 0.123 | 0.74 | 1.61 | 2.56 |
| 25ps | 0.727 | 0.254 | 0.007 | 0.040 | 0.155 | 0.74 | 2.78 | 2.99 |
| 26ps | 0.738 | 0.253 | 0.005 | 0.035 | 0.136 | 0.74 | 2.19 | 2.34 |
| 27ps | 0.744 | 0.265 | 0.047 | 0.068 | 0.175 | 1.48 | 3.80 | 2.48 |
| 28ps | 0.685 | 0.222 | 0.033 | 0.022 | 0.111 | 0.74 | 1.90 | 2.41 |
| 29ps | 0.768 | 0.293 | 0.054 | 0.105 | 0.221 | 2.21 | 4.09 | 3.36 |
| 30ps | 0.809 | 0.320 | 0.077 | 0.119 | 0.247 | 4.43 | 4.97 | 3.72 |
| 31ps | 0.803 | 0.325 | 0.101 | 0.122 | 0.251 | 5.91 | 4.24 | 4.45 |
| 32ps | 0.649 | 0.195 | 0.005 | 0.031 | 0.085 | 0.74 | 1.32 | 1.39 |
| 33ps | 0.763 | 0.272 | 0.050 | 0.049 | 0.167 | 1.48 | 2.63 | 3.36 |
| 34ps | 0.604 | 0.176 | 0.030 | 0.031 | 0.064 | 0.74 | 1.02 | 0.88 |
| 35ps | 0.704 | 0.239 | 0.059 | 0.060 | 0.149 | 1.48 | 2.63 | 2.92 |
| 36ps | 0.611 | 0.172 | 0.013 | 0.016 | 0.053 | 0.74 | 0.58 | 0.80 |
| 37ps | 0.728 | 0.321 | 0.226 | 0.249 | 0.298 | 12.55 | 5.70 | 3.36 |
| 38ps | 0.678 | 0.248 | 0.103 | 0.122 | 0.180 | 5.17 | 3.51 | 2.34 |
| 39ps | 0.706 | 0.280 | 0.138 | 0.166 | 0.228 | 8.86 | 4.24 | 2.99 |
| 40ps | 0.645 | 0.230 | 0.126 | 0.121 | 0.164 | 5.91 | 2.92 | 2.12 |
| 41ps | 0.591 | 0.161 | 0.000 | 0.003 | 0.048 | 0.00 | 0.58 | 0.95 |
| 42ps | 0.744 | 0.275 | 0.021 | 0.062 | 0.201 | 1.48 | 3.51 | 3.72 |
| 43ps | 0.705 | 0.225 | 0.039 | 0.033 | 0.106 | 1.48 | 1.46 | 1.75 |
| 44ps | 0.682 | 0.288 | 0.195 | 0.212 | 0.269 | 10.33 | 5.70 | 3.36 |
| 45ps | 0.625 | 0.204 | 0.038 | 0.068 | 0.128 | 1.48 | 2.34 | 1.83 |
| 46ps | 0.643 | 0.184 | 0.001 | 0.009 | 0.059 | 0.00 | 0.88 | 1.31 |
| 47ps | 0.578 | 0.171 | 0.002 | 0.030 | 0.087 | 0.00 | 1.90 | 1.39 |
| 48ps | 0.671 | 0.243 | 0.108 | 0.111 | 0.174 | 5.17 | 3.66 | 2.34 |
| 49ps | 0.637 | 0.233 | 0.110 | 0.139 | 0.181 | 5.91 | 3.66 | 2.26 |
| 50ps | 0.697 | 0.241 | 0.011 | 0.057 | 0.162 | 1.48 | 3.22 | 2.77 |
| 51ps | 0.714 | 0.286 | 0.066 | 0.153 | 0.257 | 5.91 | 5.41 | 3.58 |
| 52ps | 0.713 | 0.261 | 0.073 | 0.112 | 0.193 | 2.95 | 3.95 | 2.63 |
| 53ps | 0.737 | 0.351 | 0.365 | 0.311 | 0.335 | 16.24 | 6.14 | 3.72 |
| 54ps | 0.765 | 0.405 | 0.337 | 0.386 | 0.437 | 20.67 | 8.92 | 5.04 |
| 55ps | 0.703 | 0.321 | 0.125 | 0.228 | 0.329 | 11.07 | 6.87 | 4.31 |
| 56ps | 0.752 | 0.396 | 0.486 | 0.408 | 0.391 | 24.36 | 7.46 | 4.16 |
| 57ps | 0.665 | 0.272 | 0.199 | 0.208 | 0.234 | 9.60 | 4.24 | 2.77 |
| 58ps | 0.764 | 0.315 | 0.093 | 0.159 | 0.280 | 5.17 | 6.14 | 3.94 |
| 59ps | 0.728 | 0.326 | 0.180 | 0.236 | 0.318 | 10.33 | 6.58 | 3.87 |
| 60ps | 0.639 | 0.187 | 0.002 | 0.022 | 0.080 | 0.00 | 1.75 | 1.46 |
| 61ps | 0.645 | 0.204 | 0.002 | 0.030 | 0.118 | 0.74 | 2.34 | 2.19 |
| 62ps | 0.564 | 0.144 | 0.000 | 0.006 | 0.039 | 0.00 | 0.58 | 0.88 |
| 63ps | 0.747 | 0.359 | 0.200 | 0.273 | 0.379 | 13.29 | 8.04 | 4.89 |
| 64ps | 0.819 | 0.331 | 0.080 | 0.134 | 0.262 | 3.69 | 5.12 | 3.94 |
| 65ps | 0.692 | 0.214 | 0.034 | 0.024 | 0.085 | 0.74 | 1.02 | 1.61 |
| 66ps | 0.814 | 0.390 | 0.257 | 0.295 | 0.382 | 13.29 | 7.46 | 4.75 |
| 67ps | 0.743 | 0.247 | 0.038 | 0.040 | 0.117 | 2.21 | 1.75 | 1.90 |
| 68ps | 0.761 | 0.300 | 0.031 | 0.108 | 0.246 | 2.21 | 4.83 | 4.09 |
| 69ps | 0.694 | 0.286 | 0.042 | 0.142 | 0.272 | 4.43 | 5.56 | 3.94 |
| 70ps | 0.743 | 0.258 | 0.008 | 0.033 | 0.158 | 0.74 | 2.78 | 3.21 |
| 71ps | 0.739 | 0.246 | 0.019 | 0.032 | 0.124 | 0.74 | 1.75 | 2.19 |
| 72ps | 0.696 | 0.216 | 0.004 | 0.017 | 0.084 | 0.74 | 1.02 | 1.53 |
| 73ps | 0.725 | 0.280 | 0.033 | 0.090 | 0.226 | 2.21 | 4.68 | 3.80 |
| 74ps | 0.749 | 0.299 | 0.059 | 0.107 | 0.252 | 1.48 | 5.70 | 3.87 |
| 75ps | 0.545 | 0.141 | 0.000 | 0.001 | 0.028 | 0.00 | 0.29 | 0.44 |
| 76ps | 0.724 | 0.290 | 0.090 | 0.140 | 0.256 | 5.91 | 5.41 | 3.80 |
| 77ps | 0.665 | 0.215 | 0.009 | 0.045 | 0.126 | 0.74 | 2.49 | 2.12 |
| 78ps | 0.620 | 0.185 | 0.001 | 0.015 | 0.086 | 0.00 | 1.46 | 1.90 |
| 79ps | 0.791 | 0.286 | 0.023 | 0.050 | 0.184 | 0.74 | 3.36 | 3.29 |
| 80ps | 0.628 | 0.176 | 0.006 | 0.018 | 0.054 | 0.74 | 0.73 | 0.95 |
| 81ps | 0.784 | 0.304 | 0.033 | 0.095 | 0.240 | 2.21 | 4.97 | 3.94 |
| 82ps | 0.751 | 0.314 | 0.104 | 0.171 | 0.287 | 5.17 | 6.00 | 3.80 |
| 83ps | 0.664 | 0.209 | 0.006 | 0.026 | 0.102 | 0.74 | 1.75 | 1.83 |
| 84ps | 0.800 | 0.399 | 0.305 | 0.346 | 0.400 | 19.93 | 7.90 | 4.67 |
| 85ps | 0.757 | 0.289 | 0.066 | 0.105 | 0.218 | 4.43 | 4.39 | 3.36 |
| 86ps | 0.809 | 0.379 | 0.172 | 0.254 | 0.377 | 10.33 | 7.75 | 4.89 |
| 87ps | 0.770 | 0.373 | 0.184 | 0.270 | 0.387 | 13.29 | 7.60 | 5.18 |
| 88ps | 0.504 | 0.126 | 0.000 | 0.006 | 0.032 | 0.00 | 0.73 | 0.51 |
| 89ps | 0.564 | 0.142 | 0.000 | 0.003 | 0.026 | 0.00 | 0.29 | 0.51 |
| 90ps | 0.643 | 0.200 | 0.043 | 0.060 | 0.106 | 2.21 | 1.90 | 1.61 |
| 91ps | 0.692 | 0.226 | 0.037 | 0.058 | 0.126 | 1.48 | 2.34 | 1.83 |
| 92ps | 0.732 | 0.378 | 0.427 | 0.384 | 0.391 | 18.45 | 7.90 | 4.09 |
| 93ps | 0.800 | 0.502 | 0.775 | 0.572 | 0.518 | 31.00 | 9.94 | 5.04 |
| 94ps | 0.630 | 0.213 | 0.206 | 0.115 | 0.111 | 6.64 | 1.61 | 1.24 |
| 95ps | 0.747 | 0.404 | 0.535 | 0.408 | 0.409 | 22.88 | 7.90 | 4.45 |
| 96ps | 0.676 | 0.291 | 0.134 | 0.188 | 0.281 | 8.86 | 5.70 | 3.94 |
| 97ps | 0.698 | 0.265 | 0.071 | 0.125 | 0.210 | 6.64 | 4.09 | 3.07 |
| 98ps | 0.666 | 0.288 | 0.335 | 0.236 | 0.261 | 11.07 | 5.41 | 2.99 |
| 99ps | 0.692 | 0.258 | 0.123 | 0.128 | 0.201 | 4.43 | 4.24 | 2.85 |
| 100ps | 0.679 | 0.241 | 0.019 | 0.073 | 0.183 | 1.48 | 4.09 | 2.92 |

Supplementary Figure 1. The Flow chart of multi-stage virtual screening without consensus scoring.

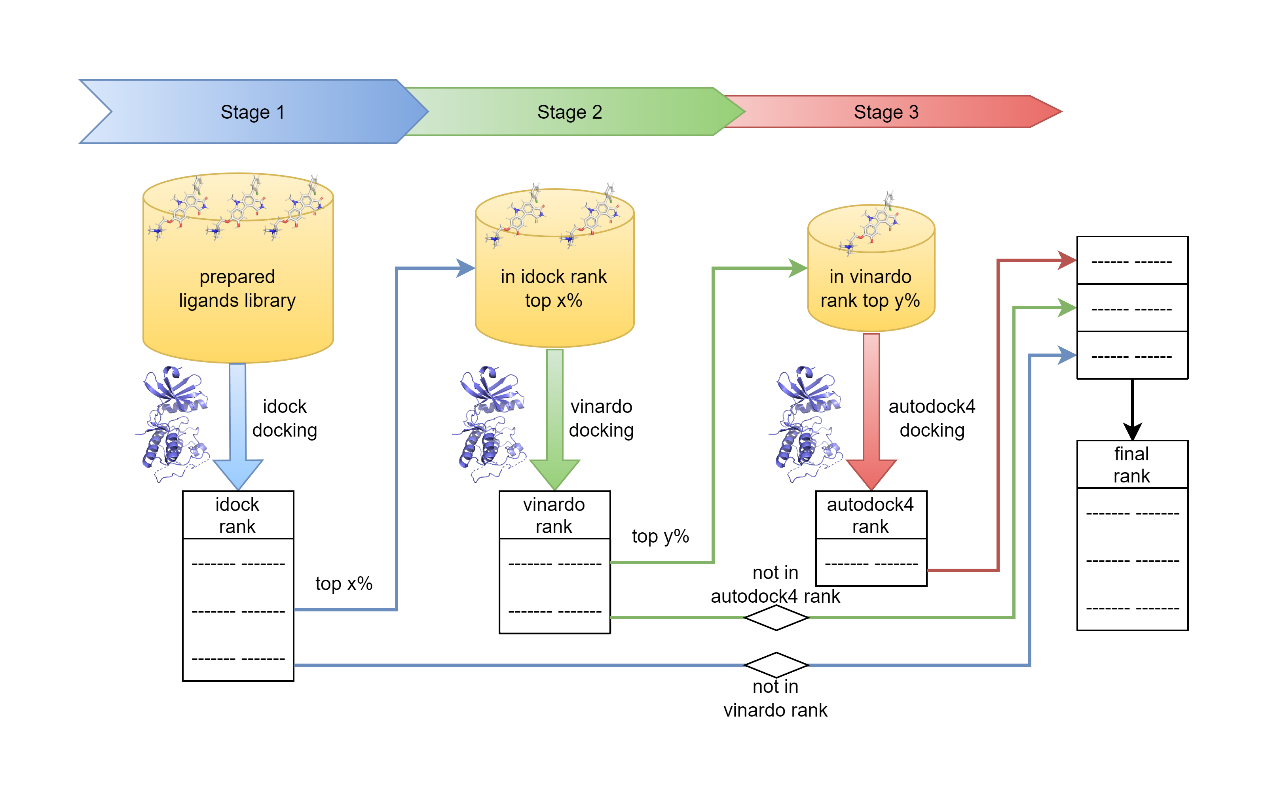

Supplementary Method

The input sequence used by ColabFold is as follows. The ':' symbol in the sequence is used to split two amino acid sequences:

For HIVPR, the sequence is selected according to the protein structure of PDB ID 1xl2, and the following data is used as the input sequence:

PQITLWQRPLVTIKIGGQLKEALLDTGADDTVLEEMSLPGRWKPKMIGGIGGFIKVRQYDQILIEICGHKAIGTVLVGPTPVNIIGRNLLTQIGCTLNF:PQITLWQRPLVTIKIGGQLKEALLDTGADDTVLEEMSLPGRWKPKMIGGIGGFIKVRQYDQILIEICGHKAIGTVLVGPTPVNIIGRNLLTQIGCTLNF

For HIVRT, the sequence is selected according to the protein structure of PDB ID 3lan, and the following data is used as the input sequence:

PISPIETVPVKLKPGMDGPKVKQWPLTEEKIKALVEICTEMEKEGKISKIGPENPYNTPVFAIKKKDSTKWRKLVDFRELNKRTQDFWEVQLGIPHPAGLKKKKSVTVLDVGDAYFSVPLDEDFRKYTAFTIPSINNETPGIRYQYNVLPQGWKGSPAIFQSSMTKILEPFRKQNPDIVIYQYMDDLYVGSDLEIGQHRTKIEELRQHLLRWGLTTPDKKHQKEPPFLWMGYELHPDKWTVQPIVLPEKDSWTVNDIQKLVGKLNWASQIYPGIKVRQLCKLLRGTKALTEVIPLTEEAELELAENREILKEPVHGVYYDPSKDLIAEIQKQGQGQWTYQIYQEPFKNLKTGKYARMRGAHTNDVKQLTEAVQKITTESIVIWGKTPKFKLPIQKETWETWWTEYWQATWIPEWEFVNTPPLVKLWYQLEKEPIVGAETFYVDGAANRETKLGKAGYVTNRGRQKVVTLTDTTNQKTELQAIYLALQDSGLEVNIVTDSQYALGIIQAQPDQSESELVNQIIEQLIKKEKVYLAWVPAHKGIGGNEQVDKLVS:IETVPVKLKPGMDGPKVKQWPLTEEKIKALVEICTEMEKEGKISKIGPENPYNTPVFAIKKKDSTKWRKLVDFRELNKRTQDFWEVQLGIPHPAGLKKKKSVTVLDVGDAYFSVPLDEDFRKYTAFTIPSINNETPGIRYQYNVLPQGWKGSPAIFQSSMTKILEPFRKQNPDIVIYQYMDDLYVGSDLEIGQHRTKIEELRQHLLRWGLTTPDKKHQKEPPFLWMGYELHPDKWTVQPIVLPEKDSWTVNDIQKLVGKLNWASQIYPGIKVRQLCKLLRGTKALTEVIPLTEEAELELAENREILKEPVHGVYYDPSKDLIAEIQKQGQGQWTYQIYQEPFKNLKTGKYARMRGAHTNDVKQLTEAVQKITTESIVIWGKTPKFKLPIQKETWETWWTEYWQATWIPEWEFVNTPPLVKLWYQL

The detailed configuration of ColabFold is as follows:

{

"num_queries": 1,

"use_templates": true,

"use_amber": true,

"msa_mode": "MMseqs2 (UniRef+Environmental)",

"model_type": "AlphaFold2-multimer-v2",

"num_models": 5,

"num_recycles": 3,

"num_ensemble": 1,

"model_order": [1,2,3,4,5],

"keep_existing_results": false,

"rank_by": "multimer",

"max_msa": null,

"pair_mode": "unpaired+paired",

"host_url": "https://api.colabfold.com",

"stop_at_score": 100.0,

"stop_at_score_below": 0,

"recompile_padding": 1.0,

"recompile_all_models": false,

"commit": "26de12d3afb5f85d49d0c7db1b9371f034388395",

"is_training": false,

"version": "1.3.0"

}

Select the first ranked and relaxed structure, hivpr_23119_relaxed_rank_1_model_ 5.pdb, is used as the AlphaFold structure of HIVPR.

Select the first ranked and relaxed structure, hivrt_88c27_relaxed_rank_1_model_ 3.pdb, is used as the AlphaFold structure of HIVRT.
